# High satellite repeat turnover in great apes studied with short- and long-read technologies

**DOI:** 10.1101/470054

**Authors:** Monika Cechova, Robert S. Harris, Marta Tomaszkiewicz, Barbara Arbeithuber, Francesca Chiaromonte, Kateryna D. Makova

## Abstract

Satellite repeats are a structural component of centromeres and telomeres, and in some instances their divergence is known to drive speciation. Due to their highly repetitive nature, satellite sequences have been understudied and underrepresented in genome assemblies. To investigate their turnover in great apes, we studied satellite repeats of unit sizes up to 50 bp in human, chimpanzee, bonobo, gorilla, and Sumatran and Bornean orangutans, using unassembled short and long sequencing reads. The density of satellite repeats, as identified from accurate short reads (Illumina), varied greatly among great ape genomes. These were dominated by a handful of abundant repeated motifs, frequently shared among species, which formed two groups: (1) the (AATGG)_n_ repeat (critical for heat shock response) and its derivatives; and (2) subtelomeric 32-mers involved in telomeric metabolism. Using the densities of abundant repeats, individuals could be classified into species. However clustering did not reproduce the accepted species phylogeny, suggesting rapid repeat evolution. Several abundant repeats were enriched in males vs. females; using Y chromosome assemblies or FIuorescent In Situ Hybridization, we validated their location on the Y. Finally, applying a novel computational tool, we identified many satellite repeats completely embedded within long Oxford Nanopore and Pacific Biosciences reads. Such repeats were up to 59 kb in length and consisted of perfect repeats interspersed with other similar sequences. Our results based on sequencing reads generated with three different technologies provide the first detailed characterization of great ape satellite repeats, and open new avenues for exploring their functions.

## Introduction

Heterochromatin is the gene-poor and highly compacted portion of the genome. It is typically dominated by *satellite repeats* – long arrays of tandemly repeated non-coding DNA (Sueoka 1961; Kit 1961) that consist of smaller units organized into higher-order repeat structures. Heterochromatin is abundant, for instance, at telomeres and centromeres of human chromosomes (Sujiwattanarat et al. 2015). A similar enrichment of heterochromatin with satellite repeats is widespread at the centromeres of many animal and plant genomes (Melters et al. 2013).

While labeled as “junk DNA” in the past, heterochromatin was later found to fulfill important functions in the genome (Walker 1971; Yunis and Yasmineh 1971; Ferree and Barbash 2009). Heterochromatin satellite repeat expansions have been associated with changes in gene expression and methylation (Brahmachary et al. 2014; Quilez et al. 2016). It has also been proposed that heterochromatin aids in maintaining cellular identity by repressing genes that are not specific to a particular cell lineage (reviewed in (Becker et al. 2016)). For instance, the heterochromatin-associated histone mark H3K9me3 blocks reprogramming to pluripotency (Soufi et al. 2012). Additionally, heterochromatin loss is part of the normal aging process (Zhang et al. 2015). Similarly, heterochromatin changes during stress. For instance, gene silencing at heterochromatin is less effective at high temperatures in yeast (Ayoub et al. 1999; Gowen and Gay 1933); heterochromatin-induced gene inactivation (known as “position-effect variegation”) is sensitive to temperature in both yeast and *Drosophila* (Allshire et al. 1995; Gowen and Gay 1933; Spofford 1976); and the latter effect was shown to be variable within a natural *Drosophila* population (Kelsey and Clark 2017). Moreover, in the rods of the retinas of nocturnal mammals, heterochromatin is localized towards the central regions of the nucleus and acts as a lens to channel light (Solovei et al. 2009).

Despite a growing interest in understanding these important functions of heterochromatin, satellite repeats are frequently underrepresented in genomic studies – due to the difficulties in sequencing and assembling these highly similar sequences (Chaisson et al. 2015). Thus, they remain understudied. The lack of information about satellite repeats is particularly alarming given their high abundance, e.g., alpha satellites were estimated to constitute approximately 3% of the human genome (Manuelidis 1978; Hayden et al. 2013). Relatedly, satellite repeats are likely plentiful in yet unassembled gaps in the human genome (Miga et al. 2014; Stephens and Iyer 2018). One of the largest uncharacterized gaps in the human genome is located in the Male-Specific region of the Y chromosome (MSY), which contains six types of satellite repeat sequences (DYZ1, DYZ2, DYZ3, DYZ17, DYZ18, and DYZ19) (Skaletsky et al. 2003).

Heterochromatin exhibits remarkable *interspecific variability* in size and structure. Such variability can be frequently observed even between closely related species. For instance, on the long arm of the Y chromosome, heterochromatin is the major component in human and gorilla, but is virtually absent in chimpanzee (Gläser et al. 1998) – notwithstanding the fact that human, gorilla, and chimpanzee diverged less than 8 million years (MY) ago (Glazko and Nei 2003). As another example, whereas 20% of the genome of *Drosophila melanogaster* is composed of satellite DNA, this percentage is as low as 0.5% for *D. erecta* and as high as 50% for *D. virilis* (Gall et al. 1971; Lohe and Brutlag 1987); the estimated divergence time between *D. erecta* and *D. melanogaster* is 13 MY, and is 63 MY between *D. virilis* and *D. melanogaster* (Tamura et al. 2004). The differences in satellite repeat abundance in nine *Drosophila* species were proposed to result predominantly from lineage-specific gains accumulated over the past 40 MY of evolution (Wei et al. 2018). Due to its rapid evolutionary turnover, heterochromatin can serve as a species barrier (Yunis and Yasmineh 1971). For instance, the female hybrids between *D. melanogaster* males and *D. simulans* females are not viable because, during cell division, they fail to properly separate the satellite 359-bp repeat on the X chromosome (Ferree and Barbash 2009; Rošić et al. 2014).

Profound *intraspecific* variability in heterochromatin has also been reported, including that among humans (Altemose et al. 2014; Miga et al. 2014). For instance, the length of the DYZ1 satellite repeat varies considerably among major Y chromosome haplogroups; DYZ1 is longer in Y chromosomes belonging to the predominantly Asian O haplogroup than in those belonging to the predominantly African E haplogroup (Altemose et al. 2014). The centromeric array of the X chromosome was shown to vary in length among different human populations by as much as an order of magnitude (0.5-5 Mb) (Miga et al. 2014). Some human neocentromeres were found to harbor only very short (as short as 15-kb) heterochromatin domains leading to a defect in sister chromatid cohesion (Alonso et al. 2010).

In addition to satellite repeats with relatively long repeat units (e.g., alpha satellites with repeat unit of ∼171 bp), three classes of satellite repeats with unit sizes ≤50 bp are of a particular interest due to their abundance or function in great apes. These include (AATGG)_n_ satellite, telomeric satellite (TTAGGG)_n_, and AT-rich 32-unit Subterminal satellites (StSats). The (AATGG)_n_ repeat is the source of Human Satellites 2 and 3 (HSat2 and HSat3) (Altemose et al. 2014). On chromosome 9, it also encodes a long noncoding RNA that is critical for the heat shock response in human cells (Goenka et al. 2016). Previous studies investigated the variability, abundance, and length distribution of the (AATGG)_n_ repeat in the human genome (Tagarro et al. 1994; Skaletsky et al. 2003; Altemose et al. 2014; Subramanian et al. 2003). This repeat was also identified in orangutan, chicken, maize, sea urchin, and *Daphnia* (Grady et al. 1992; Flynn et al. 2017), however its variation in great ape species was never studied. The telomeric (TTAGGG)_n_ satellite functions to maintain genome stability; telomere loss is correlated with cell division and aging (Lanza et al. 2000; Rizvi et al. 2014). Subterminal satellites (StSats), present in the genomes of chimpanzee, bonobo, and gorilla (Royle et al. 1994), localize proximal to telomeres (Ventura et al. 2012; Royle et al. 1994; Koga et al. 2011) and were proposed to play a role in telomere metabolism (Novo et al. 2013) and meiotic telomere clustering important for homolog recognition and pairing (in a process similar to that identified in plants (Calderón et al. 2014; Bass et al. 2000)).

In this study, we characterize turnover of satellites with repeat units ≤50 bp among six great ape species – human, chimpanzee, bonobo, gorilla, Bornean orangutan, and Sumatran orangutan – which diverged less than ∼14 MY ago (Goodman et al. 2005). We focus on repeats that constitute portions of long arrays of satellite DNA and use them as a proxy for heterochromatin (Wei et al. 2014). This approximation is needed because of challenges in the direct identification of heterochromatin due to its transient nature in various cells of individuals throughout their lifetime. In this manuscript, we, first, identify satellite repeats in short sequencing reads generated with the low-error-rate Illumina technology, and investigate their inter- and intraspecific variation. We pinpoint repeats with higher incidence in males than females and, for some of these repeats, confirm location on the Y chromosome using existing Y assemblies or fluorescent in situ hybridization (FISH). Next, we use the repeated motifs identified from low-error-rate short reads as queries to decipher the lengths and densities of ape satellite repeats from error-prone long reads (both Pacific Biosciences, or PacBio, and Oxford Nanopore, or Nanopore). To the best of our knowledge, ours is the first study of inter- and intraspecific satellite repeat variability, repeat expansions and correlations, as well as of male-biased repeats, in great apes.

## Results

### Repeat identification in short reads

To study inter- and intraspecific variability of satellite repeats in great apes, we utilized 100-or 150-base-pair (bp) Illumina sequencing reads generated for 79 individuals (57 females and 22 males; Table S1) as a part of the Ape Diversity Project (ADP) (Prado-Martinez et al. 2013). These included chimpanzees (Nigeria-Cameroon, Eastern, Central, and Western chimpanzees), bonobos, gorillas (Eastern lowland, Cross river, and Western lowland gorillas), Sumatran orangutans, and Bornean orangutans (Table S1). Additionally, in order to match the library preparation protocol that was used for these great ape data, we used sequencing reads for 9 human males from diverse populations generated as part of the Human Genome Diversity Project (HGDP) (Meyer et al. 2012; Cann et al. 2002; Rosenberg et al. 2002). After filtering (see Methods), in this set of 79 + 9 = 88 individuals, the median number of reads per individual was 190,722,592 (Table S1).

Sequencing reads are expected to present a more complete picture of satellite repeat distributions than the existing reference genome assemblies (Lower et al. 2018). To annotate repeats in sequencing reads, we used Tandem Repeats Finder, TRF (Benson 1999) (when available, 150-bp reads were trimmed to 100 bps for consistency) and focused on repeats with a repeated unit of ≤50 bp (so that at least two units could fit within a 100-bp read). This approach does not allow detection of satellites with longer repeated units, such as centromeric alpha satellites, for which even a single repeated unit would not fit within a short sequencing read, however is geared towards accurate identification of satellite repeats with shorter repeated units. Additionally, in order to study long satellite arrays likely to be present in the heterochromatin, we only retained sequencing reads in which repeated arrays covered at least 75% of the read length (i.e. ≥75 bp, see Methods). This effectively removed most microsatellites from our data set. As a result, we identified 5,494 distinct repeated motifs (later called *satellite repeated motifs*, or *repeated motifs*) across the studied species and verified that they were not artifacts of read length or software choice (Supplementary Note 1).

### Inter- and intraspecific variability

#### Repeat density varies among great ape species

We compared the overall satellite repeat density (computed cumulating occurrences for all types of repeated motifs) among the studied ape species and subspecies (Fig. 1A). For each individual, *satellite repeat density* (later called *repeat density*) was computed as the total number of kilobases annotated in satellite repeats per million bases of sequencing reads (kb/Mb). First, we verified that technical replicates – different Illumina lanes/runs for the same individual – had highly correlated repeat densities (Fig. S1). Second, we verified that repeat density and sequencing depth were not correlated with each other (Fig. S2). Illumina PCR+ libraries were generated for ADP (Prado-Martinez et al. 2013) and HGDP (Cann et al. 2002; Rosenberg et al. 2002; Meyer et al. 2012); while the types of repeated motifs identified were likely unaffected by the amplification step during library preparation, their densities might have been (Supplementary Note 2) and thus the precise repeat densities we report here might differ from the actual densities in the studied genomes. However, biases due to PCR amplification should be limited (see next paragraph for an analysis of human PCR-libraries suggesting minimal bias). Moreover, because all samples were processed with the same library preparation protocol, any existing biases should be concordant and not affect comparisons of numbers among and within species (Fig. 1A). We observed the highest average repeat densities (across individuals) in Western and Eastern lowland gorillas (103 and 74.0 kb/Mb, respectively), and the lowest in human (11.9 kb/Mb) and Sumatran orangutan (22.6 kb/Mb).

**Figure 1.**
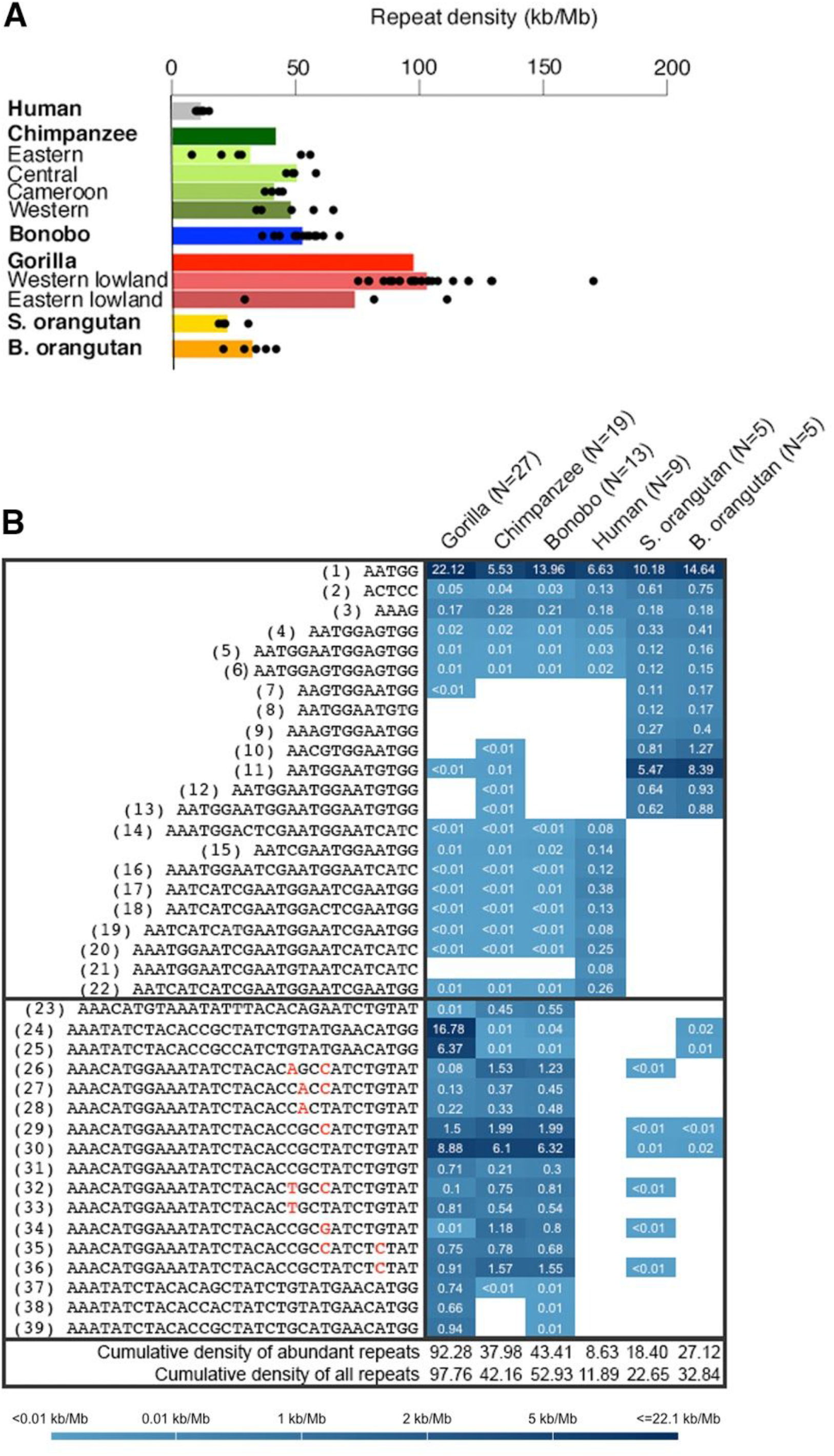
Densities and similarity among satellite repeats in great apes. **(A) Intra-and interspecific variation in overall repeat density.** Repeat densities are plotted for each species and subspecies. Each dot represents a single individual, and bars are mean values. For species comprising subspecies, a species-level average is also represented as a bar. Human (N=9, black), bonobo (N=13, blue), chimpanzee (N=19, green), gorilla (N=27, red), S. orangutan is Sumatran orangutan (N=5, yellow), and B. orangutan is Bornean orangutan (N=5, orange). The cross river gorilla has sample size of 1 and is only included in the species-level analysis. **(B) Heatmap of average repeat densities (across individuals) for the 39 abundant repeats in each of the six species.** Color coding from dark to light blue represents high to low values. Repeats present at less than 100 loci per 20 million reads are considered absent (white cells). (AATGG)_n_-derived and 32-mer-derived repeated motifs are separated by a horizontal line. Cumulative densities of abundant repeats and of all repeats are calculated as averages across all individuals.

#### Great ape genomes harbor only a handful of abundant repeated motifs, many of which are shared among species and are phylogenetically related

We next investigated whether great ape genomes possess a few highly abundant repeated motifs, or many different repeated motifs present at relatively low abundance. We ranked motifs by abundance and found that the six great ape species we considered (subspecies were combined for this analysis) contain only a small number of abundant repeated motifs: usually ≤12 in each of the species (Fig. S3). There were a total of 39 unique motifs in the set of 12 motifs × 6 species = 72 repeated motifs with density ranking 12 or higher in the six species analyzed (Fig. S3, Table S2). These 39 repeated motifs had overall average densities (across individuals) of 8.63 kb/Mb, 38.0 kb/Mb, 43.4 kb/Mb, 92.3 kb/Mb, 18.4 kb/Mb, and 27.1 kb/Mb in the six species (Fig. 1B), and represent approximately 73%, 90%, 82%, 94%, 81%, and 83% (i.e. very large portions) of the total satellite repeat density we found in the human, chimpanzee, bonobo, gorilla, Sumatran orangutan, and Bornean orangutan genomes, respectively. Notably, when we compared densities of these 39 repeats between nine humans sequenced with the PCR+ protocol used also for non-human apes throughout our study and nine other humans sequenced with a PCR-protocol (Fig. S4), we observed minimal differences beyond expected interindividual variation, suggesting only small effects of PCR amplification on our repeat density estimates.

As a control, we searched for the telomeric (TTAGGG)_n_ repeat which we expect to be present in our data set; in our data this repeat has ranks 42, 112, 144, 321, 43, and 38 in the genomes of human, chimpanzee, bonobo, gorilla, Sumatran orangutan, and Bornean orangutan genomes, respectively (Fig. S3). Thus, it is not one of the most abundant repeats in great apes. It constitutes less than 0.23%, 0.10%, 0.06%, 0.02%, 0.10% and 0.11% of the total satellite repeat density in each of these genomes, with repeat density ranging from 0.0227 to 0.0422 kb/Mb among species (Table S3).

The 39 abundant repeated motifs we identified had varying levels of sharing among species (Fig. 1B). Six motifs were present in all six species analyzed. The (AATGG)_n_ repeat, shared by all six species, was the most abundant repeat in humans (with an average density of 6.63 kb/Mb) as well as in gorilla, bonobo, Sumatran orangutan, and Bornean orangutan (with average densities of 22.1 kb/Mb, 14.6 kb/Mb, 10.2 kb/Mb and 14.6 kb/Mb, respectively), and the second most abundant repeat in chimpanzee (with an average density of 5.53 kb/Mb). The next most abundant repeated motifs in human and orangutans were phylogenetically related to the (AATGG)_n_ (Figs. 1B, S5, Table S2). Their overall average densities (excluding (AATGG)_n_ itself) were 1.62 kb/Mb, 9.22 kb/Mb, and 13.7 kb/Mb in the genomes of human, Sumatran orangutan, and Bornean orangutan, respectively (Fig. 1B). In addition to (AATGG)_n_ and repeated motifs related to it, we identified highly similar Subterminal Satellite (StSat) 32-mers (Royle et al. 1994; Koga et al. 2011; Ventura et al. 2012) and a 31-mer related to them (all differing by 1-2 bases; Figs. 1B, S5). These repeats were abundant in the genomes of chimpanzee, bonobo, and gorilla with overall average densities of 15.8 kb/Mb, 15.8 kb/Mb, and 39.6 kb/Mb, respectively. In fact, one of these 32-mers was the motif with the highest repeat density in chimpanzee (6.10 kb/Mb). 32-mers were absent from the human genomes analyzed, and were very sparse in the orangutan genomes (Fig. 1B). We found no relationship between the degree to which a repeated motif was shared across the six species and its repeat density (Fig. S6). In conclusion, the overall satellite repeat content in great ape genomes appears to be driven by only a few highly abundant repeated motifs, many of which are shared among species and are phylogenetically related to each other (Fig. S5).

#### The majority of less abundant repeated motifs are species-specific

We subsequently analyzed the 5,455 repeated motifs constituted by the initial set minus the 39 abundant repeats discussed in the previous section, and found substantial differences among great ape species when profiling their presence/absence (Fig. S7B). Despite the relatively recent divergence of the species considered (Goodman et al. 2005), as many as 3,170 of the 5,455 distinct repeated motifs were species-specific. Among them, 2,312 were gorilla-specific, while only 262 were human-specific. As expected, the chimpanzee and bonobo sister species shared many repeated motifs (a total of 947, representing 75% and 78% of all repeats identified in each species, respectively), and so did the Sumatran and Bornean orangutan sister species (a total of 217, representing 99% and 97% of all repeats identified in each species, respectively). Interestingly, we found a positive relationship between the number of species-specific repeated motifs and mean repeat density in a species (Fig. S8; human is an outlier in this analysis). These results did not change qualitatively when we considered the same number of individuals per species (Figs. S7C-H and S8B).

#### Substantial differences exist among individuals, in repeat presence/absence as well as density

The majority of the 39 abundant motifs were present in all individuals of a given species (Fig. S7A) but exhibited substantial variability in repeat density among them (Table S4, Fig. S9). For instance, the average fold difference for the (AATGG)_n_ repeat among two unrelated human males in our study was 1.23. Other motifs, especially those of lower abundance, although identified in a species, were only present in a subset of individuals (Fig. S7B).

#### Relatedness of the studied species based on satellite repeat data

##### (a) Individuals can be classified into species based on 14 unique, most abundant repeated motifs

We investigated whether the densities of the 39 abundant repeats found across great ape genomes (Table S2) could separate individuals into species. We started with an exploratory Principal Component Analysis (PCA, Fig. 2A) of their densities. In the space of the first three components (which explain 98% of the variance; Table S5), individuals belonging to different species formed fairly well-separated groups. Next, we attempted to directly classify individuals into species. We found that using the densities of just 14 most abundant repeats (from the set of 39) already produced excellent classification performance; leave-one-out cross-validation resulted in accuracy of ∼96% for a Linear Discriminant Analysis (LDA) classifier with uniform priors (Fig. 2B), and ∼93% for a Random Forest classifier (Table S5). Using up to 20 most abundant repeats, the accuracy of LDA classification was as high as 97% (Fig. 2B).

**Figure 2.**
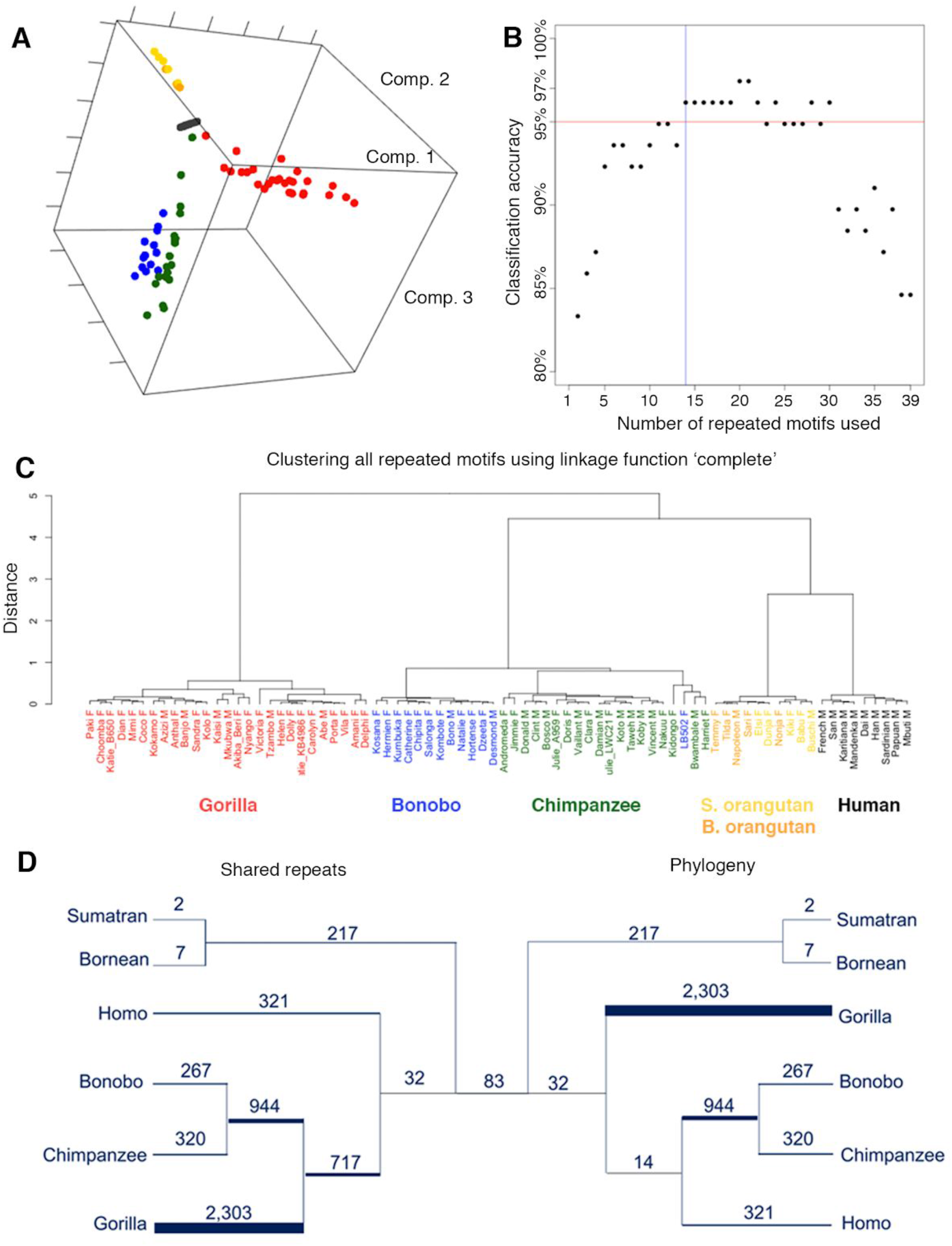
Relatedness of 88 analyzed individuals belonging to six great ape species. **(A) Principal Component Analysis.** Individuals are plotted as circles in the space of the first three principal components extracted from the densities of the 39 abundant repeats, which explain 98% of the variance. Colors correspond to the six species: human (black), bonobo (blue), chimpanzee (green), gorilla (red), Sumatran orangutan (orange), and Bornean orangutan (gold). **(B) Cross-validation accuracy of a Linear Discriminant Analysis classification of individuals into species, as a function of the number of abundant repeats used.** The 39 abundant repeats were progressively added to the LDA classifier in decreasing order of abundance. Using the motif indexes from Fig. 1B, this order was: 1, 30, 17, 24, 11, 22, 29, 10, 20, 36, 25, 12, 3, 26, 13, 15, 34, 32, 39, 2, 35, 4, 18, 33, 9, 16, 23, 19, 37, 5, 8, 21, 27, 28, 31, 6, 7, 14, and 38. The accuracy increases as the first 20 repeated motifs are added, and then decreases due to overfitting. The blue vertical line marks the first 14 repeated motifs, and the red horizontal one the 95% accuracy level. **(C) Hierarchical clustering of individuals.** This clustering employs the densities of all 5,494 repeated motifs, Spearman correlations, and complete linkage. **(D) Species topology based on repeats presence/absence.** A schematic figure showing repeated motifs unique to a species (terminal branches) and those that are shared among the species descending from internal branches. On the left, the tree is built based on the presence/absence of repeated motifs, iteratively joining species sharing the most repeated motifs. On the right, the tree is built according to the accepted species phylogeny (Goodman et al. 2005) and the number of shared repeated motifs is indicated. The branch widths are proportional to the number of repeated motifs (branch lengths are uninformative). 83 repeated motifs (the number shown in the middle) were shared among all six studied species.

##### (b) Hierarchical clustering based on repeat densities usually does not reproduce the accepted species phylogeny

Based on the results in (a), the question naturally arises of whether a hierarchical clustering of individuals would reproduce the accepted species phylogeny (Goodman et al. 2005). To address this, we computed distances between individuals based on Pearson and, separately, Spearman correlation coefficients – the latter being a more robust measure of similarity. Specifically, each individual was represented by a vector of repeat densities (for Pearson), or of ranks of such densities (for Spearman) and, for each pair of individuals, the correlation was calculated between their two vectors. For both these correlation types, we performed two separate analyses: (1) using all 5,494 unique repeated motifs; and (2) using the 39 abundant repeats. Also, two different linkage functions were used to implement the hierarchical clustering in each analysis and with each correlation type: “single” (which joins two clusters based on the minimal distance between individuals and thus mimics the maximum parsimony approach), and “complete” (which instead uses the maximal distance and in general tends to form compact clusters; see Methods). In these analyses species always formed well-separated clusters (Fig. S10). Unexpectedly, in most of the analyses – using Pearson correlations and complete linkage, as well as using Spearman correlations and complete linkage – humans clustered with orangutans in both (1) and (2) (Figs. 2C, S9A,C,E-G), contradicting the accepted species phylogeny. Similarly, using Spearman correlation and single linkage, orangutans, but not humans, clustered with gorilla and chimpanzee/bonobo, also contradicting the accepted species phylogeny (Fig. S10H). The higher-level agglomeration only reproduced the accepted species phylogeny in scenarios where Pearson correlations and single linkage were used (on all, or on only the 39 abundant repeats; Figs. S10B,D).

##### (c) Phylogeny based on presence/absence of repeated motifs does not reproduce the accepted species phylogeny

We observed a similar pattern estimating a phylogeny based solely on the number of shared repeated motifs (in terms of their presence/absence). Chimpanzee, bonobo, and gorilla (sharing 717 repeated motifs) formed a cluster that did not include human (Fig. 2D, left), departing from the accepted great apes phylogeny (Fig. 2D, right). Note that many highly abundant repeats were also shared among chimpanzee, bonobo, and gorilla (Fig. 1B). In contrast, human, chimpanzee, and bonobo, while having a more recent common ancestor, shared as few as 14 repeated motifs (Fig. 2D). The pattern was the same even after the exclusion of the StSat repeated motifs (Fig. S11). Taken together, both distances in repeat densities (Fig. 2C) and configurations of shared (vs. not shared) repeats (Fig. 2D) across species, show a distortion of the signals as compared with the accepted species phylogeny. This suggests an especially rapid evolution of satellite repeats among great apes.

#### The densities of the 39 abundant repeats display high correlations, particularly for similar repeated motifs

Next, we computed Spearman correlation coefficients between pairs of repeats among the 39 abundant ones found in great apes genomes. Here, each repeat is represented by a vector of ranks for its densities across individuals (in each species). The significance of these correlations was tested against a chance background scenario simulated by random reshuffling of repeated motif labels (Figs. 3 and S12; see Methods). Most correlation coefficients were positive and rather large. Furthermore, we found that blocks with strong positive correlations tended to comprise phylogenetically related repeated motifs (Figs. 3 and S5). Negative and moderately large coefficients (r<-0.5) were also observed 11). In general, negative correlations were rare and mostly associated with the (AAAG)_n_ repeat (Fig. S12).

**Figure 3.**
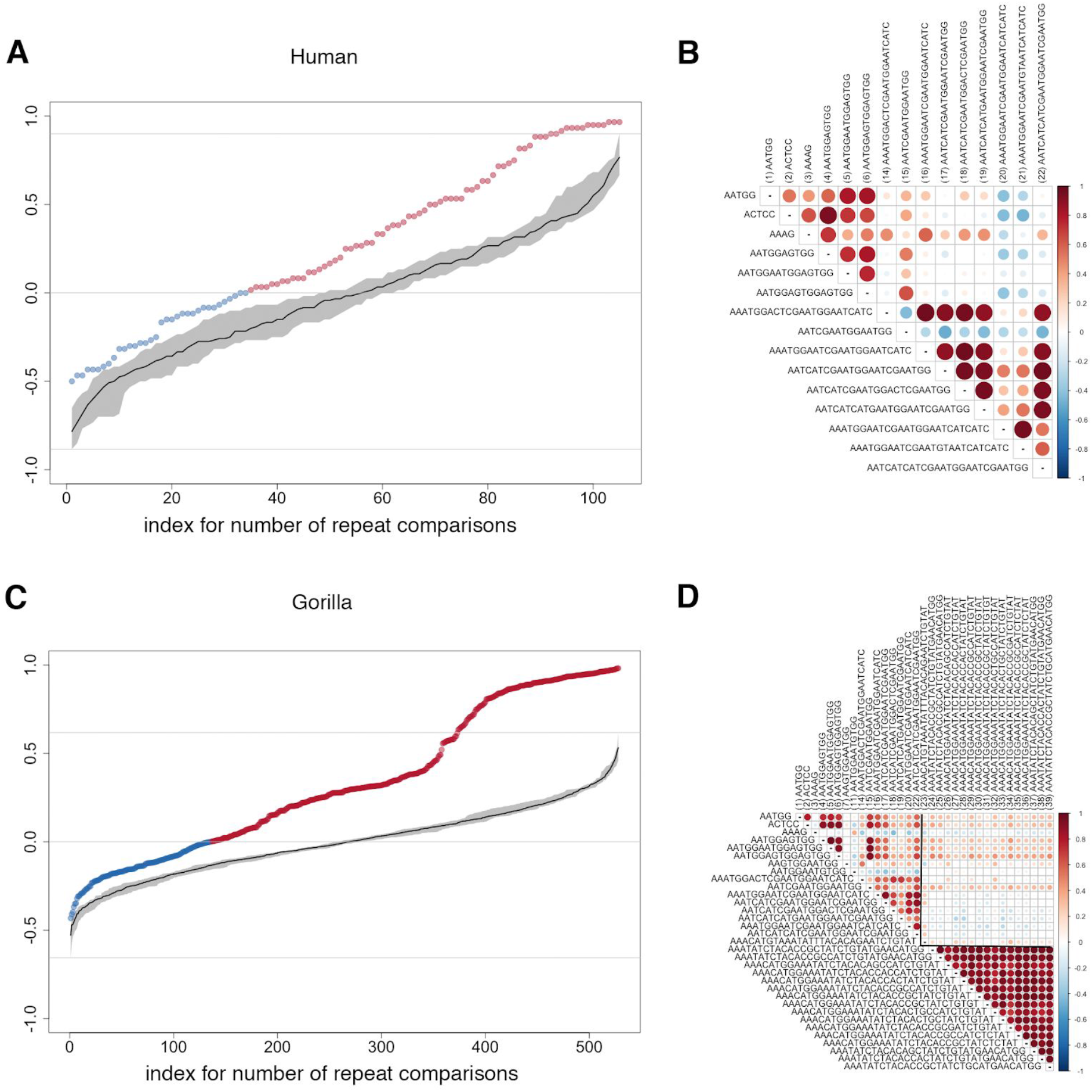
Spearman correlations for the densities of the 39 abundant repeats in human and gorilla. Colored dots in the upper (**A**; Human, n=105 comparisons) and lower (**C**; Gorilla, n=528 comparisons) left panels show observed correlations between pairs of repeats plotted in non-decreasing order, in red when positive and in blue when negative. Chance background correlations, again in non-decreasing order, are plotted in black with variation bands in gray (see Methods). The heatmaps in the upper (**B**; Human) and lower (**D**; Gorilla) right panels show the correlations corresponding to each repeat pair, with various intensities of red (positive) and blue (negative). The size of the circles is also proportional to that of the correlation. Fig. S9 provides the same information for the other species. Only repeats present in the relevant species are shown.

The correlations between abundant repeats densities differed across great ape species. For example, in human, we observed many more positive correlations than expected by chance, but also a substantial number of negative correlations (Figs. 3A-B). In contrast, gorilla had a sizable subset of repeats with very high and significant positive correlations (coefficients >0.8), but very few negative correlations (Figs. 3C-D). Also interestingly, the two orangutan species displayed different patterns. In Sumatran orangutan, just as in human, we observed more positive than negative correlations – but none of the coefficients were significant. In Bornean orangutan, positive correlations were significant and negative correlations were not.

### Male-biased repeats

#### Male-biased repeats are among the most abundant

We next focused on identifying repeats potentially located on great apes Y chromosomes, based on the expectation that they should be significantly more frequent in males than in females, i.e. male-biased. We considered all chimpanzee, bonobo and gorilla individuals, as well as ten orangutan individuals (combining five Sumatran and five Bornean). In addition to the nine human males from HGDP (Meyer et al. 2012; Cann et al. 2002; Rosenberg et al. 2002), we also used three fathers and three mothers from human trios (Table S1, see Methods). We restricted attention to repeated motifs with density above 0.5 kb/Mb in any given species. For each such motif we calculated the average male-to-female density ratio across individuals and assessed significance of the difference in repeat density between males and females with a Mann-Whitney test (Table S6). Since our sample sizes were relatively small, we used a high p-value cutoff (alpha=0.2) to compensate for lack of power – this, admittedly, increases the chances of false positives in our results. Our analysis resulted in a total of 18 significantly male-biased repeated motifs, which are candidates to be located on great apes Y chromosomes: one in human ((AATGG)_n_), five in chimpanzee, nine in bonobo, fourteen in gorilla, and one in orangutans ((ACTCC)_n_) (Table S6). Interestingly, all the significantly male-biased motifs were among the most abundant repeated motifs in the ape genomes (ranging between 1st and 14th in the species-specific ranks).

#### Male-biased 32-mers can be found on the gorilla and bonobo Y chromosomes

We further restricted attention to male-biased 32-mer StSats, which had higher incidence in males than females in chimpanzee, bonobo, and gorilla (Table S6), and searched for additional evidence that they indeed might be located on the Y chromosomes of these species. First, we screened the Y chromosome assemblies of chimpanzee (Hughes et al. 2010) and gorilla (Tomaszkiewicz et al. 2016) for occurrences of these male-biased StSats (see Methods; no bonobo Y chromosome assembly is currently available). We found them in the latter but not in the former. This could be explained by the fact that long PacBio reads, which are more likely to capture these StSats, were used to generate the gorilla’s Y assembly, and not the chimpanzee’s. However it is also possible that some of these StSats are indeed absent from the chimpanzee Y chromosome (see next paragraph).

Second, to experimentally assess whether male-biased StSats (Table S6) are present on the Y chromosomes of bonobo and chimpanzee, we performed FISH. We used two probes (see Methods); a degenerate probe containing the sequences of two male-biased StSats (Table S6), and a probe containing the flow-sorted bonobo Y chromosome. These probes were hybridized to metaphase spreads of bonobo and chimpanzee males. The StSat probe hybridized to (sub)telomeric locations of most chromosomes (Figs. 4A-B), suggesting an association with heterochromatin. Moreover, both probes hybridized to the bonobo Y chromosome, confirming Y localization (Fig. 4D) – consistent with our computational predictions (the p-values for bonobo male-to-female abundance differences were 0.03 and 0.05 for the two StSats included in the degenerate probe; Table S6). FISH could not confirm the presence of the same StSat probe on the chimpanzee Y chromosome (Fig. S13B) – again consistent with our computational analysis, which provided only weak evidence of male bias for the studied StSats in chimpanzee (p-values of 0.2 and 0.2 for the two StSats included in the degenerate probe; Table S6). In summary, we identified several male-biased repeats in the genomes of great ape species, and for a number of them we were able to validate their Y chromosome location either by examining Y assemblies or by FISH experiments.

**Figure 4.**
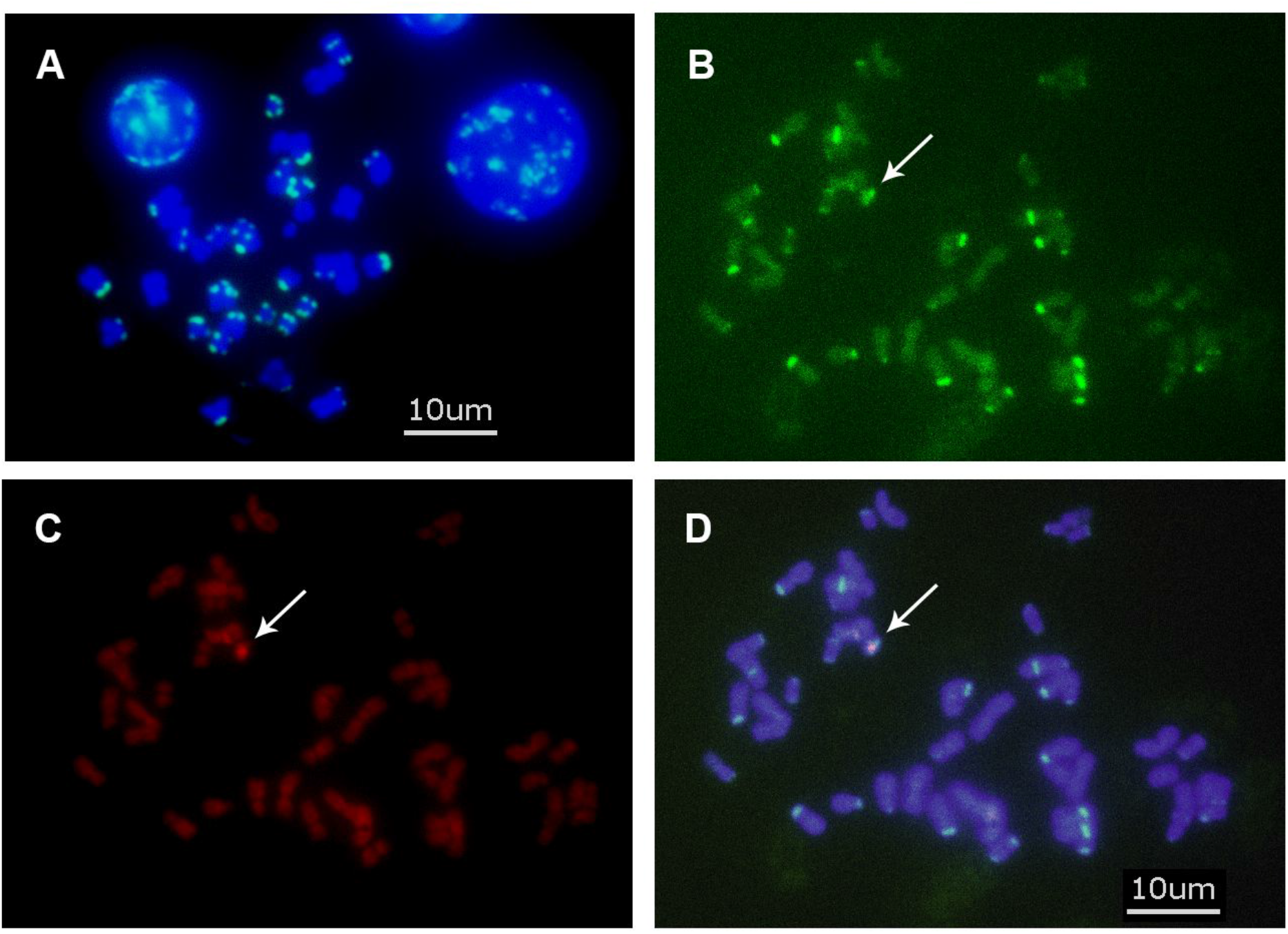
Fluorescent in situ hybridization (FISH) analysis. Hybridization of: **(A)** the 5’-amine-modified probe Pan32 (with a candidate 32-mer male-biased motif sequence) to DAPI-counterstained chimpanzee male chromosomes; **(B)** the 5’-amine-modified probe Pan32 to bonobo male chromosomes; **(C)** the whole bonobo Y chromosome painting probe (WBY) to bonobo male chromosomes; and **(D)** both the WBY and Pan32 probes to DAPI-counterstained bonobo male chromosomes. The arrow indicates the location of the bonobo Y chromosome. The 5’-amine-modified probe Pan32 is labeled with Alexa Fluor (green). The WBY probe is labeled with digoxigenin (red). Scale bar = 10 um.

### Estimating satellite repeat abundance and length with long-read data

Because short-read technologies can only provide information about total repeat abundances, and satellite repeats are routinely under-represented in sequenced assemblies, one can take advantage of long reads, e.g., as produced by Nanopore or PacBio, to provide a presumably less biased view of repeated array lengths. Unfortunately, adequate software to retrieve repeats from long sequencing reads, which are notoriously error-prone (with error rates around ∼15-16% for both PacBio and Nanopore (Jain et al. 2018a; Rhoads and Au 2015)), does not currently exist. To address this limitation, we developed Noise-Cancelling Repeat Finder (NCRF; (Harris et al. 2018)), a stand-alone software that can recover repeat length distributions from long reads notwithstanding their high error rates. NCRF initially identifies continuous arrays of highly similar repeated motifs (imperfect repeats). This is vital as arrays comprising a dominant motif and one or more derived motifs represent an important facet of biological variability (Plohl et al. 2008). Since the direct *de novo* identification of satellite repeats from error-prone long reads is challenging, we used the 39 abundant Illumina-derived repeated motifs identified above (see section ‘Repeat identification in short reads’) as queries for the screening of long reads by NCRF.

To evaluate densities and lengths of these 39 motifs in long read technologies using NCRF, we sequenced six great ape individuals, one from each species of great apes, on one Nanopore MinION flow cell (Table S7-8), and employed publicly available PacBio sequencing reads available for four great ape species (Table S7 and S9)(Gordon et al. 2016; Kronenberg et al. 2018). For our Nanopore data, the longest observed read was 206 kb and the read length N50 ranged from 26 to 37 kb among samples (Table S8). In comparison, using a single flow cell of publicly available PacBio data for each species, the longest observed read was 184 kb and the read length N50 ranged from 19 to 34 kb among samples (Table S9). Concerning repeat densities we found with NCRF (Fig. 5), for both PacBio and Nanopore reads, the general patterns were consistent with those inferred from Illumina reads with TRF (Fig. 1B) – however the exact densities differed. The differences can be in part explained by the fact that different individuals of the same species were sequenced using each technology. An additional factor could be that Nanopore and PacBio reads employ distinct library preparation and sequencing protocols that are prone to different biases (see Discussion). Interestingly, some of the repeated motifs abundant in short-read data, such as the (AATGG)_n_ repeat, were not as abundant in long-read data.

**Figure 5.**
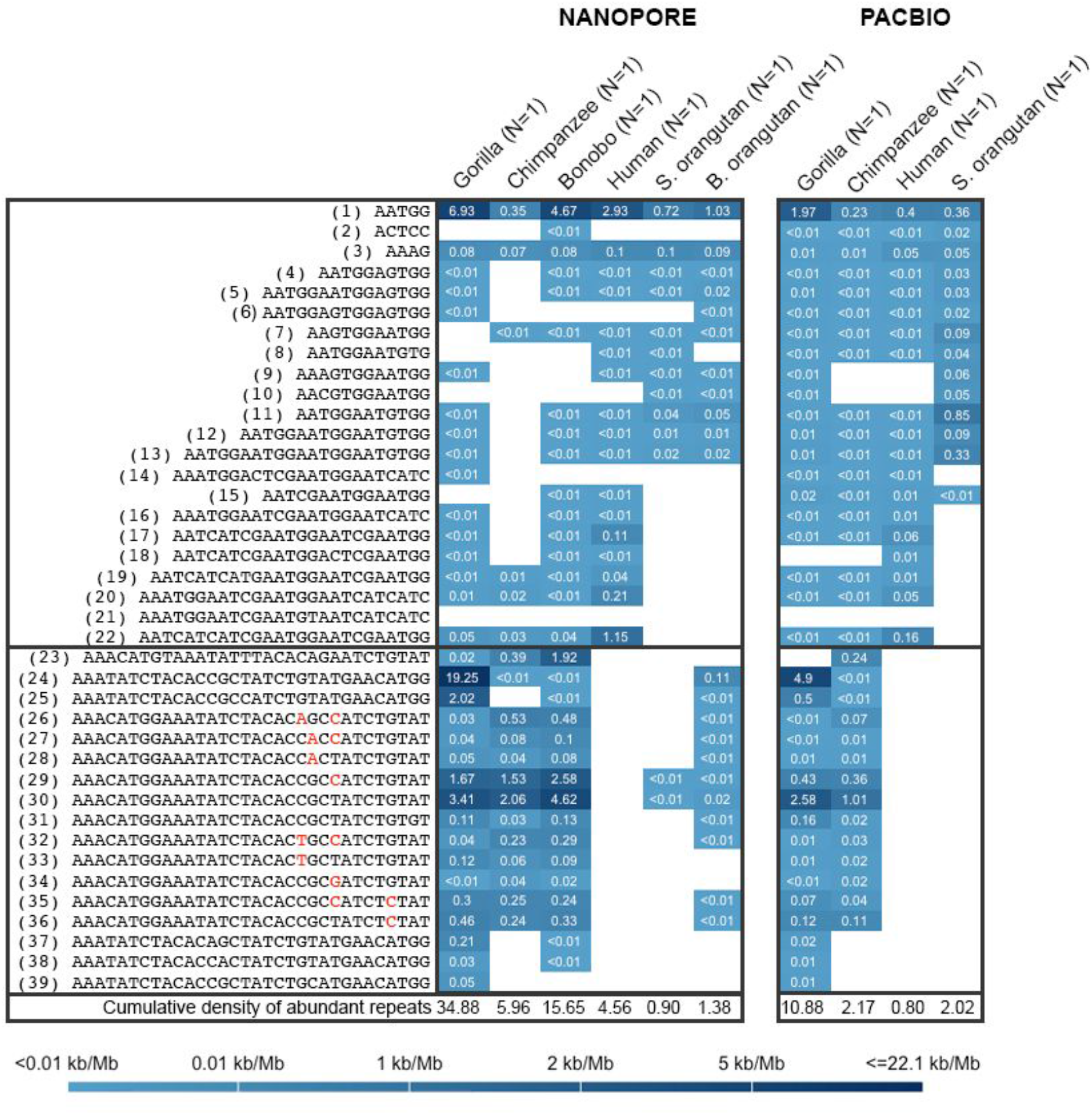
Repeat densities inferred from long sequencing reads generated with Nanopore (in-house) and PacBio (from public datasets) technologies. Note that left and right panels represent different individuals. The whole-genome PacBio data for bonobo and Bornean orangutan are not available.

We also discovered that long satellite arrays were frequently a mix of more than one motif, present in perfect patches interspersed with highly similar, yet different, sequences. To come to this conclusion, we proceeded as follows. First, we verified that long reads were able to capture the full lengths of satellite repeats (Figs. S14-S15), as demonstrated by the fact that in the majority of cases long reads encompassed complete repeat arrays (depending on the species, 90-95% and 99% of repeat arrays were nested within individual reads in Nanopore and PacBio, respectively, Table S10). The longest repeat arrays we recovered were for (AATGG)_n_ and 32-mers (Fig. 6), some of which were over 59 kb (Table S11, Fig. S14). Last, we focused on the arrays with a single dominant motif and, depending on the species, found that at least 10-25% of all arrays were composed of a mix of different repeated motifs (Table S12). This is likely an underestimation, as we only detected overlaps in repeat annotations among the 39 most abundant repeated motifs. With PacBio, the longest repeat arrays we recovered were over 17 kb (Table S11, Fig. S15). Taken together, our results suggest frequent interspersion of perfect repeats with highly similar repeated motifs.

**Figure 6.**
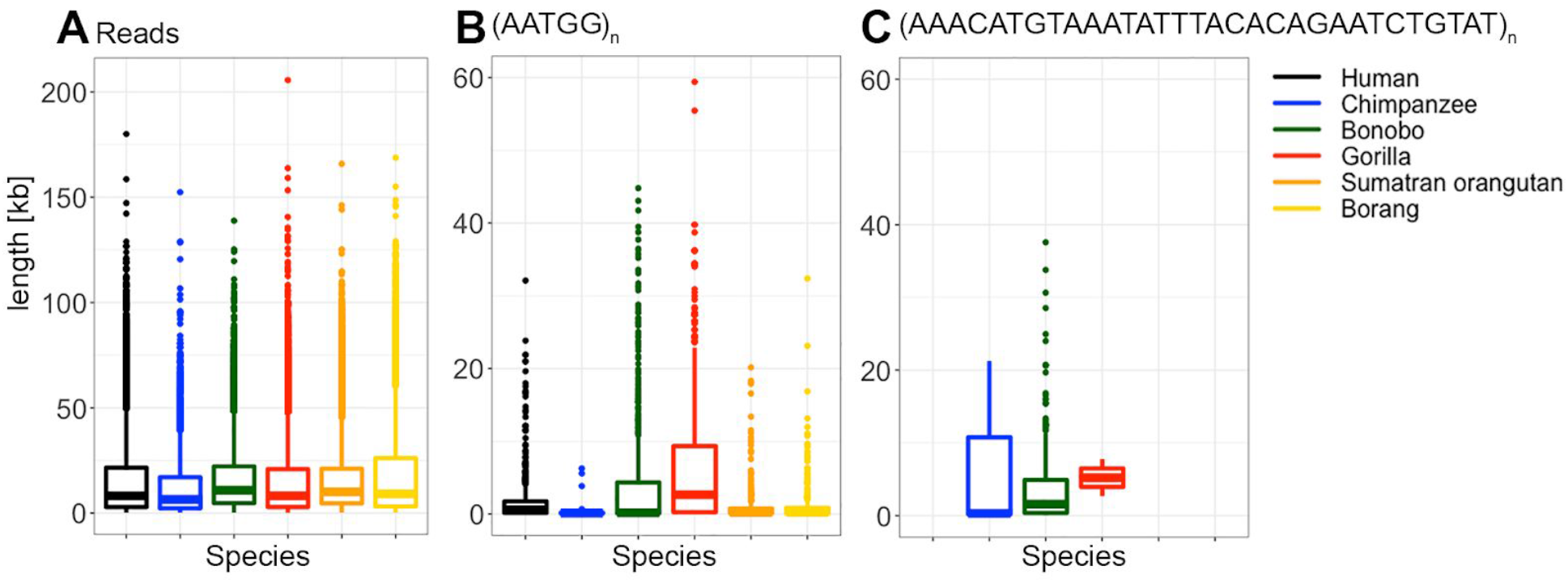
Box plots of lengths of (A) reads, (B) repeated motif (AATGG)_n_, and (C) one 32-mer recovered, from Nanopore data.

As a control, we studied the repeat density of the telomeric (TTAGGG)_n_ satellite using these long-read data, even though it is not one of the 39 most abundant repeats. The repeat density of this satellite was rather low for both technologies (the ranges for its density across species were 0.00194-0.0330 kb/M and 0.0110-0.0974 kb/M for Nanopore and PacBio, respectively, Fig. S16A-B, Table S3), consistent with our findings from Illumina reads (Table S3). Nevertheless we still found a substantial number of long (>500-bp) arrays of (TTAGGG)_n_ in PacBio (the longest arrays were 10.4, 3.9, 7.2 and 10.4 kb for human, chimpanzee, gorilla and Sumatran orangutan, respectively, Fig. S16E) and also some in our smaller-scale Nanopore data (the longest arrays were 0.8 kb and 4 kb for human and Bornean orangutan, respectively, Fig. S16D). Moreover, these telomeric satellite arrays were predominantly located towards the ends of reads, further implying their telomeric location (Fig. S16).

## Discussion

Satellite repeats constitute a large portion of the human genome (Jain et al. 2018a; Spinelli 2003), yet they have been routinely underexplored in the genomes of great apes (Kronenberg et al. 2018). Our study fills this gap; it provides a detailed characterization of this important component of hominid genomes and demonstrates a remarkable divergence of satellite repeats with unit sizes ≤50 bp among ape species separated by less than 14 MY (Glazko and Nei 2003).

### Satellite repeats in great ape genomes

#### The (AATGG)_n_ repeat and its derivatives

We determined the (AATGG)_n_ repeat to be abundant in great ape species. Independent of sequencing technology used, its density was usually highest in gorilla (second highest with Nanopore), rather high in orangutans, human, and bonobo, and lowest in chimpanzee (Figs. 1B and 5). This is in agreement with a suggestion that, during primate evolution, amplification of HSat3, for which the (AATGG)_n_ repeat is the source, peaked in gorilla and orangutan lineages (Jarmuż et al. 2007). We also found high intraspecific variability in the density of (AATGG)_n_, sometimes reaching up to 1.51-fold pairwise difference between individuals of the same species (Table S4). These findings strongly argue for the rapid evolution of this repeat.

We found that (AATGG)_n_ is ubiquitously present in all great ape individuals in our study, suggesting that it performs an important function. It is located at pericentromeric regions of acrocentric chromosomes (Lee et al. 1997), can fold into a non-B DNA conformation (Grady et al. 1992; Zhu et al. 1996; Chou et al. 2003), and was suggested to participate in forming centromeres (Grady et al. 1992). Importantly, under conditions of stress, the (AATGG)_n_ repeat is transcribed from three to four 9q12 loci into long noncoding RNAs which, together with several proteins, form nuclear stress bodies and play a critical role in heat shock response (Nakahori et al. 1986; Jolly et al. 2004; Goenka et al. 2016; Biamonti and Vourc’h 2010). In fact, such RNAs were recently shown to be required to “provide full protection against the heat-shock-induced cell death” via contributing to transcriptional silencing (Goenka et al. 2016). Some of these RNAs can be very long (Jolly et al. 2004), with polyadenylated transcripts ranging from 2 to >5 kb (Goenka et al. 2016). In agreement with this observation, we found that some (AATGG)_n_ imperfect arrays, which can be part of these transcripts, can be over 59 kb long.

Our study has also identified abundant repeated motifs that were derived from (AATGG)_n_ (Figs. 1B and S5). Interestingly, some of them, including the (AATGG)_n_ repeat itself, are matching substrings of the most common 24-mers indicative of a specific HSat subfamily (Altemose et al. 2014) — either with no mismatches (AATGG, ACTCC, and AAAG) or with one mismatch (AATGGAATGGAGTGG, AATGGAGTGG, AATGGAATGTG, AATCGAATGGAATGG). This provides an independent confirmation that they form satellite repeats.

#### Subterminal Satellites

Another interesting group of satellite repeats highlighted by our study are the phylogenetically related, AT-rich 32-mers also called Subterminal satellites (StSats) due to their proximity to telomeres, as demonstrated by our and other studies (Royle et al. 1994)(Ventura et al. 2012; Royle et al. 1994; Koga et al. 2011)(Royle et al. 1994). Independent of the sequencing technology used, we found that these repeats are highly abundant in gorilla, still very abundant in chimpanzee and bonobo, but absent in human. These findings corroborate early studies hypothesizing that these repeats were present in the common ancestor of hominids (albeit in small amounts), and then lost in the human lineage (Ventura et al. 2012; Royle et al. 1994; Koga et al. 2011). The loss of StSats in orangutans was also proposed (Ventura et al. 2012; Royle et al. 1994; Koga et al. 2011), however our analysis suggests that such loss was incomplete, as we can still find StSat traces in orangutan genomes using both Illumina and Nanopore read data. Consistent with the notion of a partial loss in orangutans, StSats are polymorphic in their presence/absence among orangutan individuals (Fig. S7A). In contrast, the majority of StSats are present in all gorilla, chimpanzee and bonobo individuals analyzed, suggesting that they might be functionally important in their genomes. Various roles for StSats have been proposed, including participation in meiosis (Ventura et al. 2012; Royle et al. 1994; Koga et al. 2011), telomere clustering and metabolism, as well as the regulation of replication timing in the vicinity of telomeres (Novo et al. 2013).

#### Male-biased repeats

Leveraging differences in repeat density between males and females, we identified 18 candidate male-biased repeats in great apes (Table S6). These included the (AATGG)_n_ repeat, which was previously shown to be present on the human Y chromosome as the primary repeated unit of its three common satellites (DYZ1, DYZ17, and DYZ18) (Skaletsky et al. 2003; Kunkel et al. 1976), and on the Y chromosome of orangutan, gorilla and chimpanzee/bonobo with FISH (Jarmuz et al. 2007). Additionally, we found several StSats to be male-biased and confirmed their presence in the gorilla Y assembly and in the bonobo Y chromosome using FISH (Fig. 4). This substantially increases the current knowledge of both candidate and validated Y chromosome heterochromatic repeats in great apes. Prior to our study, these repeats were underexplored because only human, chimpanzee and gorilla Y chromosome assemblies are currently available and such assemblies are mostly euchromatic (Skaletsky et al. 2003; Hughes et al. 2010; Tomaszkiewicz et al. 2016).

Differences in heterochromatin density can be one of the major contributors to the dramatic length differences observed among the Y chromosomes of great apes (Gläser et al. 1998; Hughes et al. 2010). To shed light on this, we tested whether the differences in satellite repeat content between males and females, presumably reflecting the Y chromosome repeat content, corresponds to the differences in lengths of great ape Y chromosomes. For instance, the difference in content of male-biased satellite repeats between males and females is 13.1, 8.0 and 2.5 kb/Mb for gorilla, bonobo, and chimpanzee, respectively (Table S14). In agreement with the order of these values, cytogenetic estimates indicate that, among the Y chromosomes of these three species, the gorilla’s is the longest Y chromosome, the bonobo’s is intermediate, and the chimpanzee’s is the shortest (Gläser et al. 1998). Therefore, satellite repeats may indeed be playing an important role in determining Y chromosome length variation in great apes. However, they are likely not the sole contributors; indeed, the difference in satellite repeat content between males and females for humans is only 2.0 kb/Mb, despite the fact that the human Y chromosome falls between bonobo and gorilla Ys in length (Gläser et al. 1998).

It was proposed that enrichment of different, or accumulation of unique, satellite DNA is the first step in separation of the X and Y chromosomes (Brutlag 1980). It was also hypothesized that the composition of the heterochromatin on the Y may differ from that on other chromosomes because of (1) absence of recombination; (2) a potential role of heterochromatin in silencing the Y; and (3) the small effective population size of the Y (Bachtrog 2013; Nei 1970; Charlesworth and Charlesworth 2000). Consistent with these hypotheses, some *Drosophila* species (*D. virilis*, *D. melanogaster*, *D. simulans*, and *D. sechellia*) exhibited many Y-enriched or Y-specific satellite repeats (Wei et al. 2018). In contrast, other *Drosophila* species (*D. pseudoobscura* and *D. persimilis*) have prominent abundance of transposable elements (TE) on the Y (Wei et al. 2018) — suggesting that Y chromosome degeneration occurs by satellite repeat accumulation in some species, and TE accumulation in others. These two alternatives can be explored also for the great ape Y chromosomes, once the assemblies that are currently missing become available.

The Y chromosome heterochromatin is a major source of epigenetic regulation, modulating phenotypic variation in natural populations (Lemos et al. 2010). For instance, in *Drosophila*, its content and length affect expression of autosomal genes (Lemos et al. 2008). Similarly, a repeat-rich non-coding RNA was recently found to play a role in regulating the expression of several genes in mouse testis (Reddy et al. 2018). Such a phenomenon in primates is yet to be investigated.

#### Co-occurrence of satellite repeats

Our observations suggest dependencies among the densities of many repeated motifs, and an underlying structure in their distribution in the great apes genomes – which is at least partially dictated by sequence similarity and evolution, stemming from the interspersion of longer satellite arrays with similar motifs. This echoes recent observations made for *Drosophila* (Wei et al. 2014, 2018) and *Chlamydomonas reinhardtii (Flynn et al. 2018)*. Similarly to the pattern observed in *Drosophila*, in great apes clusters of co-occurring repeats are in part driven by their sequence similarity. Several hypotheses were proposed to explain such a pattern; for instance, many similar repeat motifs can serve as recognition sites for the same DNA-binding proteins (Wei et al. 2014), and correlated motifs might be physically linked to each other due to a large-scale duplication or due to interspersion. An example of interspersion are two groups of HSat3 DNA: the first group is dominated by (AATGG)_n_ and the second group represents a mix of (AATGG)_n_ and (ACTCC)_n_ (Jarmuż et al. 2007). We also found antagonistic relationships among some repeats, in particular among (AAAG)_n_ and several other repeats. Again similar to observations made in *Drosophila* (Wei et al. 2014), this can occur when the expansion of one repeat type comes at the expense of another. The differences we found in nature and strength of dependencies among repeat densities in various great apes might be explained by differences in the overall tolerance their genomes have towards repetitive load. Future studies should incorporate data on long-distance genome interactions (e.g., Hi-C) to further explore repeat co-occurrence patterns in great ape genomes.

### Interspecific differences and lack of phylogenetic signal in repeat densities

We found drastic differences among great ape species in overall repeat content. Independent of the sequencing technology used, overall repeat density was highest in gorilla, intermediate in chimpanzee and bonobo, and lowest in human and orangutans (Figs. 1B and 5). This is primarily explained by the absence or paucity of StSats in human and orangutans, respectively. Also, while clustering based on repeat densities did correctly assign individuals into species, subsequent agglomeration did not follow the expected species phylogeny. In particular, we frequently observed chimpanzee, bonobo, and gorilla clustering together, and human clustering with orangutan (Fig. 2C). We found that similarities among chimpanzee, gorilla and bonobo individuals (Figs. S10F-H) were in part driven by StSats (data not shown) — but in certain instances they clustered together even after the exclusion of such repeats (Fig. S10I). Several explanations are possible for this unexpected observation, including incomplete lineage sorting (Kronenberg et al. 2018), parallel gains of the same repeats along different lineages, molecular drive, and segregation distortion (reviewed in (Wei et al. 2018)). Future studies should examine each of these explanations in detail. At present, what is clear is that satellite repeats have a notably high tempo of turnover and, at least at the timescale resolution of great ape evolution, do not carry phylogenetic signals. Interestingly, our results are more similar to those found for Drosophila populations than for Drosophila species (Wei et al. 2014; Wei et al. 2018).

### The power of long reads, study caveats, and future directions

One of the strengths of our study is in that we combined information from three different sequencing technologies to investigate satellite repeats. We identified repeated motifs from rather accurate short-read (Illumina) data, and augmented information about them using long reads from the Nanopore and PacBio platforms. Critically, we studied satellite repeats from sequencing *reads*, and not from *reference genomes*, thus greatly expanding our current knowledge about yet unassembled portions of great ape genomes. The use of data from long reads has allowed us to gain reliable information on repeat length. Indeed, depending on the species and technology, 90 to 99% of the repeat arrays in our study were wholly contained within single sequencing reads (Table S10). The longest repeat arrays were 59 kb and 17 kb in length, as identified using the Nanopore and PacBio platforms, respectively. Such lengths are unprecedented; the recent PacBio-augmented assembly of the sooty mangabey (a primate) identified a 52-kb repeat array, and this was the longest found in an analysis comprising as many as 719 assembled eukaryotic genomes (Surabhi et al. 2018). Our study confirms that long-read technologies are indeed suitable for the analysis of long heterochromatic satellites. This is due both to their progressively increasing read lengths, and to recent advances in the algorithms used to tackle their noisy error profiles, e.g., NCRF (Harris et al. 2018). Deciphering repeat lengths and structures will enable genotyping and assigning potential functions to a larger set of repeat arrays than previously possible. For example, Sonay and colleagues showed gene expression divergence between human and great apes to be higher for genes that encompassed tandem repeats (TRs) (Sonay et al. 2015). However, since their study required TRs to be fully encompassed within short Illumina sequencing reads, they were able to analyze only 58% of TRs present in the human reference. Nanopore sequencing was recently used to characterize the first complete human centromere on the Y chromosome (Jain et al. 2018b) and to determine the lengths of human telomeric repeats (Jain et al. 2018a). We expect a growing interest in tools and approaches operating directly on raw, ultra-long reads (Lower et al. 2018).

Many of our conclusions are robust to the use of sequencing technology. However, we did find differences in the exact values of repeat density estimates obtained from the three technologies we considered. These differences could be due to the use of different individuals between short-read and long-read technologies, but also due to the vastly different library preparation and sequencing protocols. While Illumina reads always represent short fragmented DNA, long DNA molecules used for PacBio and Nanopore sequencing could form secondary structures. We have recently shown that non-B DNA structures can affect PacBio sequencing depth and error rates (Guiblet et al. 2018). For Nanopore, fragments harboring these structures might not pass through the pores. In both cases, the representation of repeats capable of forming non-B DNA might be altered. This, for instance, might explain at least in part why the (AATGG)_n_ repeat, known to form a non-B DNA structure (Grady et al. 1992), is underrepresented in Nanopore and PacBio vs. Illumina data (Figs. 1B and 5). The telomeric repeat (TTAGGG)_n_ is known to adopt a G-quadruplex formation and this might also affect its low density in sequencing reads. Moreover, different genome k-mers are not represented equally in Nanopore sequencing, an issue that is being mitigated by advances in the Nanopore base calling algorithms (Lu et al. 2016; Ip et al. 2015). The Illumina short-read sequencing used in the first part of our study might have its own issues. The APD and HGDP sequencing libraries we analyzed were generated with the PCR+ protocol. This might have led to an overestimation of repeat densities or difficulties with sequencing of the extremely GC-rich fragments. However, human repeat densities were very similar when estimated from PCR+ vs. PCR-samples (Figs. 1B and S4), and we observed each repeat motif at each locus to be affected by PCR amplification at approximately the same rate (Supplementary Note 2). In *Drosophila* (Wei et al. 2018), omission of the PCR step improved correlation of satellite abundances between replicates. It is much more expensive to generate PCR-data on a large scale in apes than in *Drosophila*, especially when intraspecific variation, and thus multiple individuals, are of interest. However, such data should definitely be generated for great apes in the future. In this study, we did not perform the GC-bias correction (Benjamini and Speed 2012) that was employed in some other studies (e.g., (Wei et al. 2018; Flynn et al. 2017). Available GC-correction pipelines require reference genomes and are thus unsuitable for whole-genome sequencing reads with suboptimal or missing references (e.g., for Y chromosomes in most apes).

Our study focused on relatively short repeated units (<50-bp), because we identified satellite repeats from short reads (two 50-bp repeats fit a 100-bp read). Our use of such short-motif repeats as a proxy for heterochromatin is justified based on several considerations: (1) they are part of long arrays, as identified by long-read data; (2) some of them match to 24-mers differentiating HSat families (Altemose 2014); and (3) some of them have (sub)telomeric locations, as demonstrated by our FISH experiments (Fig. 4). Repeats with longer units were not considered because the computational tools to identify them *de novo* in noisy long reads do not currently exist. Some studies focused on the analysis of the 171-bp centromeric heterochromatic arrays whose sequence in the human genome has been well characterized (Jain et al. 2018b; Miga et al. 2014; Melters et al. 2013). Analyzing repeats with longer repeat units in great apes will be of great interest for future studies, once algorithms to reliably identify novel repeats from noisy long reads are developed.

## Methods

### Sequencing data and quality filtering

From the ADP (Prado-Martinez et al. 2013), we focused on 399 fastq files with forward reads because they surpassed those with reverse reads in both sample size and quality (the latter was computed using FastQC v0.11.2 for all files using 10 randomly selected reads per file). Ape individuals sequenced in multiple Illumina sequencing lanes/runs were kept separately for all the downstream processing and treated as technical replicates. Excluding 39 files with read lengths shorter than 52 bp resulted in 360 files (322, 32, and 6 files with read lengths 100 bp, 101 bp, and 151 bp, respectively). Subsequently, excluding 51 files with read counts smaller than 20,000,000 (to avoid potential sampling bias resulting from low read counts) resulted in 309 files. The files belonging to genetically close relatives to other samples (Bulera, Kowali, Suzie and Oko)(Prado-Martinez et al. 2013) were also removed, resulting in 295 fastq files. To avoid sequence bias revealed by QC analysis (over-represented k-mers present profusely toward read ends) and to remove potential sequencing errors, we discarded all reads that contained at least one base pair with a Phred quality score below 20 using FASTX-Toolkit (version 0.0.13, fastq_quality_filter −Q33 −v −q 20 −p 100).

### Identification of Repeats

Reads retained after such filtering were converted from fastq to fasta format and repeats in them were identified with TRF (version trf409.legacylinux64, parameters MATCH=2 MISMATCH=7 DELTA=7 PM=80 PI=10 MINSCORE=50 MAXPERIOD=2000 −l 6 −f −d −h −ngs) (Benson 1999). The resulting repeats were parsed using the script parseTRFngs.py (see GitHub repository) that implements collapsing of the same group of repeats (shifts and reverse complements) into a single representative. We required each repeat array to be at least 75 bp in length. Finally, we used median repeat densities across all technical replicates to compute satellite repeat densities for each individual. To verify that technical replicates from the same individual were consistent in their repeat estimates, we measured the tightness of these estimates computing intraclass correlation coefficients between technical replicates for the 100 most abundant repeats (we used the R package ICCbare). The median intraclass correlation coefficient was 0.96 (Fig. S1).

To avoid duplicates in the output, the recovered repeats were further filtered and formatted. Namely, we merged all repeats that shared the basic repeated unit and were in close vicinity (less than the minimal unit length of the two neighboring repeats) to each other. Reads containing the same repeated motif can map to either reference or reverse strand, and the annotated repeats can start with a different leading nucleotide. Thus, we report the data on occurrences of a repeated motif whose phase was chosen alphabetically, and combine the data for motifs and their reverse complement sequences. Because the same long stretches of repeats can have different beginnings (e.g. AATGG and ATGGA differ by a 1-bp shift) or can be present on different strands (e.g. AATGG and CCATT), we reformatted all repeats into the lexicographically smallest rotations. This means that for all possible rotations (1-bp shift followed by 1-bp increments of shift size up to the unit length) and both possible strands, we picked only one representative. This representative is the first repeat in alphabetical order out of all generated possibilities that we described above.

### Calculation of repeat frequency and density

We required each repeated motif to be present at ≥100 loci per 20 million reads. For repeated motifs that passed these filters, we calculated the corresponding repeat densities after normalizing for the read length and the read count after filtering. To calculate repeat density for each species, we included only those repeats that were present in at least one individual of that species. In order to display repeat densities in the heatmaps, they were first converted to kb/Mb and then rounded to two decimal places.

### Correlations of repeat co-occurrences

To assess the significance of observed correlations of repeat motifs (using Spearman coefficient and ranks based on the repeat density), we generated 10 reshuffled datasets of the original repeat densities of 39 abundant repeats separately for each species (visualized as grey band in Fig. 3). Reshuffling was done as follows: in a matrix of individuals x repeats, we kept the content of the matrix, but randomly re-assigned column names, so that the biological associations among repeats were broken and those occurring were due to chance.

### Sequence similarity and inter-relatedness among the 39 most abundant repeated motifs

The sequence similarity was calculated using MEGA7 (Kumar et al. 2016). Only substitutions (and not insertions or deletions) were considered. The pairwise distances were calculated using the number of differences (both transitions and transversions) and treating gaps with pairwise deletion (Fig. S5). For each species, we calculated mean repeat density across all individuals.

### Length distribution for long reads

Repeated motifs were identified in long reads using NoiseCancelingRepeatFinder, version 0.09.03 (Harris et al., 2018, submitted). The current version of the algorithm can be downloaded from: github.com/makovalab-psu/NoiseCancellingRepeatFinder/. For more detailed information on how to run the program, see Supplementary Note 3. For PacBio and Nanopore sequencing, --scoring=pacbio (M=10 MM=35 IO=33 IX=21 DO=6 DX=28) and --scoring=nanopore (M=10 MM=63 IO=51 IX=98 DO=27 DX=34) options were used, respectively. The maxnoise parameter was set to 20% to retain long reads with noisy repeat arrays. Subsequently, the repeated arrays were analyzed for their motif composition and each array was assigned to a motif that comprises more than 50% of an array.

### Experimental validations of male-biased repeats

#### Preparation of the probes

The whole bonobo Y chromosome painting probe (WBY) was prepared from flow-sorted bonobo Y chromosomes and labeled with biotin-16-dUTP (Jena BioScience) using DOP-PCR according to (Yang et al. 2009). Oligonucleotide probe (Pan32) (/5AmMC12/ATCTGTATAAACATGGAAATATCTACACCGCY) was prepared and labeled using Alexa Fluor oligonucleotide amine labeling kit (Invitrogen).

#### FISH

Metaphases were prepared from chimpanzee male and female lymphoblastoid cell line and from bonobo male fibroblast cell line following a standard protocol of colcemid treatment, hypotonization and methanol/acetic acid fixation (Howe et al. 2014). Slides were pre-treated with acetone for 10 min and aged at 65°C for 1 h. Subsequently, the slides were denatured in the alkaline solution (Sigma) for 5 min, followed by neutralization in 1M Tris-HCl, pH 7.5, and one wash in 1x PBS for 4 min. Next, a series of dehydration washes were performed as follows: 70% EtOH at −20°C for 4 min, 70% EtOH for 2 min, 90% EtOH for 2 min, and 100% EtOH for 4 min. The WBY probe was denatured in hybridization buffer at 75°C for 15 min and pre-annealed at 37°C for 30 min. Subsequently, 25 ng of the Pan32 probe was applied to the hybridization area and incubated at 37°C for 12 h for chimpanzee male chromosomes as well as for bonobo male chromosomes. In a separate FISH experiment, the mix of 25 ng of WBY and 25 ng of Pan32 was applied to the hybridization area and incubated at 37°C for 24 h for bonobo male chromosomes and for 48 h for chimpanzee male and female chromosomes (cross-species FISH). Post-hybridization washes were performed in 0.5x SSC at 50°C for 5 min, 2x SSCT at 37°C for 5 min, and 1x PBS at at 37°C for 5 min. For slides with the mix of probes, an additional step of probe detection with Cy3-Streptavidin (Sigma) was applied. Slides were stained with DAPI (Vector Laboratories) and visualized under the Keyence BZ-9000 fluorescence microscope. Photodocumentation was performed using the 100x immersion objective and the images were analyzed using BZ-Viewer and BZ Analyzer.

#### Nanopore library preparation and sequencing

DNA was extracted from male cell lines of bonobo (AG05253, Coriell Institute), gorilla (KB3781, “Jim”, San Diego Zoological Society), Bornean orangutan (AG05252, Coriell Institute), and Sumatran orangutan (AG06213, Coriell Institute) using the MagAttract High Molecular Weight DNA Kit (Qiagen, Germany). Male chimpanzee DNA sample (CH159, “Rock”) was provided by Dr. Mark Shriver and was acquired from the Bastrop Research Center. Human male DNA (J101) was provided by the University of Chicago.

Residual RNA was removed by digesting 3.5 µg of extracted DNA with 10 µg RNase A (Amresco) at 37 °C for 1 h, followed by purification with 1 volume of AMPure XP beads (Beckman Coulter). DNA integrity was visualized on a 0.5% agarose gel, DNA purity was determined with NanoDrop, and the concentration was measured with a Qubit broad-range assay. Libraries were prepared with the Native Barcoding Kit 1D (PCR-free) and the Ligation Sequencing Kit 1D (Nanopore) starting with 2 µg DNA per sample. DNA repair and end-repair were combined in one step as described in the 1D gDNA long reads without BluePippin protocol (version: GLRE_9052_v108_revB_19Dec2017; updated: 10/01/2018). Barcoding and adapter ligation were performed as indicated in the 1D Native barcoding genomic DNA (with EXP-NBD103 and SQK-LSK108) protocol (version: NBE_9006_v103_revP_21Dec2016; updated: 16/02/2018), starting with 700 ng of end-prepped DNA per sample. 250 ng of barcoded DNA per sample were pooled and all further steps were performed according to the 1D gDNA long reads without BluePippin protocol. DNA low-binding tubes as well as wide-pore low-retention pipette tips were used for DNA handling in all steps. Sequencing was performed with a MinION using a flow cell of the type FLO-MIN106-R9.4 for 48 h. This resulted in 396, 55, 667, 526, 615 and 383 Mb of data (distributed among 26, 4, 43, 36, 40, and 22 thousand reads) for human, chimpanzee, bonobo, gorilla, Sumatran and Bornean orangutan, respectively.

## Data access

Illumina sequencing reads from 79 great apes were part of the Ape Diversity Project (Prado-Martinez et al. 2013). Sequencing reads generated for human populations were generated by (Meyer et al. 2012) Additionally, human samples from the Genome in a Bottle project (Zook et al. 2015) and two human trios from 1000 Genomes Project (1000 Genomes Project Consortium et al. 2015) — with IDs HG002, HG003, HG004, NA12889, NA12890, NA12877 and NA12891, NA12892, NA12878, respectively(1000 Genomes Project Consortium et al. 2015) — were used. The publicly available PacBio data had following ids: SRR2097942 for human, SRR5269473 for chimpanzee, ERR1294100 for gorilla, and SRR5235143 for Sumatran orangutan. The Nanopore data generated are deposited under the BioProject PRJNA505331. All scripts available from the git repository are at https://github.com/makovalab-psu/heterochromatin.

## Acknowledgments

We thank Wilfried Guiblet and Arslan Zaidi for valuable biological insights. We are grateful to Marzia Cremona, Kate Anthony, Oliver Ryder, Mark Shriver, Malcolm Ferguson-Smith, Jorge Pereira, and Shaun Mahony for their assistance. Research reported in this publication was supported by the National Institute Of General Medical Sciences of the National Institutes of Health under Award Number R01GM130691. The content is solely the responsibility of the authors and does not necessarily represent the official views of the National Institutes of Health. Funding was also provided by the Eberly College of Sciences, The Huck Institute of Life Sciences, and the Institute for CyberScience, at Penn State, as well as, in part, under grants from the Pennsylvania Department of Health using Tobacco Settlement and CURE Funds. The department specifically disclaims any responsibility for any analyses, responsibility, or conclusions.

## Author contributions

The study was conceived and designed by MC and KDM. MC performed the bioinformatics analysis. MT performed FISH experiments, RSH implemented NCRF and assisted with computational analysis of long reads, and BA performed Nanopore sequencing. FCH advised with the statistical parts of the paper. The manuscript was written by MC and edited by FCH and KDM. All authors read and approved the manuscript.

