## Supplementary Information for "High satellite repeat turnover in great apes studied with short- and long-read technologies"

### SUPPLEMENTARY MATERIALS

#### SUPPLEMENTARY NOTES

##### Supplementary Note 1. Robustness of the analysis with respect to read length and software choice

In order to verify that repeats identified from Illumina sequencing datasets represent real repeats and that they are not affected by the variable read lengths in the datasets used, or by the software choice for repeat identification, we performed two analyses. First, we re-ran repeat identification using the first five million reads of the dataset SRR741327 (a single Illumina run of a bonobo orangutan named Natalie from the Ape Diversity Project, APD (Prado-Martinez et al. 2013)) with the original 150 bp reads and then, separately, with the same reads trimmed to 100 bps (this length corresponds to the majority of the APD data). The resulting profiles were very similar (Figs. A-B). Second, we re-ran repeat identification on the same 100 bp reads using different software, Phobos (Phobos – a tandem repeat search tool for complete genomes, Mayer et al., 2010*). Both software packages identified motifs of the same unit size (Figs. A and C), although Phobos has reported fewer repeat arrays. Both recognized 5-mers and 32-mers as most abundant repeats. TRF (Benson 1999) in general uses more relaxed settings, identifying a larger number of imperfect repeats, especially of 32-mers.

*Mayer, Christoph, Phobos 3.3.11, 2006-2010, <http://www.rub.de/ecoevo/cm/cm_phobos.htm>

**Figure for Sup Note 1. (A)** The number of repeated arrays of a given unit size discovered using TRF and 100bp reads. The total number of repeated arrays discovered is 330,260. **(B)** The number of repeated arrays of a given unit size discovered using TRF and 150bp reads. The total number of repeated arrays discovered is 249,545. **(C)** The number of repeated arrays of a given unit size discovered using Phobos and 100bp reads. The total number of repeated arrays discovered is 69,970.

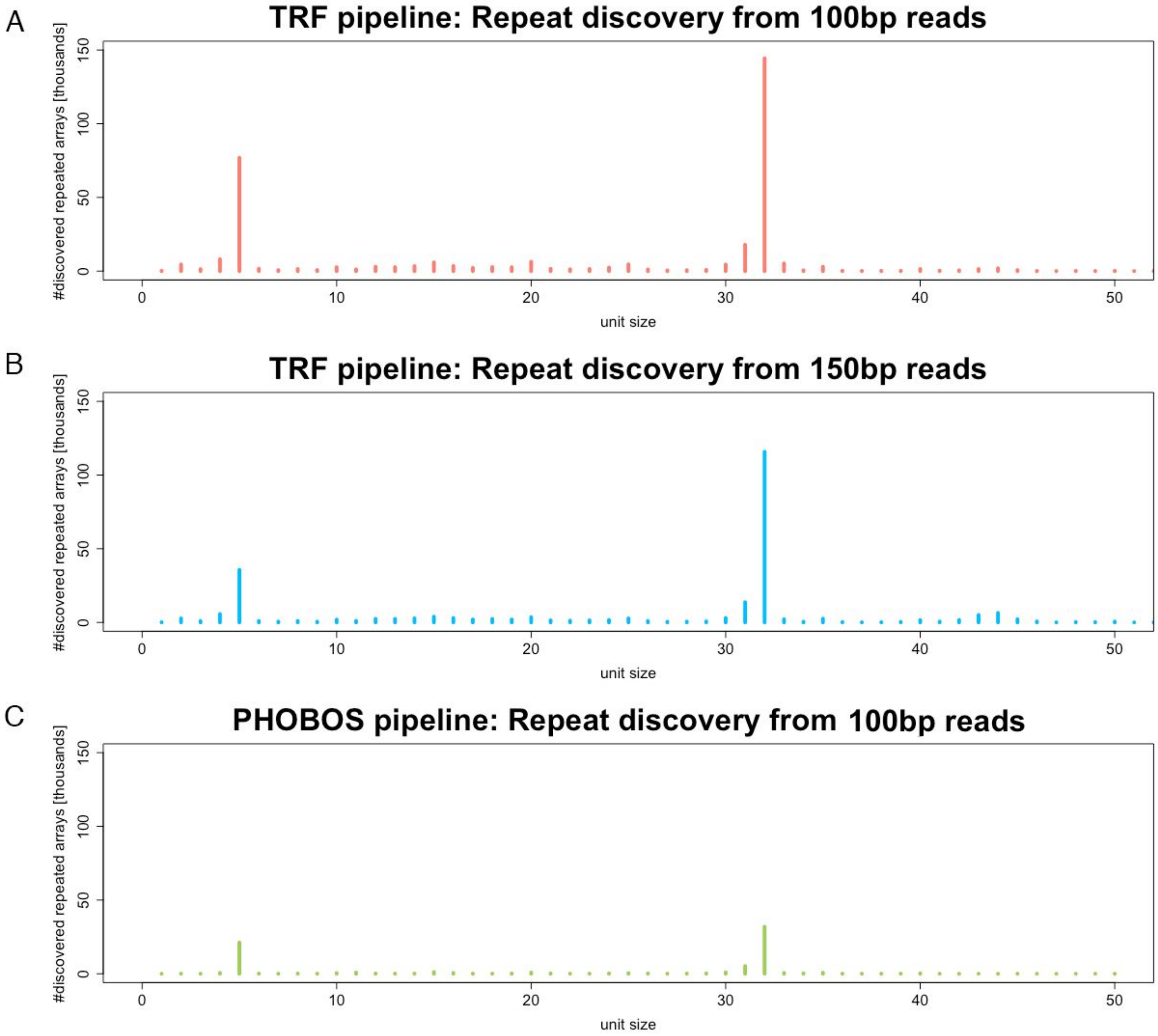

#### Supplementary Note 2. Potential effect of library preparation protocol on repeat counts and densities

We have used publicly available read data from sequencing libraries that were prepared according to the protocol described in (Prado-Martinez et al. 2013) at four sequencing facilities: CNAG, WashU, Seattle, and Stanford. All samples in that study (with the exception of samples of trios) underwent PCR amplification during library preparation and this step can introduce amplification bias (e.g. increased batch effect variability (Wei et al. 2017)). We, therefore, aimed at estimating how much the amplification step affects the repeat density of the satellites. We compared repeat density between one PCR- and two PCR+ libraries for the same orangutan individual from our previous publication (Fungtammasan et al. 2016). Using the same pipeline as for the Ape Diversity Project (ADP), the number of repeated motifs identified in at least one of the three orangutan libraries analyzed was 1,078.

We found that both PCR+ libraries reported higher repeat counts and repeat density than the PCR- library (see Figures below). Thus the presence of the PCR step led to an overestimation of the overall repeat counts and density. We corrected repeat counts by scaling them down by a factor derived from the corresponding libraries. This factor was 0.38 for the first replicate and 0.40 for the second replicate, resulting in the following formula: corrected_counts=(counts_PCR+) - slope*(counts_PCR+), where slope={0.38,0.40}. We corrected repeat densities by scaling them down by a factor derived from the corresponding libraries. This factor was 0.38 for the first replicate and 0.41 for the second replicate, resulting in the following formula: corrected_density=(density_PCR+) - slope*(density_PCR+), where slope={0.38,0.41}. Importantly, this analysis suggested that relative repeat densities remained unaffected and all repeated motifs were affected equally, with the linearly increasing effect for more abundant repeats. Even though these corrections can minimize the effects of amplification, we did not implement them in our analysis because correction factors were based on a comparison of only three libraries.

**Figure for Sup. Note S2.** (**A)** Higher repeat counts in two PCR+ libraries than in the corresponding PCR- library. The last plot shows that repeat counts in PCR- library and corrected PCR+ library correspond very tightly to each other. This suggests that linear increase in repeat counts (as the by-product of amplification) can be corrected using the proposed formula. **(B)** Higher repeat densities in two PCR+ libraries than in the corresponding PCR- library. The last plot shows that repeat densities in PCR- library and corrected PCR+ library correspond very tightly to each other. This suggests the linear increase in repeat densities (as the by-product of amplification) can be corrected using the proposed formula.

A

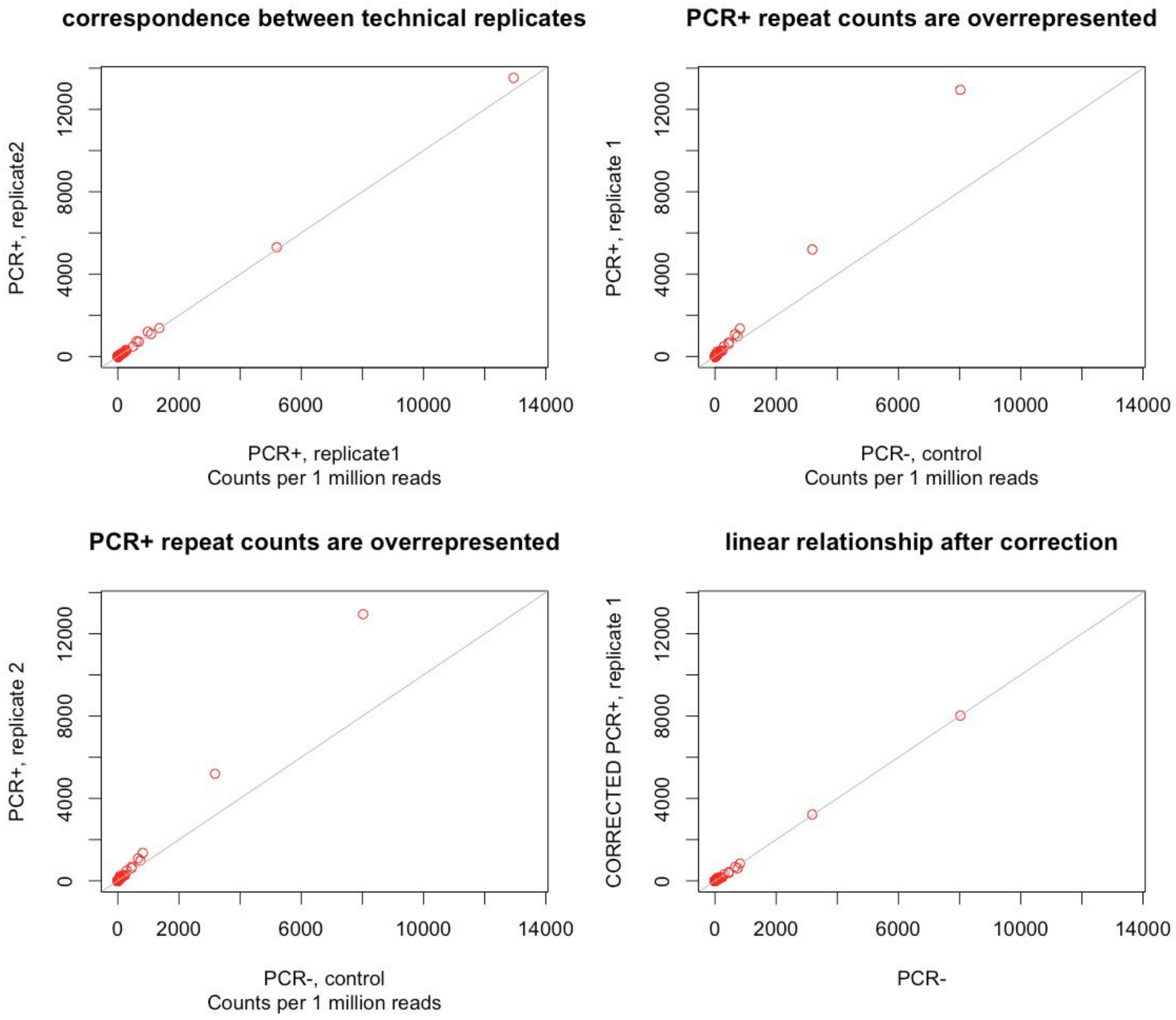
correspondence between technical replicates
PCR+, replicate2
PCR+, replicate1
Counts per 1 million reads
PCR+ repeat counts are overrepresented
PCR+, replicate 1
PCR-, control
Counts per 1 million reads
PCR+ repeat counts are overrepresented
PCR+, replicate 2
PCR-, control
Counts per 1 million reads
linear relationship after correction
CORRECTED PCR+, replicate 1
PCR-
0
2000
6000
10000
14000
4000
8000
12000

B

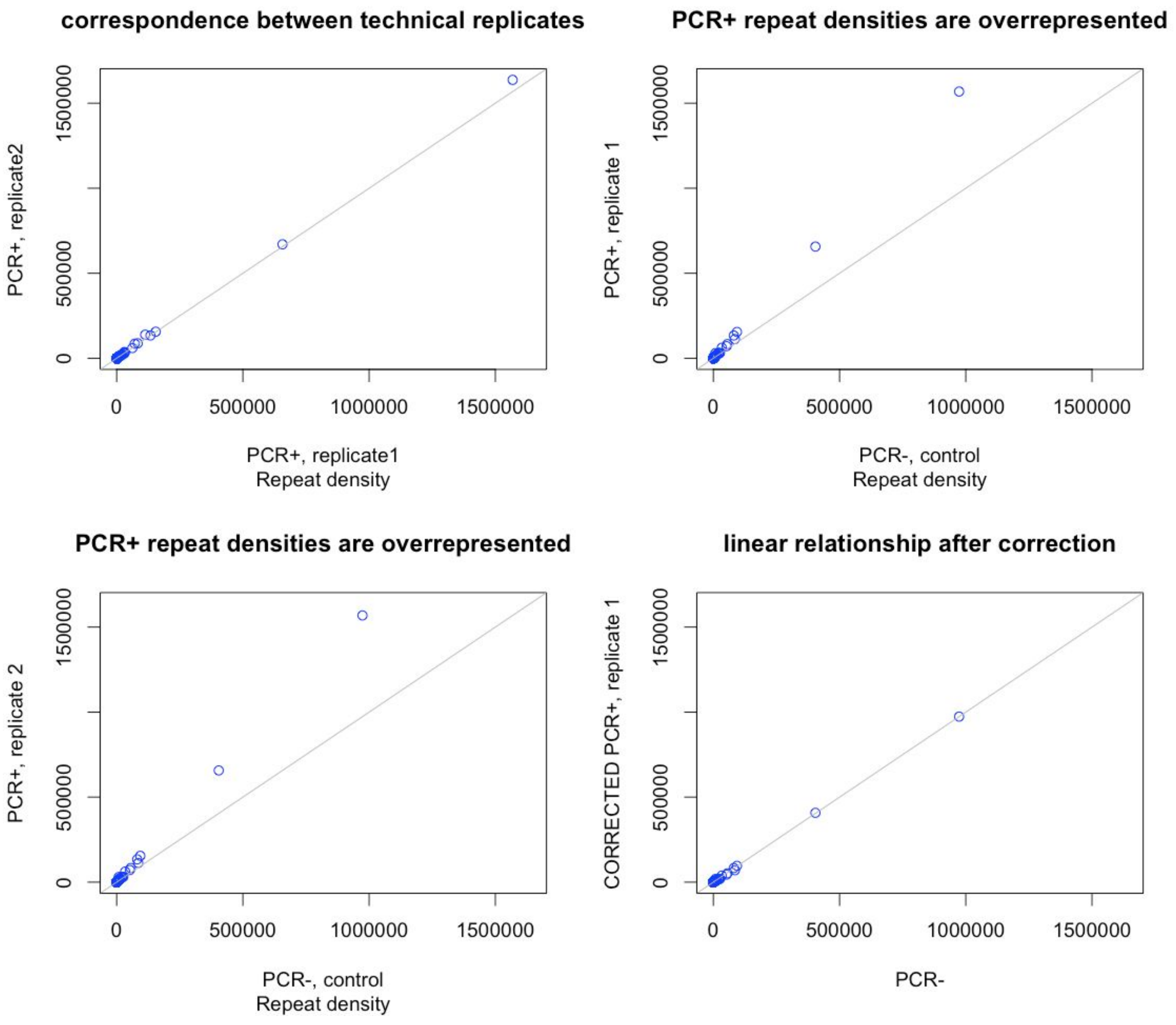
correspondence between technical replicates
PCR+, replicate2
PCR+, replicate1
Repeat density
PCR+ repeat densities are overrepresented
PCR+, replicate 1
PCR-, control
Repeat density
PCR+ repeat densities are overrepresented
PCR+, replicate 2
PCR-, control
Repeat density
linear relationship after correction
CORRECTED PCR+, replicate 1
PCR-
0
500000
1000000
1500000

#### Supplementary Note 3. Running NCRF

The following settings were used to run NCRF:

```
--scoring=nanopore --stats=events --positionalevents --maxnoise=20% --minlength=75
--scoring=pacbio --stats=events --positionalevents --maxnoise=20% --minlength=75
```

Filtering step:

```
ncrf_consensus_filter.py --winner=0.5
```

#### SUPPLEMENTARY FIGURES

**Figure S1. The repeat densities of technical replicates are tightly correlated.** The package ICCbare was used to calculate intraclass correlation coefficients or the "tightness" of the technical replicates (as some great ape individuals were sequenced as part of multiple runs). The number of technical replicates is listed in Table S1.

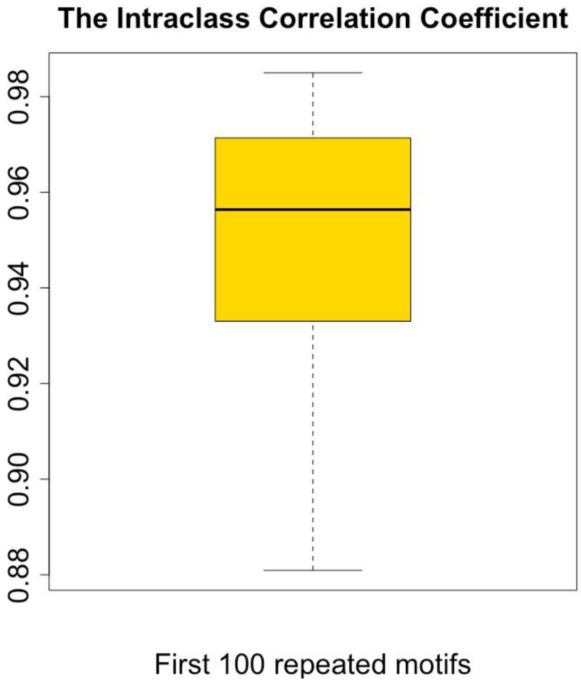

**Figure S2. Abundance of repeats in a sample is independent of the total number of reads in a sample.**

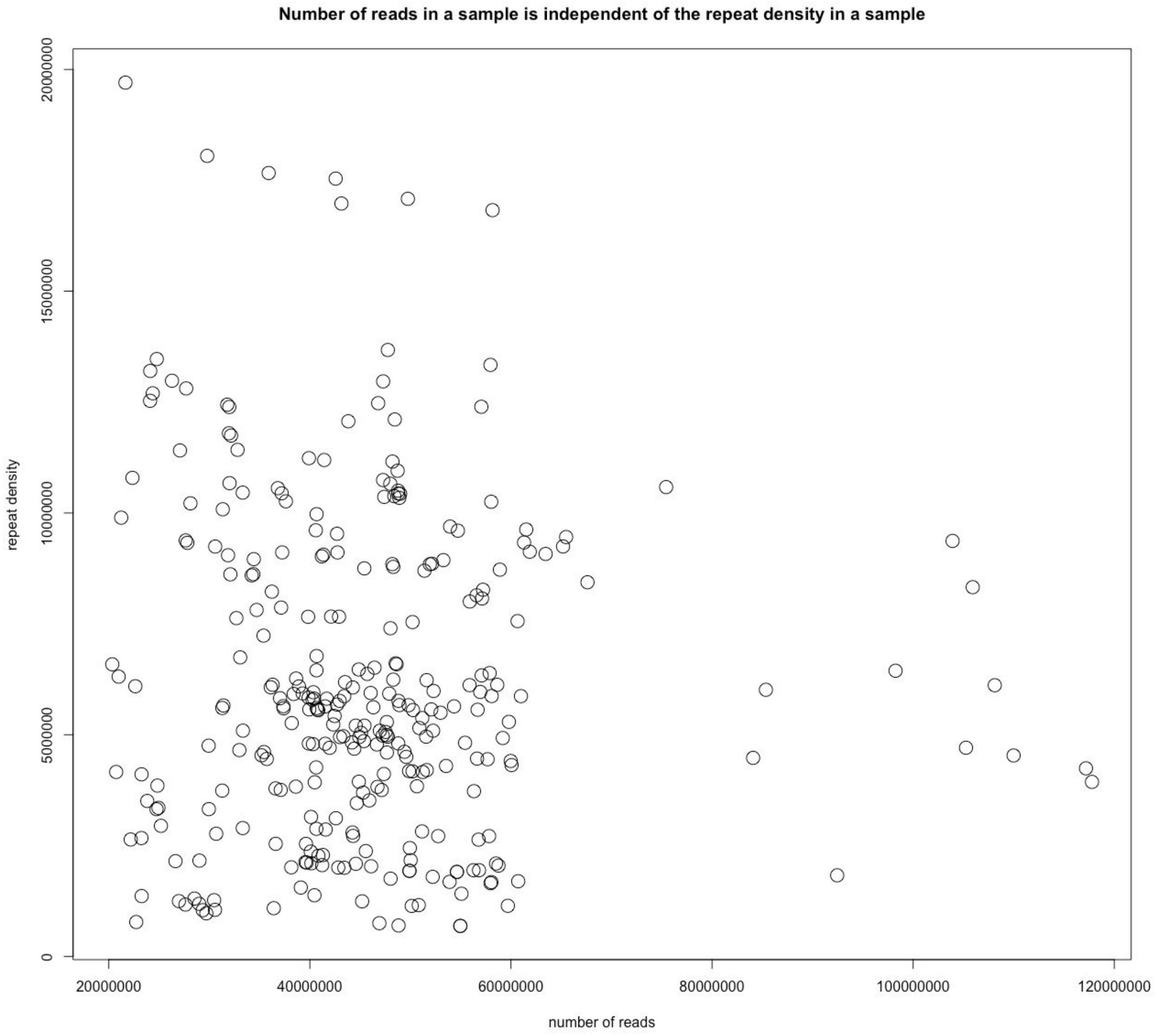

**Figure S3. 50 most abundant repeated motifs for each species.** Each repeated motif is plotted as a boxplot using repeat density (kb/Mb) from all individuals in a given species. The vertical red line separates the first 12 most abundant repeated motifs. Telomeric repeat is shown with an arrow when it is one of 50 most abundant repeats. **(A)** Human, **(B)** Chimpanzee, **(C)** Bonobo, **(D)** gorilla, **(E)** Sumatran orangutan, **(F)** Bornean orangutan.

**A**

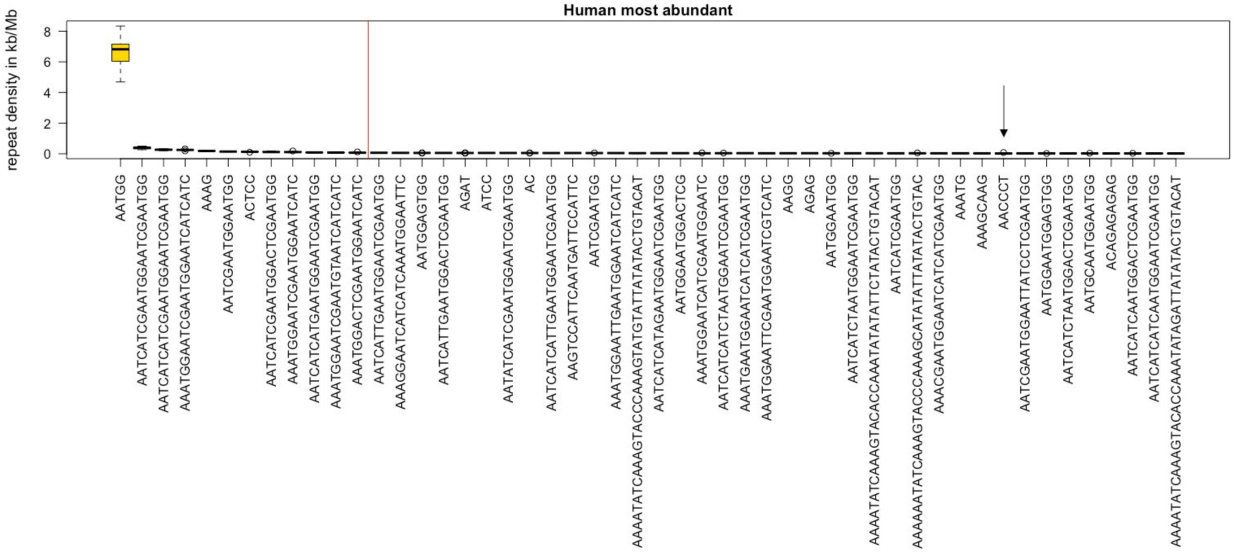


**B**

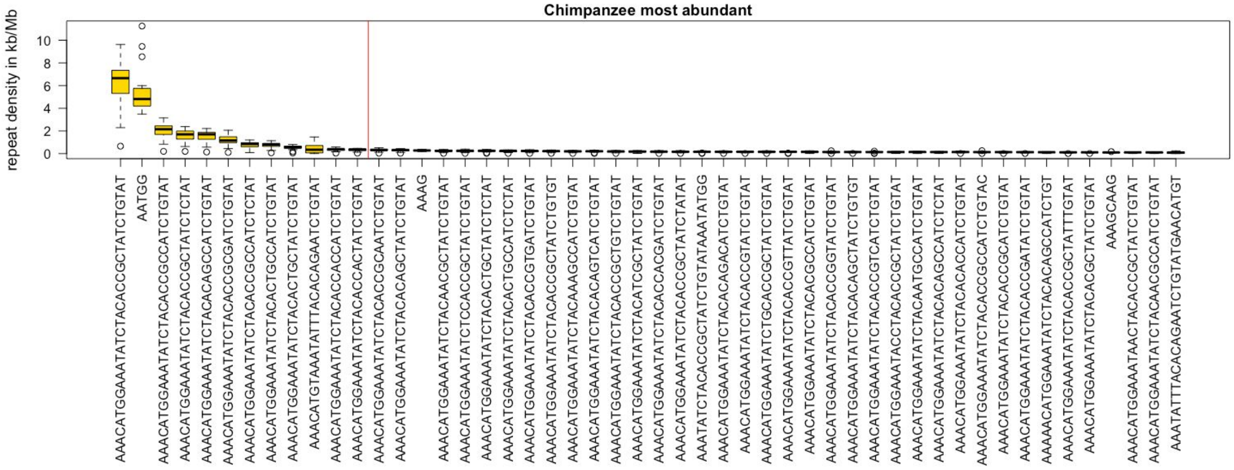

**C**

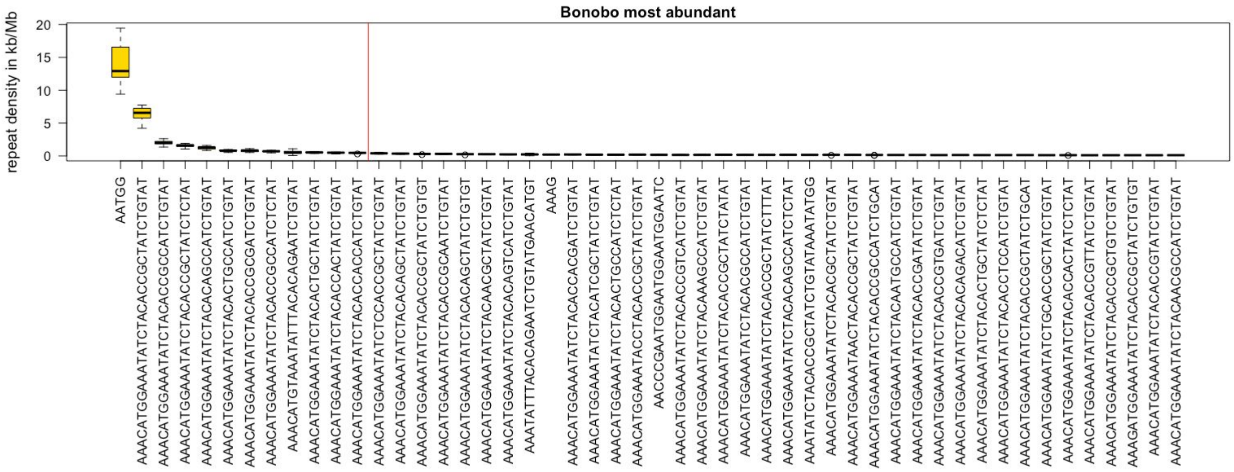
Bonobo most abundant
repeat density in kb/Mb

**D**

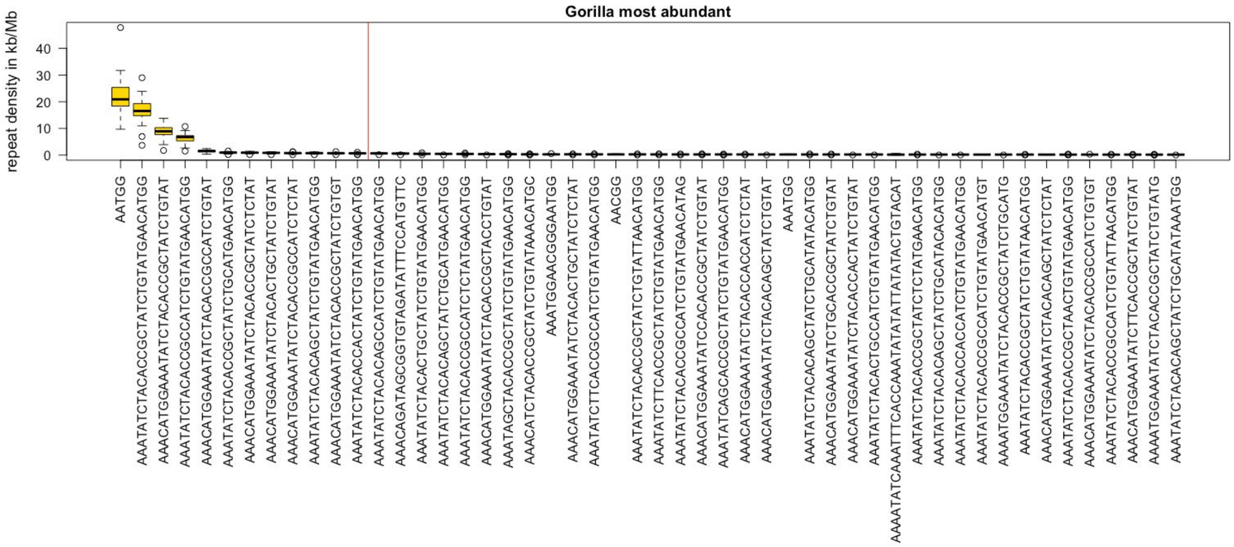
Gorilla most abundant
repeat density in kb/Mb

**E**

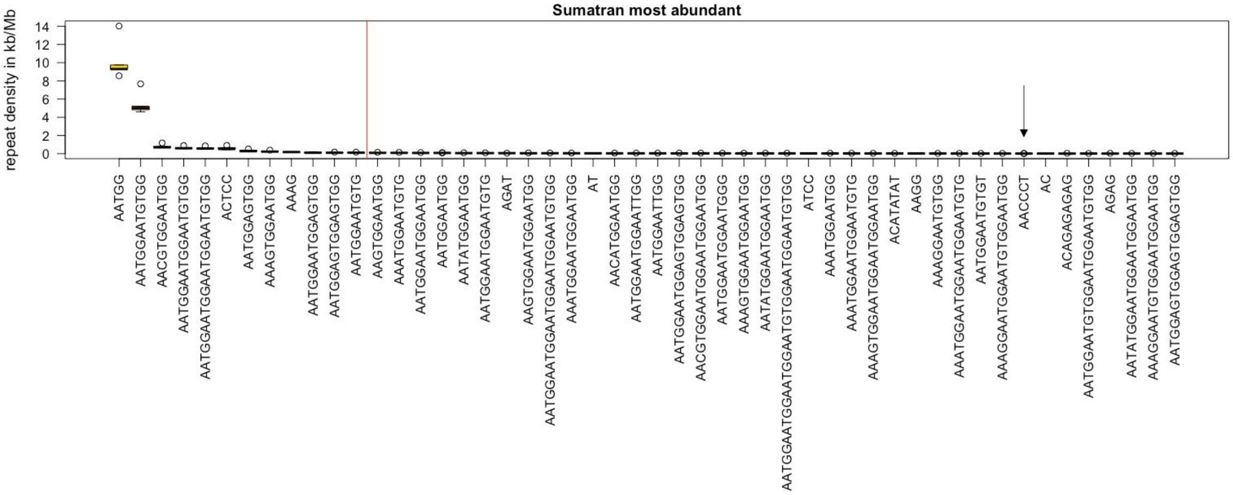

Sumatran most abundant
repeat density in kb/Mb
0
2
4
6
8
10
12
14
AATGG
AATGGAATGTGG
AACGTGGAATGG
AATGGAATGGAATGTGG
AATGGAATGGAATGGAATGTGG
ACTCC
AATGGAGTGG
AAAGTGGAATGG
AAAG
AATGGAATGGAGTGG
AATGGAGTGGAGTGG
AATGGAATGTG
AAGTGGAATGG
AAATGGAATGTG
AATGGAATGGAATGG
AATGGAATGG
AATATGGAATGG
AATGGAATGGAATGTG
AGAT
AAGTGGAATGGAATGG
AATGGAATGGAATGGAATGTGG
AAATGGAATGGAATGG
AT
AACATGGAATGG
AATGGAATGGAATTGG
AATGGAATTGG
AATGGAATGGAGTGGAGTGG
AACGTGGAATGGAATGGAATGG
AATGGAATGGAATGGG
AAAGTGGAATGGAATGG
AATATGGAATGGAATGG
AATGGAATGGAATGTGGAATGGAATGTGG
ATCC
AAATGGAATGG
AAATGGAATGGAATGTG
AAAGTGGAATGGAATGGAATGG
ACATATAT
AAGG
AAAGGAATGTGG
AAATGGAATGGAATGGAATGTG
AATGGAATGTGT
AAAGGAATGGAATGTGGAATGG
AACCCT
AC
ACAGAGAGAG
AATGGAATGTGGAATGGAATGTGG
AGAG
AATATGGAATGGAATGG
AAAGGAATGTGGAATGGAATGG
AATGGAGTGGAGTGG


**F**

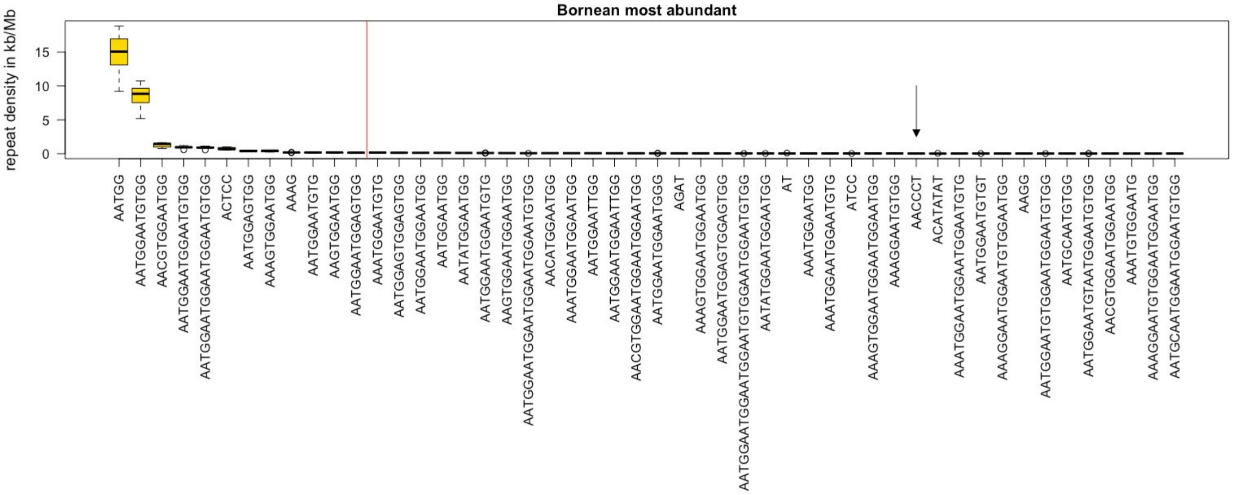

Bornean most abundant
repeat density in kb/Mb
0
5
10
15
AATGG
AATGGAATGTGG
AACGTGGAATGG
AATGGAATGGAATGTGG
AATGGAATGGAATGGAATGTGG
ACTCC
AATGGAGTGG
AAAGTGGAATGG
AAAG
AATGGAATGTG
AAGTGGAATGG
AATGGAATGGAGTGG
AAATGGAATGTG
AATGGAGTGGAGTGG
AATGGAATGGAATGG
AATGGAATGG
AATATGGAATGG
AATGGAATGGAATGTG
AAGTGGAATGGAATGG
AATGGAATGGAATGGAATGTGG
AACATGGAATGG
AAATGGAATGGAATGG
AATGGAATTGG
AATGGAATGGAATTGG
AACGTGGAATGGAATGGAATGG
AATGGAATGGAATGGG
AGAT
AAAGTGGAATGGAATGG
AATGGAATGGAGTGGAGTGG
AATGGAATGGAATGTGGAATGGAATGTGG
AATATGGAATGGAATGG
AT
AAATGGAATGG
AAATGGAATGGAATGTG
ATCC
AAAGTGGAATGGAATGGAATGG
AAAGGAATGTGG
AACCCT
ACATATAT
AAATGGAATGGAATGGAATGTG
AATGGAATGTGT
AAAGGAATGGAATGTGGAATGG
AAGG
AATGGAATGTGGAATGGAATGTGG
AATGCAATGTGG
AATGGAATGTAATGGAATGTGG
AACGTGGAATGGAATGG
AAATGTGGAATG
AAAGGAATGTGGAATGGAATGG
AATGCAATGGAATGGAATGTGG

**Figure S4. Analysis of repeat density in three human trios.** Repeat densities in kb/Mb from trio data (repeated motifs absent in human are not shown).

| | Father 77 | Father 78 | Father HG | Son 77 | Son HG | Mother 77 | Mother 78 | Mother HG | Daughter 78 |
|---|---|---|---|---|---|---|---|---|---|
| (1) AATGG | 4.22 | 4.21 | 6.1 | 4.21 | 5.76 | 3.36 | 3.7 | 5.17 | 3.71 |
| (2) ACTCC | 0.08 | 0.09 | 0.12 | 0.09 | 0.11 | 0.06 | 0.08 | 0.11 | 0.09 |
| (3) AAAG | 0.17 | 0.18 | 0.18 | 0.17 | 0.19 | 0.17 | 0.19 | 0.19 | 0.2 |
| (4) AATGGAGTGG | 0.03 | 0.04 | 0.05 | 0.04 | 0.05 | 0.03 | 0.03 | 0.05 | 0.04 |
| (5) AATGGAATGGAGTGG | 0.02 | 0.02 | 0.03 | 0.02 | 0.02 | 0.01 | 0.02 | 0.02 | 0.02 |
| (6) AATGGAGTGGAGTGG | 0.01 | 0.01 | 0.02 | 0.01 | 0.02 | 0.01 | 0.01 | 0.02 | 0.01 |
| (14) AAATGGACTCGAATGGAATCATC | 0.06 | 0.07 | 0.1 | 0.05 | 0.06 | 0.03 | 0.03 | 0.05 | 0.06 |
| (15) AATCGAATGGAATGG | 0.11 | 0.12 | 0.18 | 0.1 | 0.17 | 0.03 | 0.05 | 0.07 | 0.06 |
| (16) AAATGGAATCGAATGGAATCATC | 0.09 | 0.1 | 0.14 | 0.08 | 0.09 | 0.05 | 0.05 | 0.08 | 0.08 |
| (17) AATCATCGAATGGAATCGAATGG | 0.27 | 0.36 | 0.48 | 0.25 | 0.35 | 0.23 | 0.27 | 0.34 | 0.3 |
| (18) AATCATCGAATGGACTCGAATGG | 0.11 | 0.14 | 0.18 | 0.1 | 0.12 | 0.07 | 0.08 | 0.11 | 0.11 |
| (19) AATCATCATGAATGGAATCGAATGG | 0.05 | 0.06 | 0.08 | 0.05 | 0.06 | 0.04 | 0.05 | 0.06 | 0.05 |
| (20) AAATGGAATCGAATGGAATCATCATC | 0.16 | 0.21 | 0.26 | 0.16 | 0.21 | 0.19 | 0.21 | 0.22 | 0.19 |
| (21) AAATGGAATCGAATGTAATCATCATC | 0.04 | 0.06 | 0.08 | 0.05 | 0.06 | 0.06 | 0.06 | 0.07 | 0.05 |
| (22) AATCATCATCGAATGGAATCGAATGG | 0.21 | 0.26 | 0.36 | 0.2 | 0.26 | 0.18 | 0.2 | 0.25 | 0.22 |
| Cumulative density of abundant repeats | 5.64 | 5.94 | 8.37 | 5.58 | 7.55 | 4.53 | 5.03 | 6.80 | 5.18 |
| Cumulative density of all repeats | 8.94 | 9.62 | 13.30 | 9.31 | 11.93 | 7.71 | 8.47 | 10.99 | 8.77 |

**Figure S5. Similarity and inter-relatedness among the sequences of 39 abundant repeated motifs.** Each circle represents a repeat (indexes inside the circles match those in Fig. 1B). The sizes of circles represent four categories of repeat densities from smallest to largest: 0-0.1 kb/Mb, 0.1-1 kb/Mb, 1-10 kb/Mb, and >10 kb/Mb. The color of a link represents the number of substitutions (see Methods) needed for the shorter repeated motif to perfectly match the longer related repeated motif (yellow: one step; orange: two steps; red: three steps). The circles depicting repeated motifs related to 32-mers are filled in green; the ones corresponding to all other repeated motifs (usually related to $(AATGG)_n$) are filled in yellow.

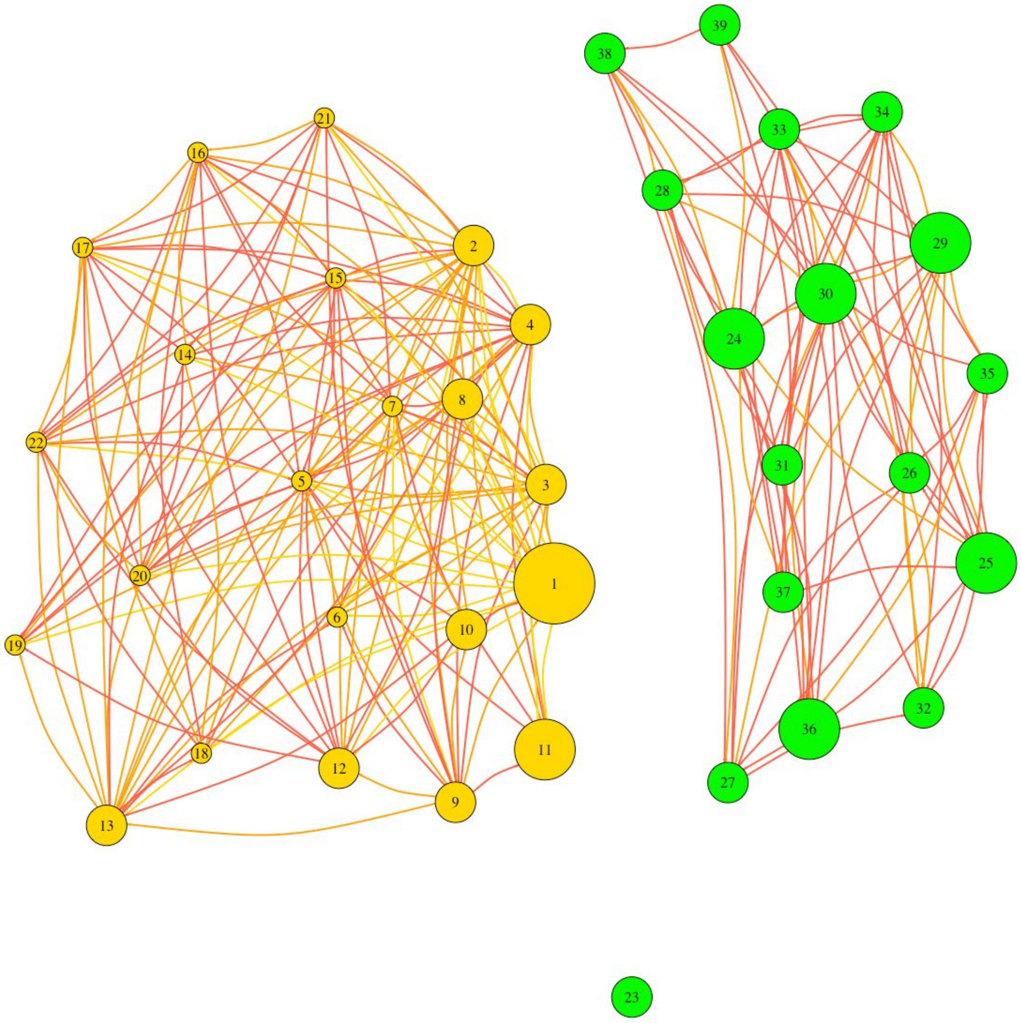

**Figure S6. The 39 most abundant repeated motifs and their corresponding repeat densities.** Repeated motifs present in all six species are shown in black, repeated motifs present in a subset of species are shown in black. The order of repeated motifs matches their order in Fig. 1B.

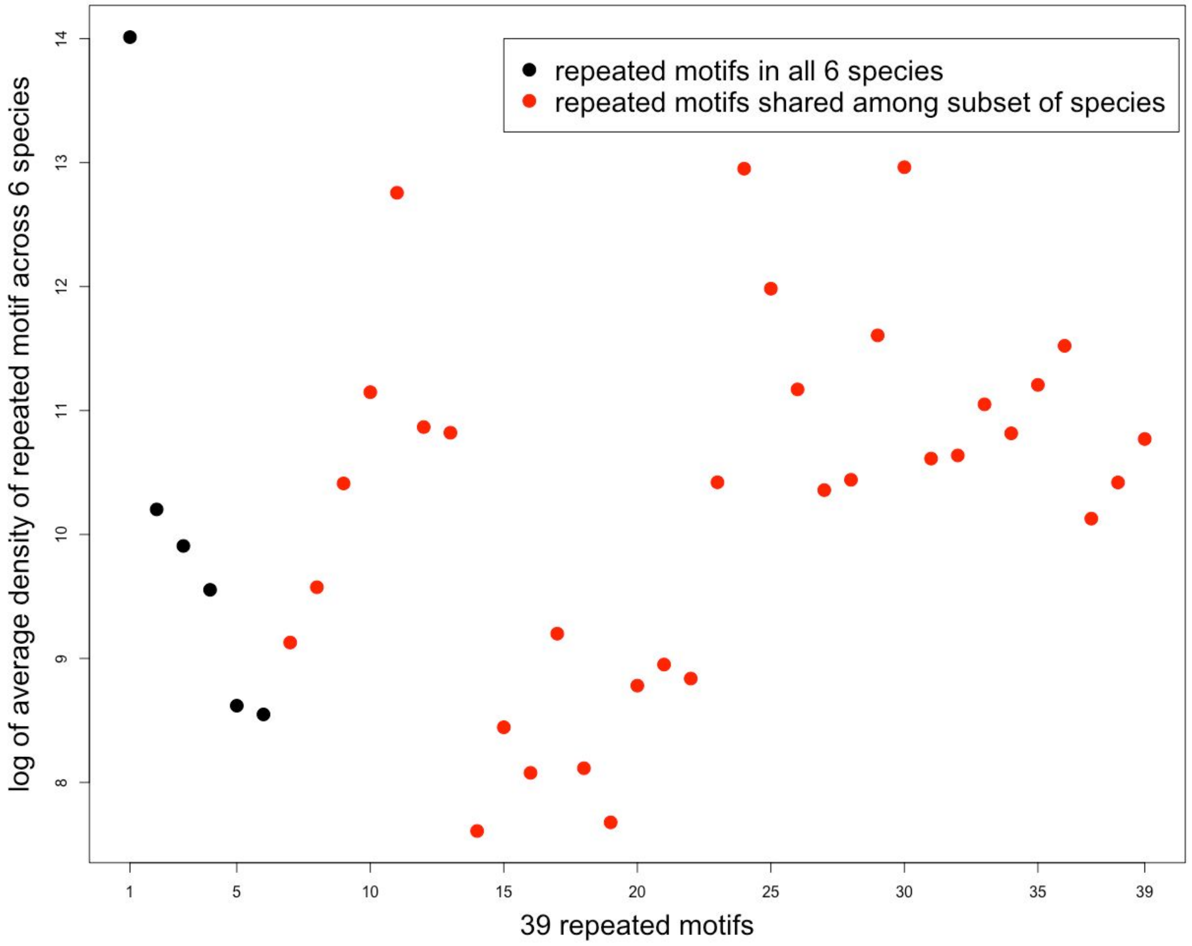

**Figure S7. Substantial differences in repeat presence/absence and in repeat density were observed among individuals.** Color coding from dark to light red represents high to low values. Note that the darkest red marks cases in which a repeat is present in all individuals of a species. **(A)** Heatmaps of proportions of individuals carrying 39 abundant repeats in each of the six species. **(B)** Heatmaps of proportions of individuals carrying remaining (less abundant) 5,455 identified repeats in each of the six species. Among abundant repeats present in each species, all (15 out of a total of 15) were present in all human individuals, and most (24 out of 31, 24 out of 33, 13 out of 19, and 13 out of 17) were present in all bonobo, gorilla, Sumatran orangutan, and Bornean orangutan individuals, respectively, but only 20 out of 33 were present in all chimpanzee individuals. The trend was the opposite for less abundant repeated motifs which often were not shared among all analyzed individuals within a species. **(C) - (H).** Same heatmap as in (B), using the data randomly subsampled to only 5 individuals for each species to match the number of individuals resequenced for each of the two orangutan species.

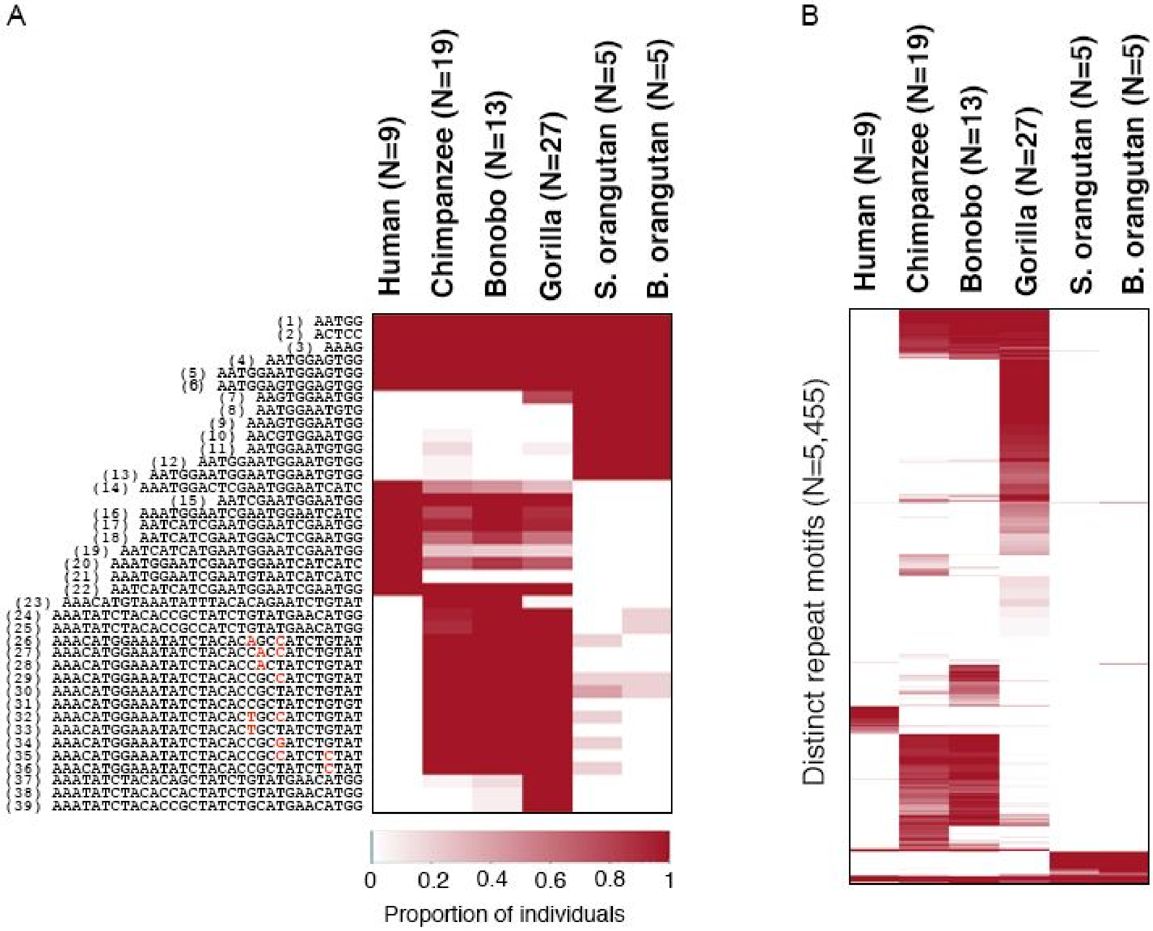

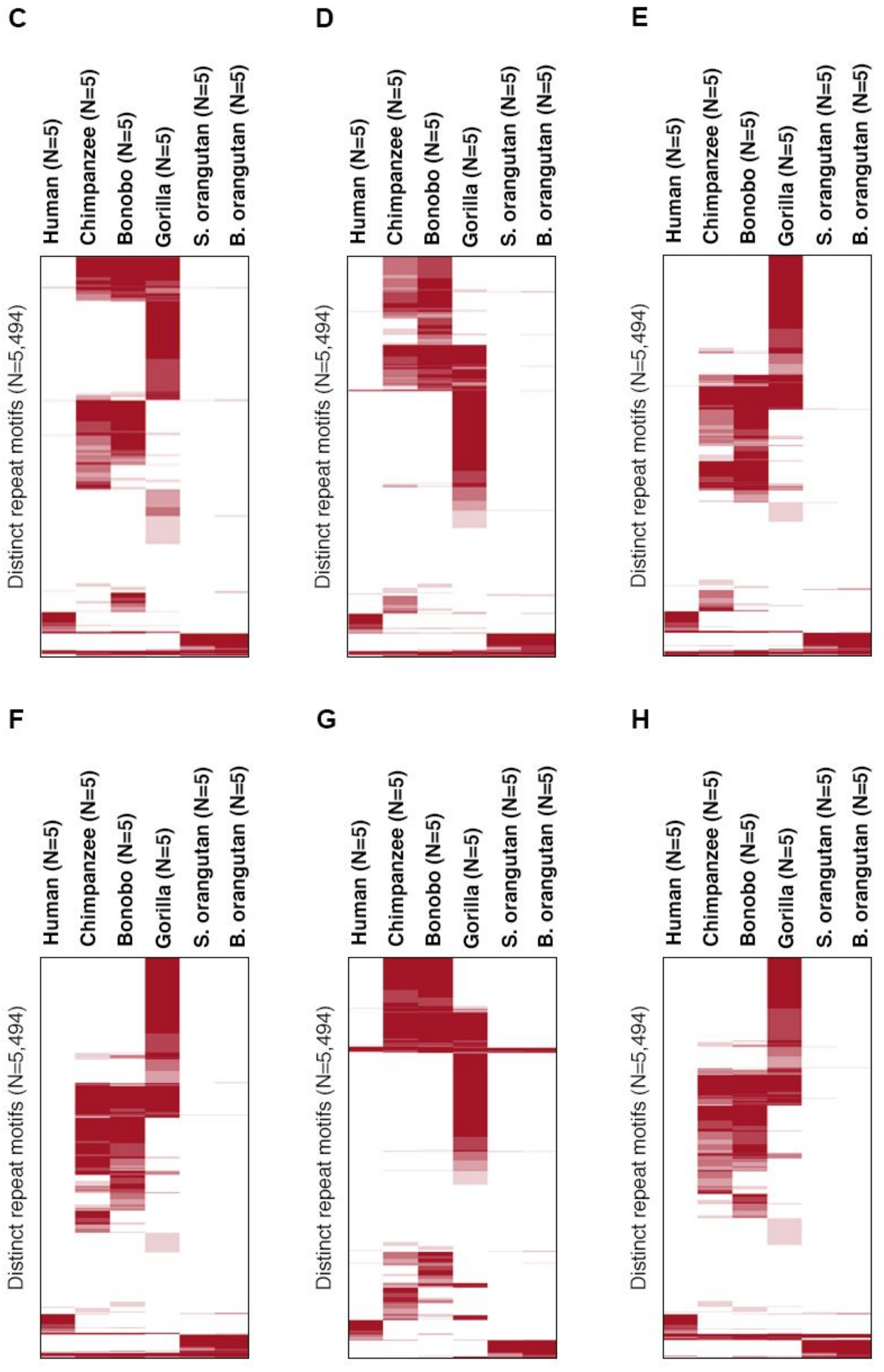
C
D
E
F
G
H
Human (N=5)
Chimpanzee (N=5)
Bonobo (N=5)
Gorilla (N=5)
S. orangutan (N=5)
B. orangutan (N=5)
Distinct repeat motifs (N=5,494)

**Figure S8. (A) A positive relationship between mean repeat density for a species and the number of species-specific repeats, with humans as an outlier.**

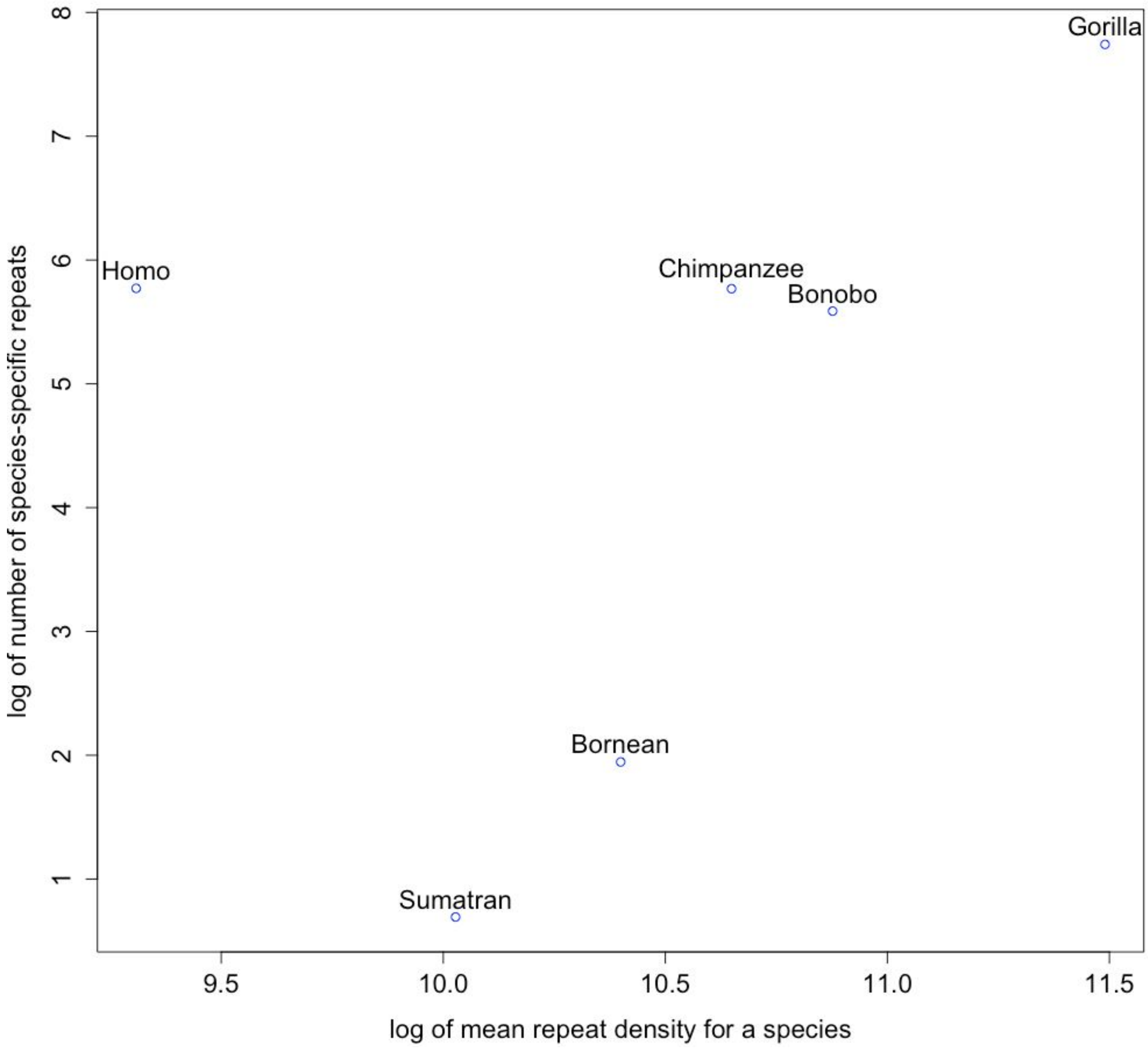

**(B) The relationship between mean repeat density for a species and the number of species-specific repeats is robust to the subsampling.** We randomly subsampled five individuals per species, equal to the number of individuals resequenced for each of the two orangutan species.

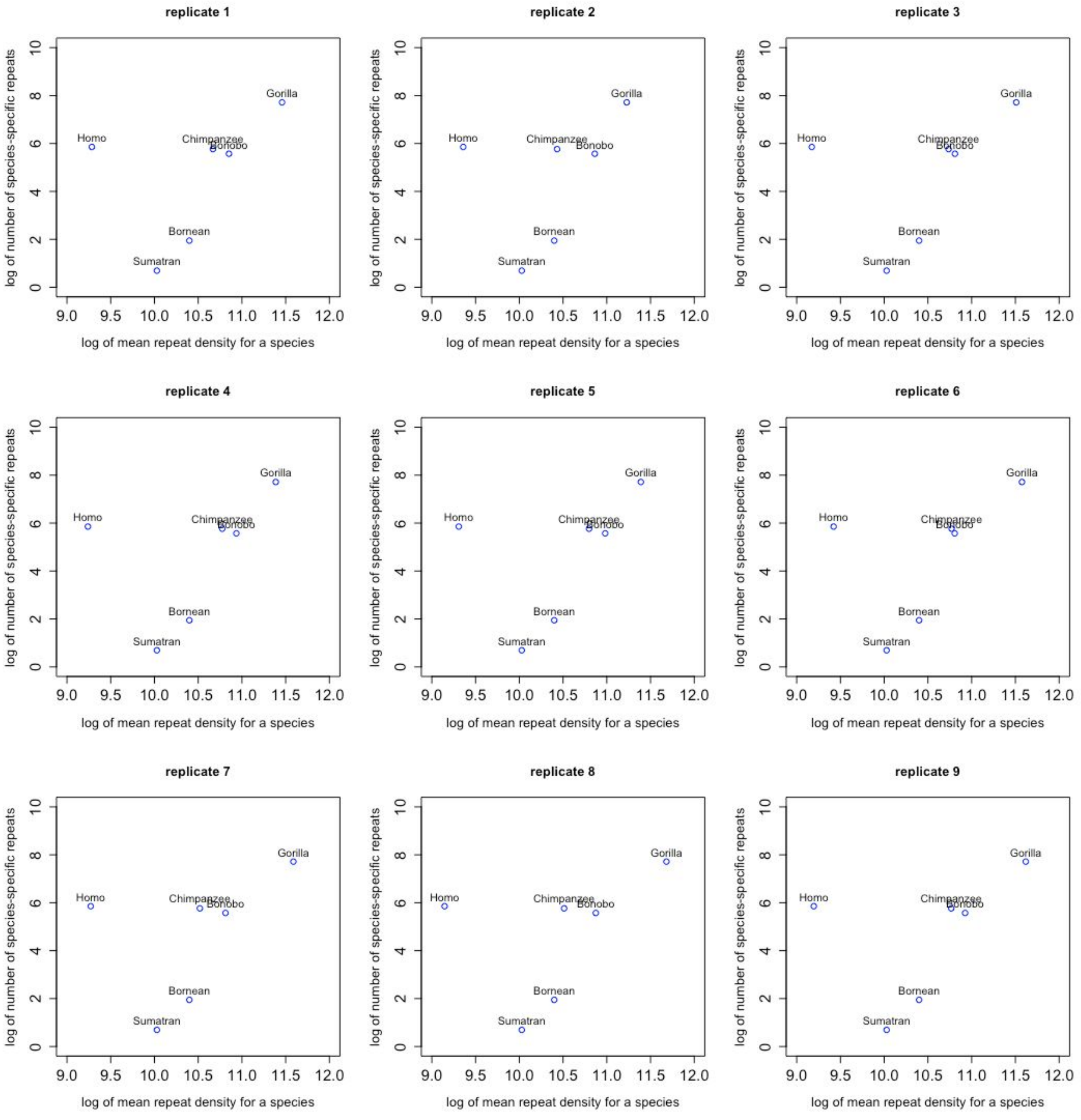

**Figure S9. Substantial differences exist among individuals, in repeat presence/absence as well as density.** Heatmap of average repeat densities for the 39 abundant repeats (+ the telomeric repeat) in the individuals of each of the six species. Color coding from dark to light blue represents high to low values (see color legend at the bottom). Repeats present at less than 100 loci per 20 million reads are considered absent (white cells). $(AATGG)_n$-derived and 32-mer-derived repeated motifs are separated by a horizontal line.

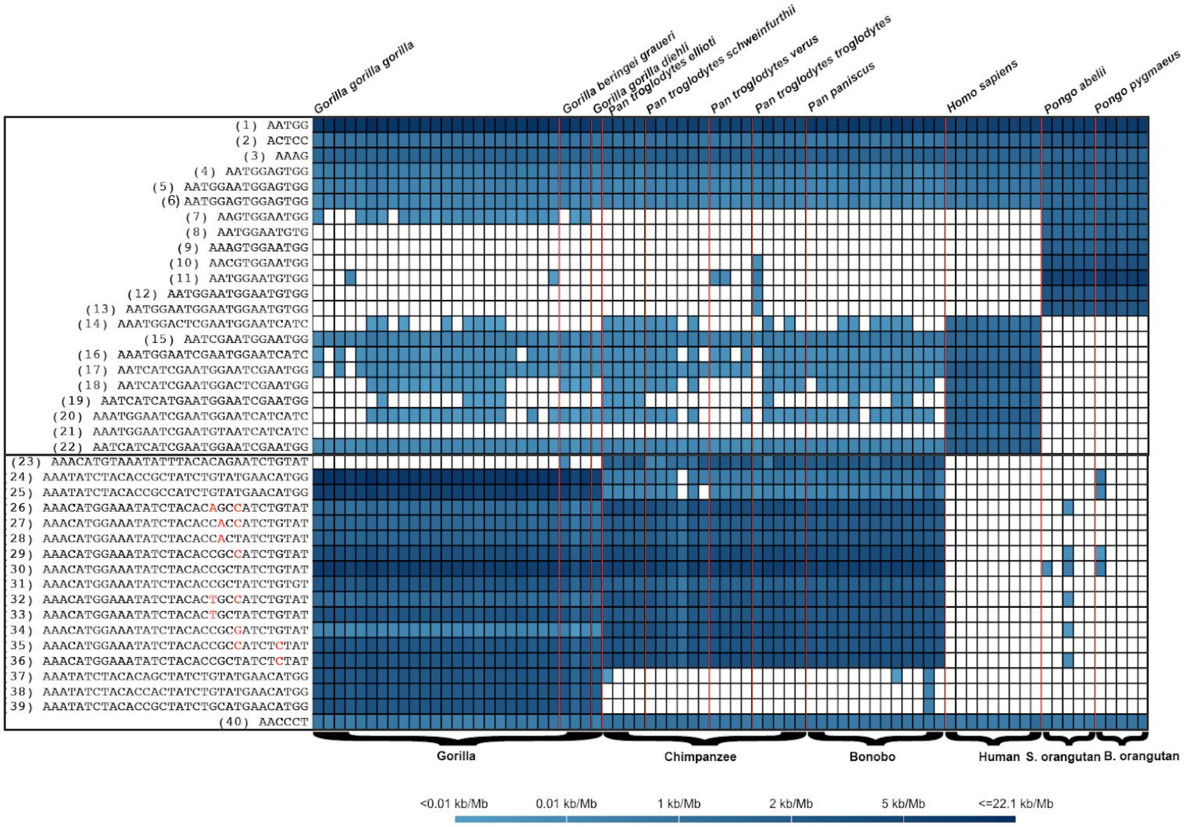

**Figure S10. Hierarchical clustering does not reproduce the accepted species phylogeny.** Human (black), bonobo (blue), chimpanzee (green), gorilla (red) and orangutan (orange) are clustered using hierarchical clustering. The distance metrics is computed as one minus correlation coefficient between repeat counts of two individuals. **(A)** Using all 5,494 repeated motifs and Pearson correlation coefficient, complete linkage. **(B)** Using all 5,494 repeated motifs and Pearson correlation coefficient, single linkage. **(C)** Using the 39 most abundant repeated motifs and Pearson correlation coefficient, complete linkage. **(D)** Using the 39 most abundant repeated motifs and Pearson correlation coefficient, single linkage. **(E)** Using all 5,494 repeated motifs and Spearman correlation coefficient, complete linkage. **(F)** Using all 5,494 repeated motifs and Spearman correlation coefficient, single linkage. **(G)** Using the 39 most abundant repeated motifs and Spearman correlation coefficient, complete linkage. **(H)** Using the 39 most abundant repeated motifs and Spearman correlation coefficient, single linkage. **(I)** Excluding StSat and using 3,043 repeated motifs and Pearson correlation coefficient, complete linkage.

**A.** Using all 5,494 repeated motifs and Pearson correlation coefficient, complete linkage.

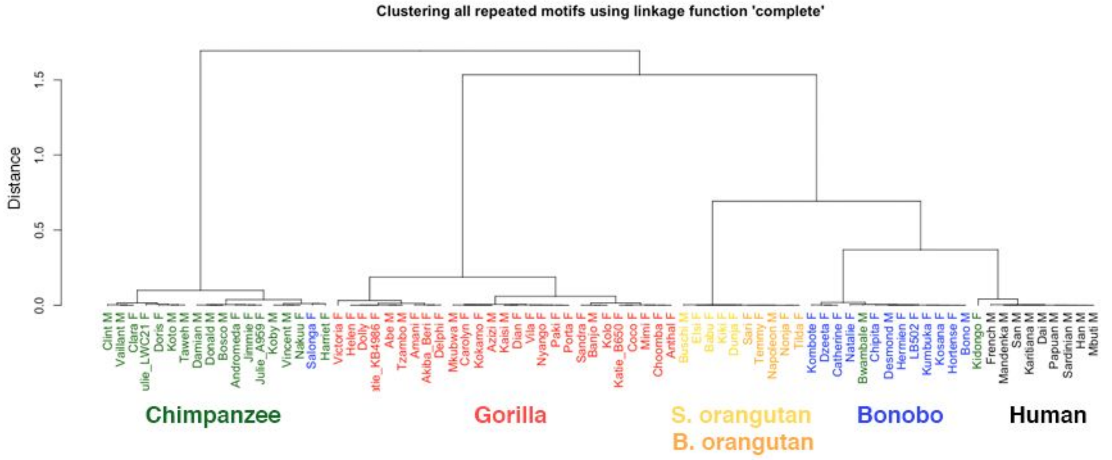


**B.** Using all 5,494 repeated motifs and Pearson correlation coefficient, single linkage.

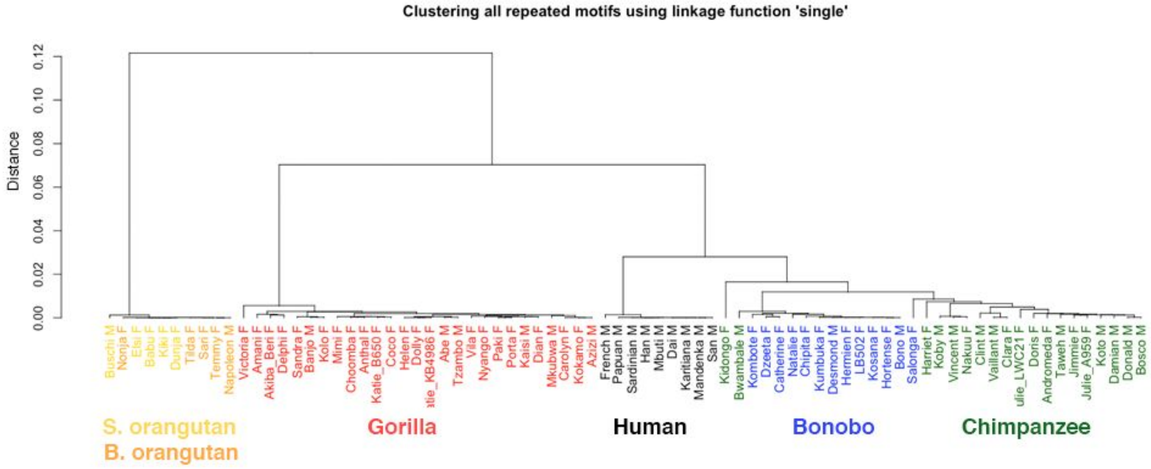

**C**. Using the 39 most abundant repeated motifs and Pearson correlation coefficient, complete linkage.

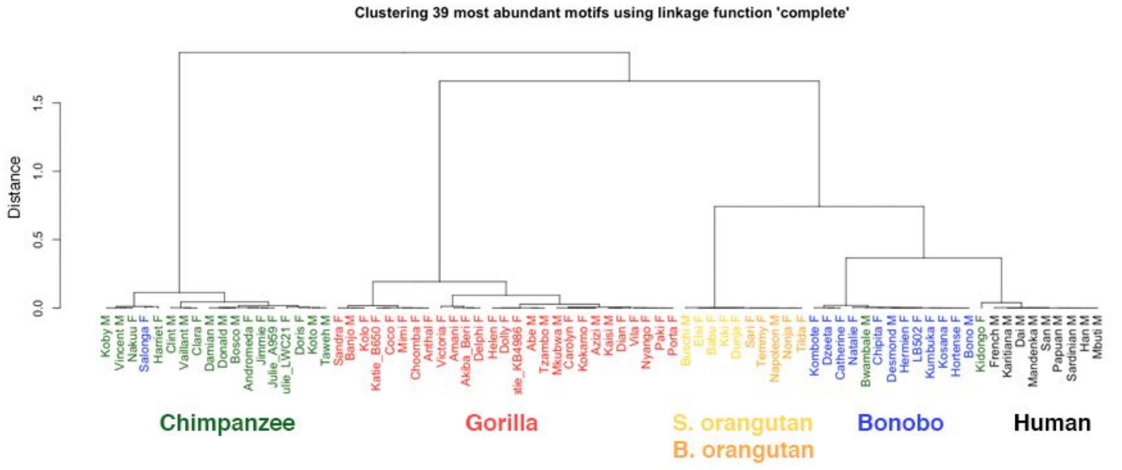


**D**. Using the 39 most abundant repeated motifs and Pearson correlation coefficient, single linkage.

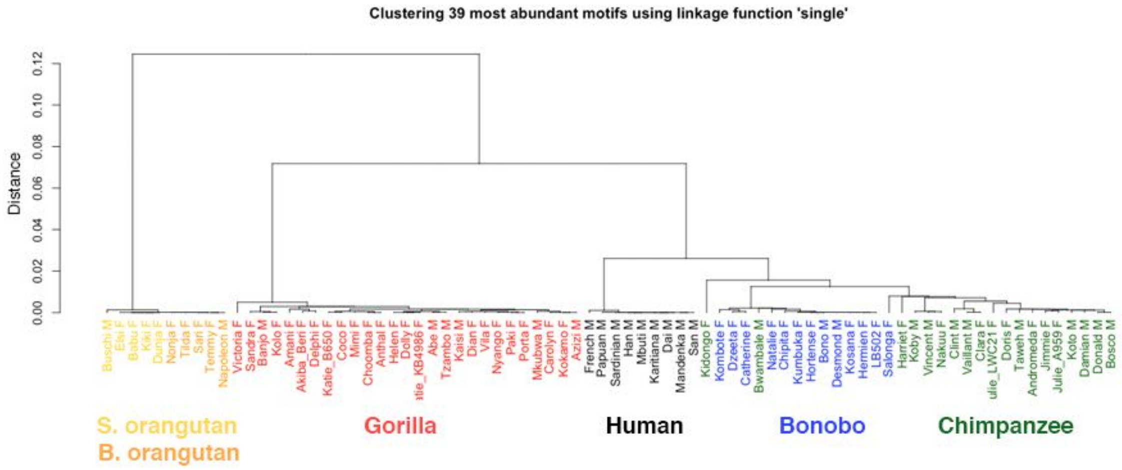


**E**. Using all 5,494 repeated motifs and Spearman correlation coefficient, complete linkage.

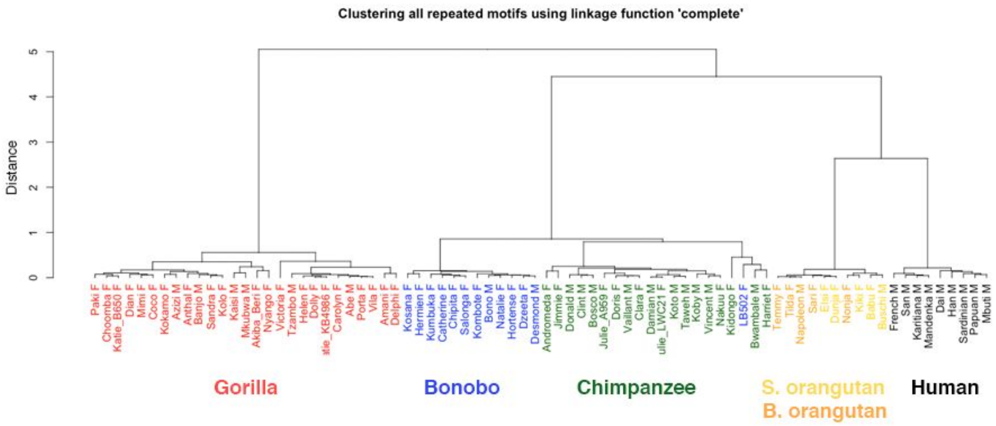

**F**. Using all 5,494 repeated motifs and Spearman correlation coefficient, single linkage.

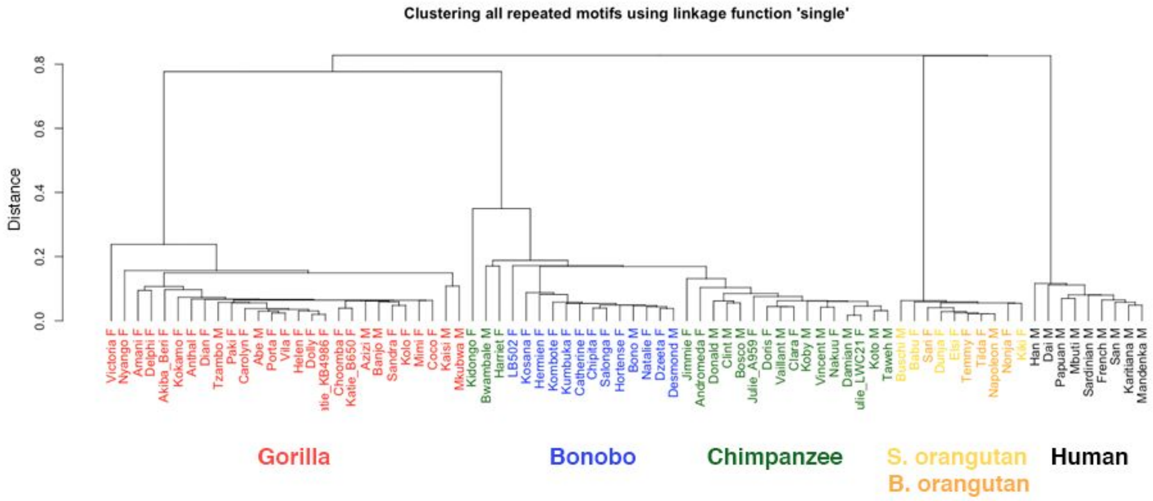


**G**. Using the 39 most abundant repeated motifs and Spearman correlation coefficient, complete linkage.

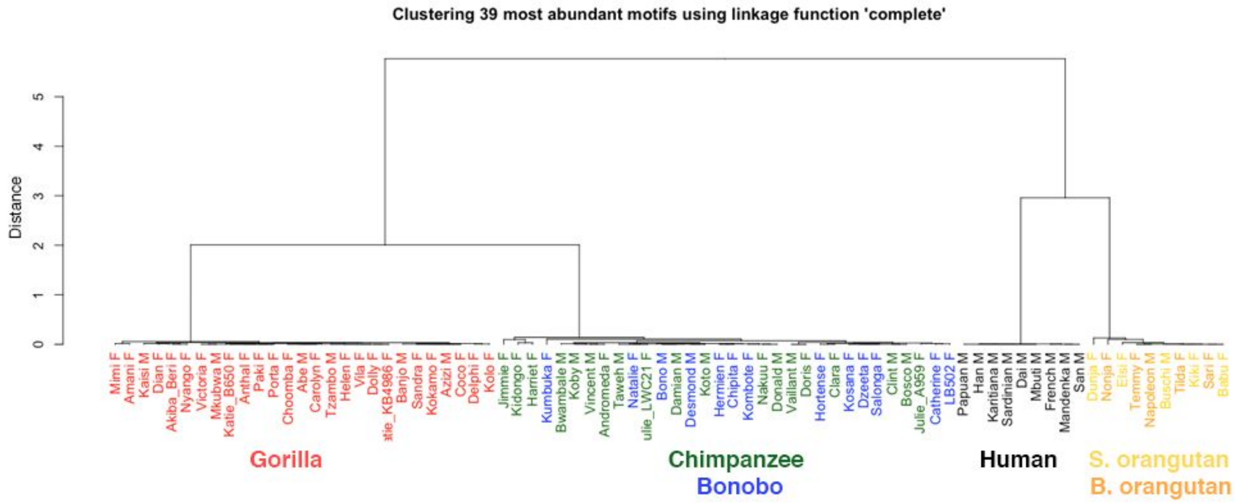

**H**. Using the 39 most abundant repeated motifs and Spearman correlation coefficient, single linkage.

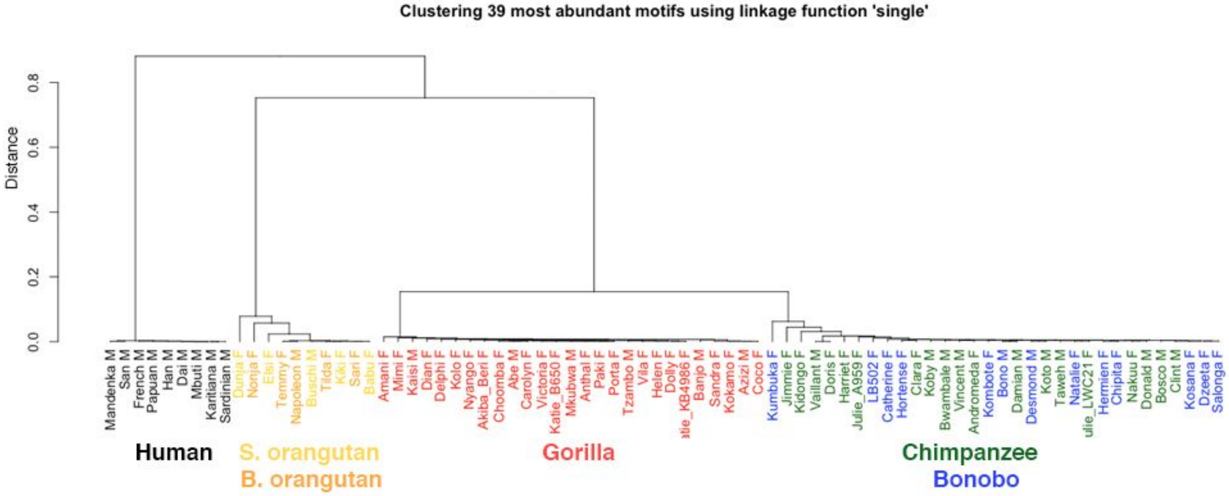


**I**

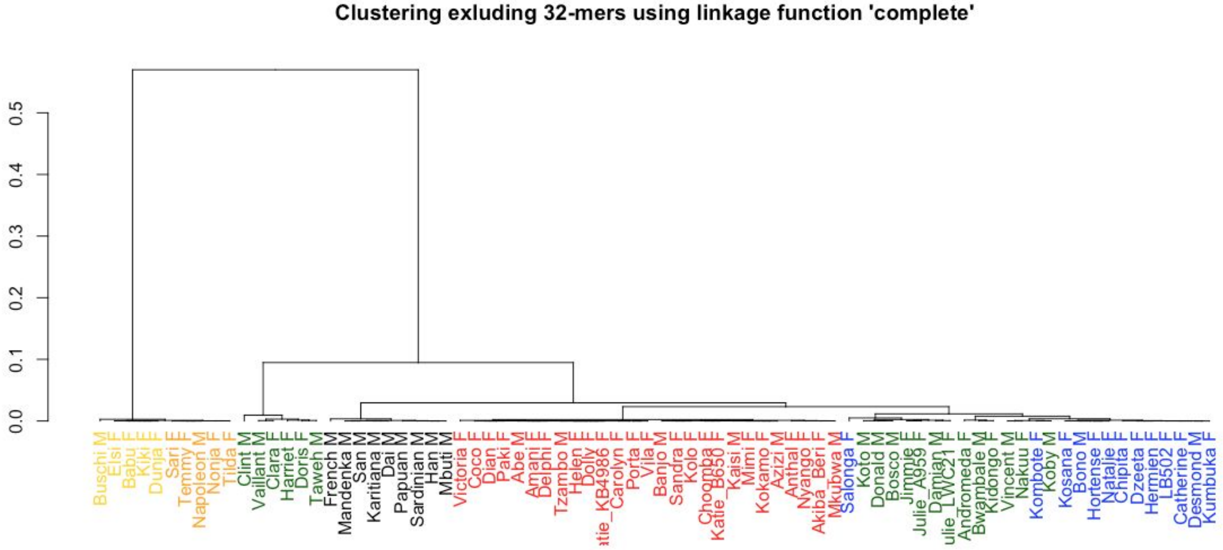

**Figure S11. Species topology based on repeats presence/absence.** A schematic figure showing repeated motifs unique to a species (terminal branches) and those that are shared among the species descending from internal branches. On the left, the tree is built based on the presence/absence of repeated motifs, after excluding StSat repeated motifs (here identified as those with unit sizes of 31-32 bp), iteratively joining species sharing the most repeated motifs. On the right, the tree is built according to the accepted species phylogeny (Goodman et al. 2005) and the number of shared repeated motifs is indicated. The branch widths are proportional to the number of repeated motifs (branch lengths are uninformative). 72 repeated motifs (the number shown in the middle) were shared among all six studied species.

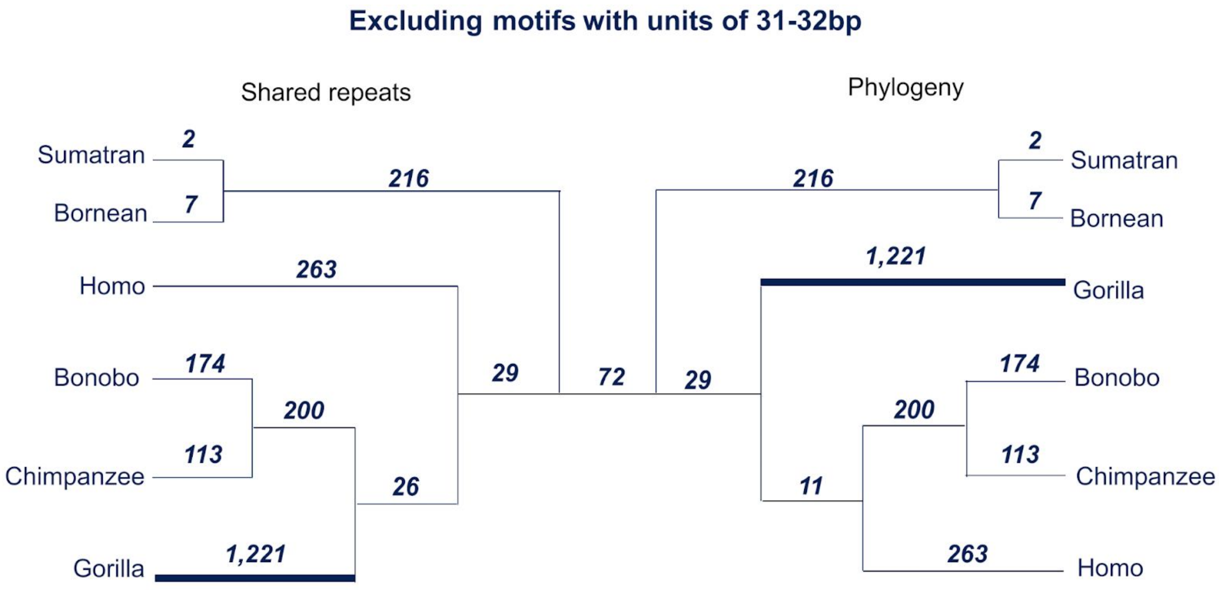

**Figure S12. Correlations in densities of 39 most abundant repeated motifs. Spearman correlation coefficients are shown.** Red and blue dots represent the observed correlation coefficients that were significantly different from the landscape of positive (red) and negative (blue) correlations, respectively, arising by chance -- these are shown in black (see Methods). The heatmap is showing pairwise correlation coefficients between the same repeated motifs. **(A)** Bonobo comparison against null expectations. **(B)** Bonobo correlation plot. **(C)** Chimpanzee comparison against null expectations. **(D)** Chimpanzee correlation plot. **(E)** Sumatran orangutan comparison against null expectations. **(F)** Sumatran orangutan correlation plot. **(G)** Bornean orangutan comparison against null expectations. **(H)** Bornean orangutan correlation plot.

**A**

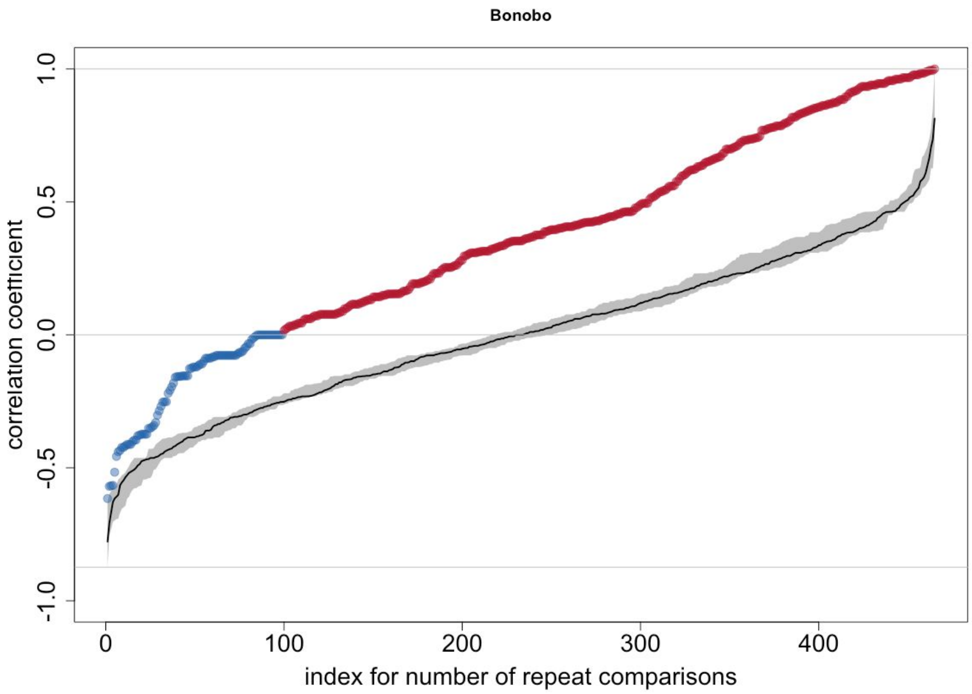
Bonobo
correlation coefficient
1.0
0.5
0.0
-0.5
-1.0
0
100
200
300
400
index for number of repeat comparisons

**B**

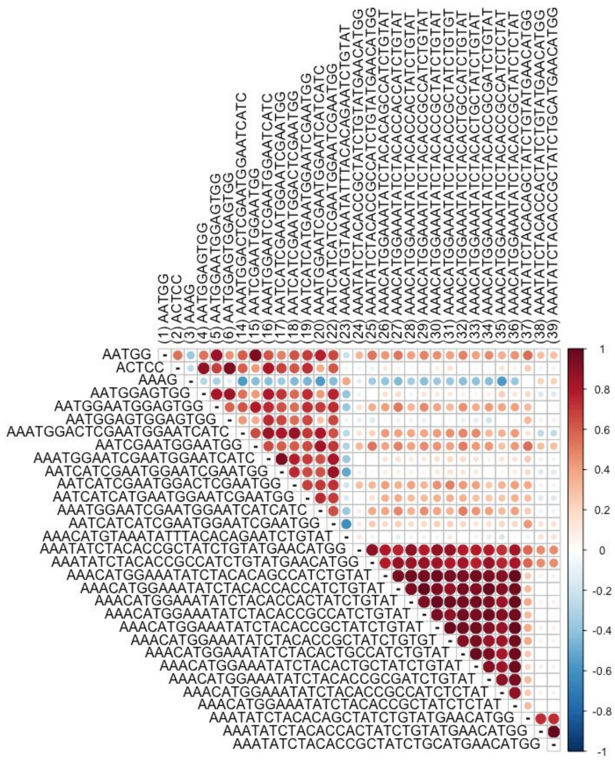
(1) AATGG
(2) ACTCC
(3) AAAG
(4) AATGGAGTGG
(5) AATGGAATGGAGTGG
(6) AATGGAGTGGAGTGG
(14) AAATGGACTCGAATGGAATCATC
(15) AATCGAATGGAATGG
(16) AAATGGAATCGAATGGAATCATC
(17) AATCATCGAATGGAATCGAATGG
(18) AATCATCGAATGGACTCGAATGG
(19) AATCATCATGAATGGAATCGAATGG
(20) AAATGGAATCGAATGGAATCATCATC
(22) AATCATCATCGAATGGAATCGAATGG
(23) AAACATGTAAATATTTACACAGAATCTGTAT
(24) AAATATCTACACCGCTATCTGTATGAACATGG
(25) AAATATCTACACCGCCATCTGTATGAACATGG
(26) AAACATGGAAATATCTACACAGCCATCTGTAT
(27) AAACATGGAAATATCTACACCACCATCTGTAT
(28) AAACATGGAAATATCTACACCACTATCTGTAT
(29) AAACATGGAAATATCTACACCGCCATCTGTAT
(30) AAACATGGAAATATCTACACCGCTATCTGTAT
(31) AAACATGGAAATATCTACACCGCTATCTGTGT
(32) AAACATGGAAATATCTACACTGCCATCTGTAT
(33) AAACATGGAAATATCTACACTGCTATCTGTAT
(34) AAACATGGAAATATCTACACCGCGATCTGTAT
(35) AAACATGGAAATATCTACACCGCCATCTCTAT
(36) AAACATGGAAATATCTACACCGCTATCTCTAT
(37) AAATATCTACACAGCTATCTGTATGAACATGG
(38) AAATATCTACACCACTATCTGTATGAACATGG
(39) AAATATCTACACCGCTATCTGCATGAACATGG
AATGG
ACTCC
AAAG
AATGGAGTGG
AATGGAATGGAGTGG
AATGGAGTGGAGTGG
AAATGGACTCGAATGGAATCATC
AATCGAATGGAATGG
AAATGGAATCGAATGGAATCATC
AATCATCGAATGGAATCGAATGG
AATCATCGAATGGACTCGAATGG
AATCATCATGAATGGAATCGAATGG
AAATGGAATCGAATGGAATCATCATC
AATCATCATCGAATGGAATCGAATGG
AAACATGTAAATATTTACACAGAATCTGTAT
AAATATCTACACCGCTATCTGTATGAACATGG
AAATATCTACACCGCCATCTGTATGAACATGG
AAACATGGAAATATCTACACAGCCATCTGTAT
AAACATGGAAATATCTACACCACCATCTGTAT
AAACATGGAAATATCTACACCACTATCTGTAT
AAACATGGAAATATCTACACCGCCATCTGTAT
AAACATGGAAATATCTACACCGCTATCTGTAT
AAACATGGAAATATCTACACCGCTATCTGTGT
AAACATGGAAATATCTACACTGCCATCTGTAT
AAACATGGAAATATCTACACTGCTATCTGTAT
AAACATGGAAATATCTACACCGCGATCTGTAT
AAACATGGAAATATCTACACCGCCATCTCTAT
AAACATGGAAATATCTACACCGCTATCTCTAT
AAATATCTACACAGCTATCTGTATGAACATGG
AAATATCTACACCACTATCTGTATGAACATGG
AAATATCTACACCGCTATCTGCATGAACATGG
1
0.8
0.6
0.4
0.2
0
-0.2
-0.4
-0.6
-0.8
-1

C

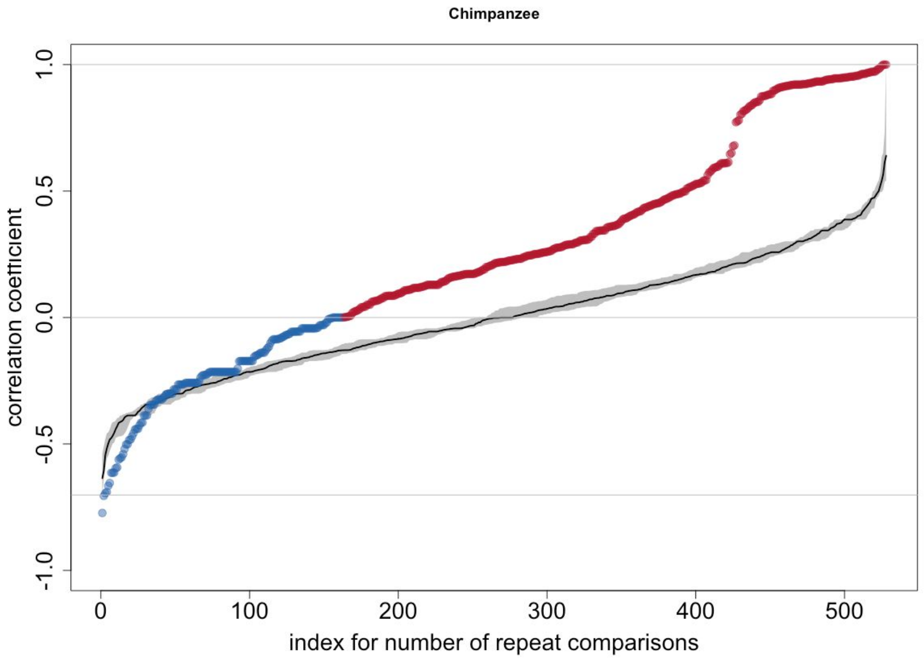
Chimpanzee
correlation coefficient
1.0
0.5
0.0
-0.5
-1.0
0
100
200
300
400
500
index for number of repeat comparisons

D

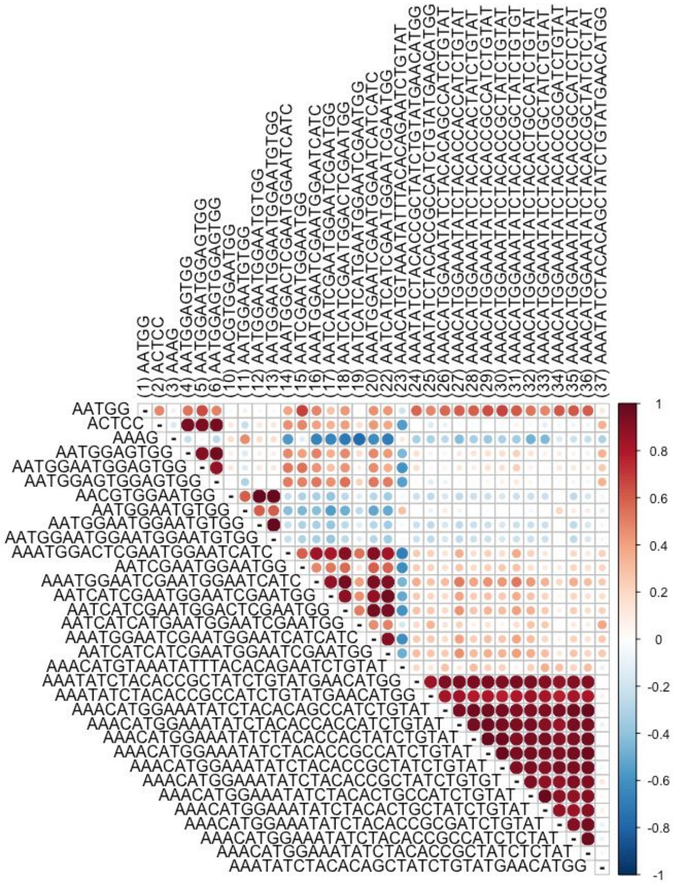
(1) AATGG
(2) ACTCC
(3) AAAG
(4) AATGGAGTGG
(5) AATGGAATGGAGTGG
(6) AATGGAGTGGAGTGG
(10) AACGTGGAATGG
(11) AATGGAATGTGG
(12) AATGGAATGGAATGTGG
(13) AATGGAATGGAATGGAATGTGG
(14) AAATGGACTCGAATGGAATCATC
(15) AATCGAATGGAATGG
(16) AAATGGAATCGAATGGAATCATC
(17) AATCATCGAATGGAATCGAATGG
(18) AATCATCGAATGGACTCGAATGG
(19) AATCATCATGAATGGAATCGAATGG
(20) AAATGGAATCGAATGGAATCATCATC
(22) AATCATCATCGAATGGAATCGAATGG
(23) AAACATGTAAATATTTACACAGAATCTGTAT
(24) AAATATCTACACCGCTATCTGTATGAACATGG
(25) AAATATCTACACCGCCATCTGTATGAACATGG
(26) AAACATGGAAATATCTACACAGCCATCTGTAT
(27) AAACATGGAAATATCTACACCACCATCTGTAT
(28) AAACATGGAAATATCTACACCACTATCTGTAT
(29) AAACATGGAAATATCTACACCGCCATCTGTAT
(30) AAACATGGAAATATCTACACCGCTATCTGTAT
(31) AAACATGGAAATATCTACACCGCTATCTGTGT
(32) AAACATGGAAATATCTACACTGCCATCTGTAT
(33) AAACATGGAAATATCTACACTGCTATCTGTAT
(34) AAACATGGAAATATCTACACCGCGATCTGTAT
(35) AAACATGGAAATATCTACACCGCCATCTCTAT
(36) AAACATGGAAATATCTACACCGCTATCTCTAT
(37) AAATATCTACACAGCTATCTGTATGAACATGG
AATGG
ACTCC
AAAG
AATGGAGTGG
AATGGAATGGAGTGG
AATGGAGTGGAGTGG
AACGTGGAATGG
AATGGAATGTGG
AATGGAATGGAATGTGG
AATGGAATGGAATGGAATGTGG
AAATGGACTCGAATGGAATCATC
AATCGAATGGAATGG
AAATGGAATCGAATGGAATCATC
AATCATCGAATGGAATCGAATGG
AATCATCGAATGGACTCGAATGG
AATCATCATGAATGGAATCGAATGG
AAATGGAATCGAATGGAATCATCATC
AATCATCATCGAATGGAATCGAATGG
AAACATGTAAATATTTACACAGAATCTGTAT
AAATATCTACACCGCTATCTGTATGAACATGG
AAATATCTACACCGCCATCTGTATGAACATGG
AAACATGGAAATATCTACACAGCCATCTGTAT
AAACATGGAAATATCTACACCACCATCTGTAT
AAACATGGAAATATCTACACCACTATCTGTAT
AAACATGGAAATATCTACACCGCCATCTGTAT
AAACATGGAAATATCTACACCGCTATCTGTAT
AAACATGGAAATATCTACACCGCTATCTGTGT
AAACATGGAAATATCTACACTGCCATCTGTAT
AAACATGGAAATATCTACACTGCTATCTGTAT
AAACATGGAAATATCTACACCGCGATCTGTAT
AAACATGGAAATATCTACACCGCCATCTCTAT
AAACATGGAAATATCTACACCGCTATCTCTAT
AAATATCTACACAGCTATCTGTATGAACATGG
1
0.8
0.6
0.4
0.2
0
-0.2
-0.4
-0.6
-0.8
-1

**E**

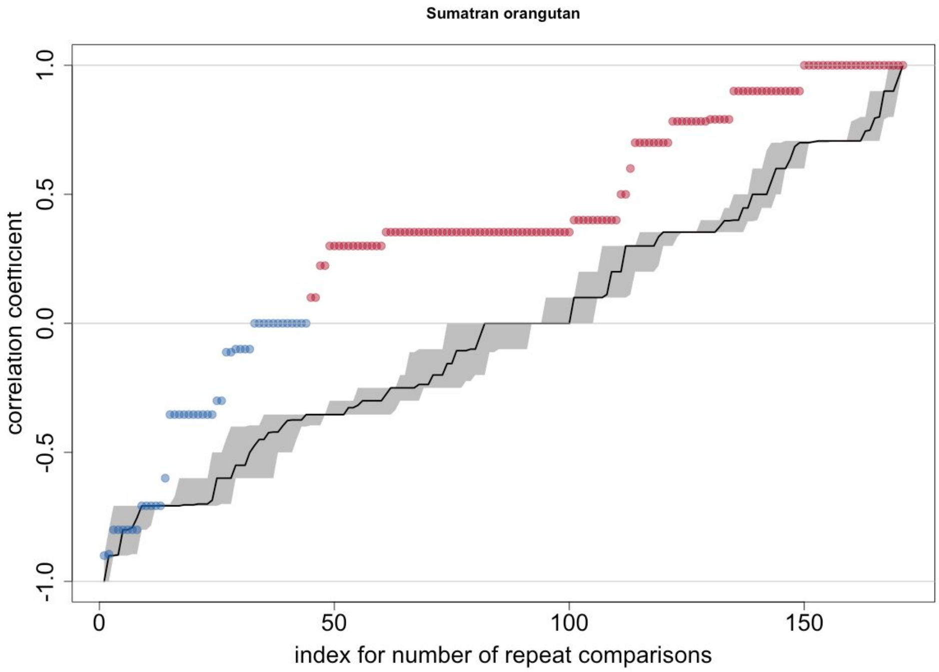

Sumatran orangutan
correlation coefficient
1.0
0.5
0.0
-0.5
-1.0
0
50
100
150
index for number of repeat comparisons


**F**

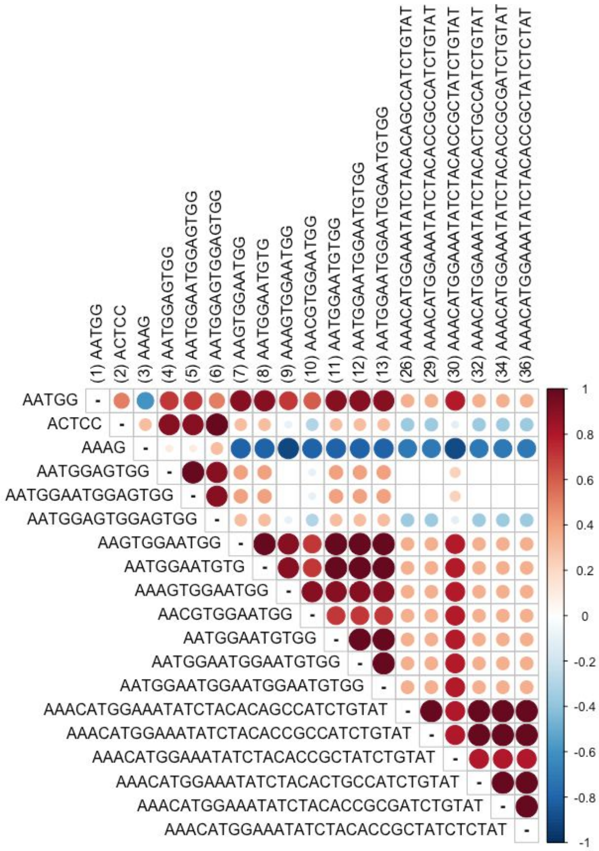

(1) AATGG
(2) ACTCC
(3) AAAG
(4) AATGGAGTGG
(5) AATGGAATGGAGTGG
(6) AATGGAGTGGAGTGG
(7) AAGTGGAATGG
(8) AATGGAATGTG
(9) AAAGTGGAATGG
(10) AACGTGGAATGG
(11) AATGGAATGTGG
(12) AATGGAATGGAATGTGG
(13) AATGGAATGGAATGGAATGTGG
(26) AAACATGGAAATATCTACACAGCCATCTGTAT
(29) AAACATGGAAATATCTACACCGCCATCTGTAT
(30) AAACATGGAAATATCTACACCGCTATCTGTAT
(32) AAACATGGAAATATCTACACTGCCATCTGTAT
(34) AAACATGGAAATATCTACACCGCGATCTGTAT
(36) AAACATGGAAATATCTACACCGCTATCTCTAT
AATGG
ACTCC
AAAG
AATGGAGTGG
AATGGAATGGAGTGG
AATGGAGTGGAGTGG
AAGTGGAATGG
AATGGAATGTG
AAAGTGGAATGG
AACGTGGAATGG
AATGGAATGTGG
AATGGAATGGAATGTGG
AATGGAATGGAATGGAATGTGG
AAACATGGAAATATCTACACAGCCATCTGTAT
AAACATGGAAATATCTACACCGCCATCTGTAT
AAACATGGAAATATCTACACCGCTATCTGTAT
AAACATGGAAATATCTACACTGCCATCTGTAT
AAACATGGAAATATCTACACCGCGATCTGTAT
AAACATGGAAATATCTACACCGCTATCTCTAT
1
0.8
0.6
0.4
0.2
0
-0.2
-0.4
-0.6
-0.8
-1

**G**

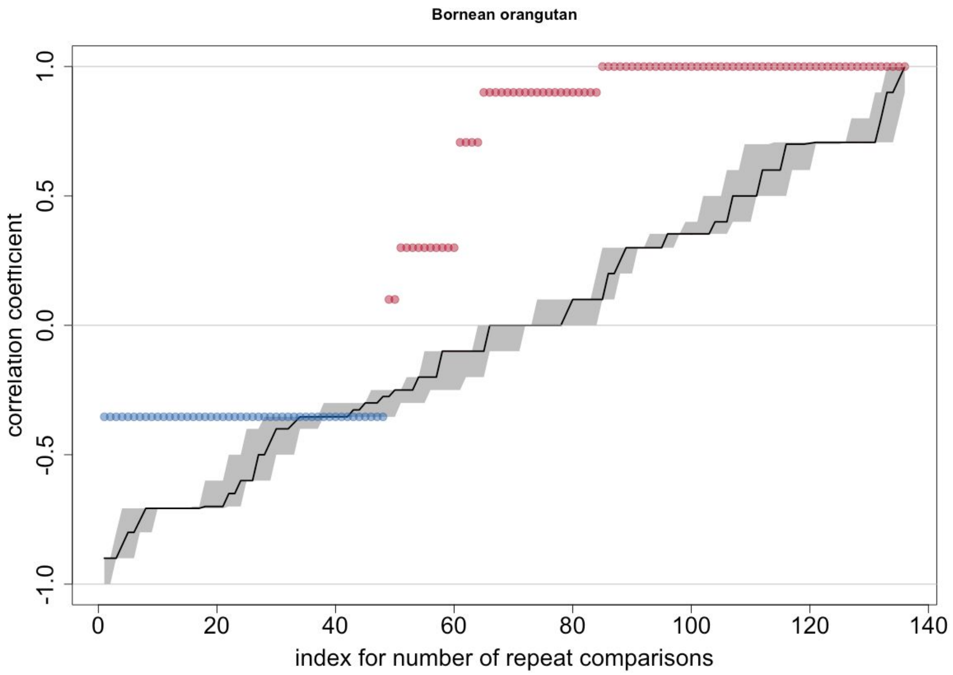
Bornean orangutan
correlation coefficient
index for number of repeat comparisons
1.0
0.5
0.0
-0.5
-1.0
0
20
40
60
80
100
120
140

**H**

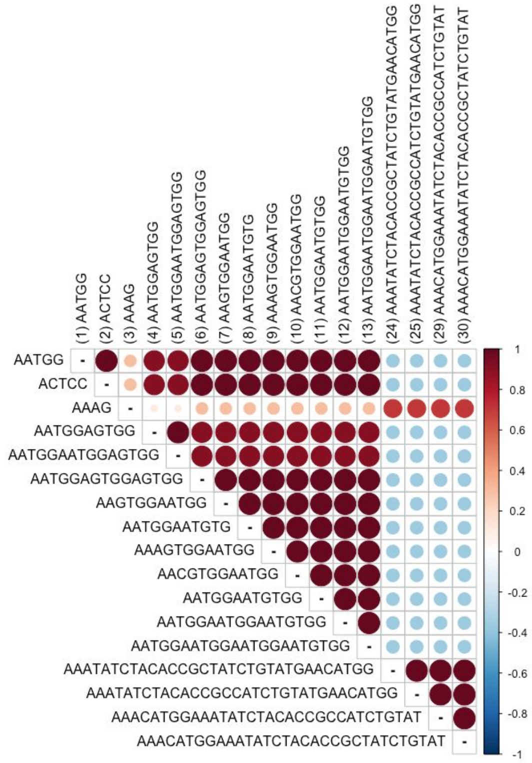
(1) AATGG
(2) ACTCC
(3) AAAG
(4) AATGGAGTGG
(5) AATGGAATGGAGTGG
(6) AATGGAGTGGAGTGG
(7) AAGTGGAATGG
(8) AATGGAATGTG
(9) AAAGTGGAATGG
(10) AACGTGGAATGG
(11) AATGGAATGTGG
(12) AATGGAATGGAATGTGG
(13) AATGGAATGGAATGGAATGTGG
(24) AAATATCTACACCGCTATCTGTATGAACATGG
(25) AAATATCTACACCGCCATCTGTATGAACATGG
(29) AAACATGGAAATATCTACACCGCCATCTGTAT
(30) AAACATGGAAATATCTACACCGCTATCTGTAT
AATGG
ACTCC
AAAG
AATGGAGTGG
AATGGAATGGAGTGG
AATGGAGTGGAGTGG
AAGTGGAATGG
AATGGAATGTG
AAAGTGGAATGG
AACGTGGAATGG
AATGGAATGTGG
AATGGAATGGAATGTGG
AATGGAATGGAATGGAATGTGG
AAATATCTACACCGCTATCTGTATGAACATGG
AAATATCTACACCGCCATCTGTATGAACATGG
AAACATGGAAATATCTACACCGCCATCTGTAT
AAACATGGAAATATCTACACCGCTATCTGTAT
1
0.8
0.6
0.4
0.2
0
-0.2
-0.4
-0.6
-0.8
-1

**Figure S13. Fluorescent in situ hybridization (FISH) analysis of: (A) DAPI-counterstained chimpanzee male chromosomes (lymphoblastoid cell line), the white arrow indicates the location of the Y chromosome, (B) both the whole bonobo Y chromosome painting probe (WBY) and the 5'-amine-modified 32-mer containing probe (Pan32) to DAPI-counterstained chimpanzee male chromosomes (the red arrow indicates that the chimpanzee Y chromosome is positive only for the WBY probe and not for the oligonucleotide probe), (C) only the 5'-amine-modified Pan32 probe to chimpanzee male chromosomes, (D) only the WBY probe to chimpanzee male chromosomes (the red arrow indicates the location of the Y chromosome).** The 5'-amine-modified Pan32 probe is labeled with Alexa Fluor (green). The WBY probe is labeled with digoxigenin (red). Scale bar = 10 um.

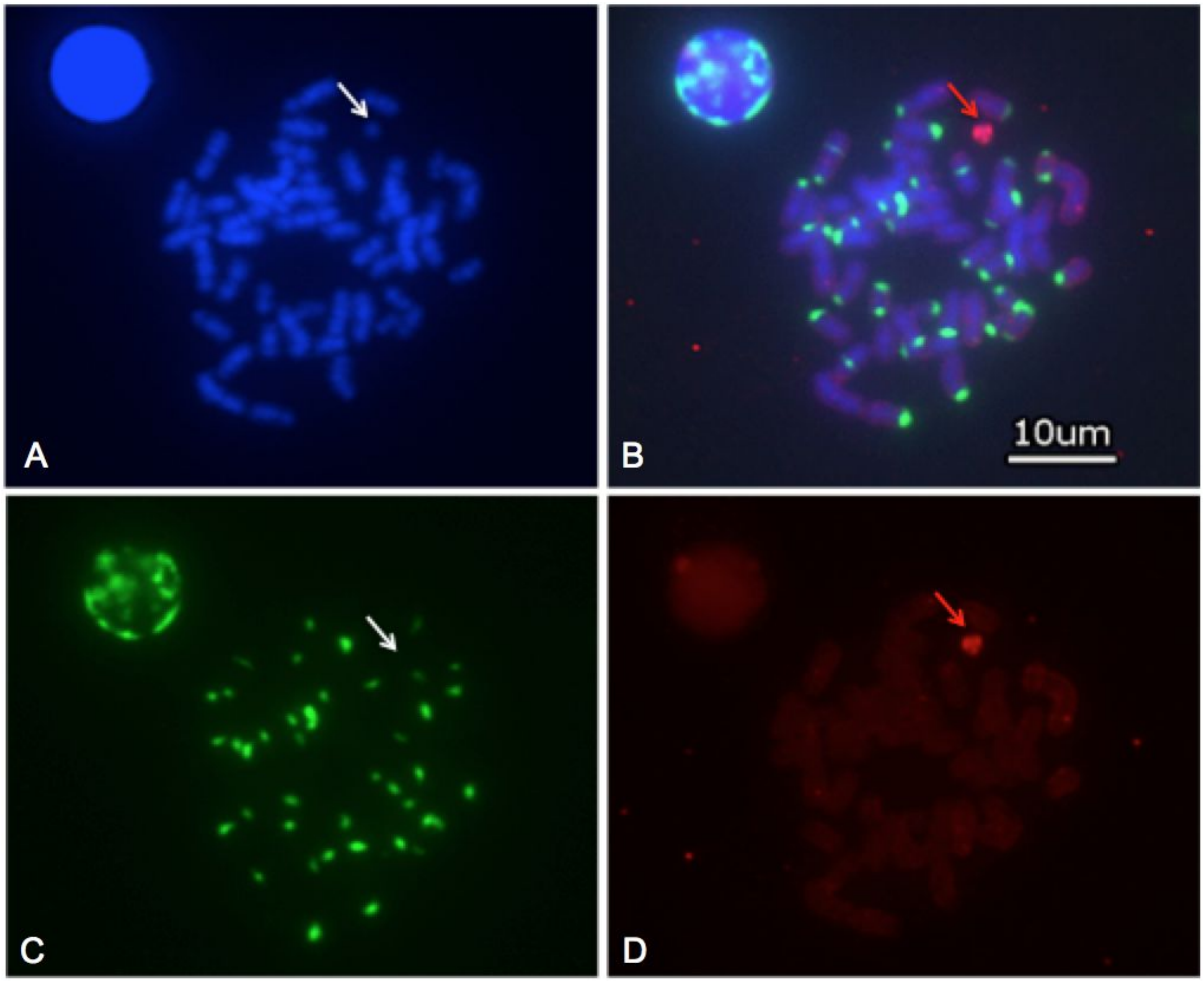

**Figure S14. Lengths of repeat arrays in Nanopore data.** Boxplots are plotted with outline=TRUE option. **(A)** Human. **(B)** Chimpanzee. **(C)** Bonobo. **(D)** Gorilla. **(E)** Sumatran orangutan. **(F)** Bornean orangutan.

## A

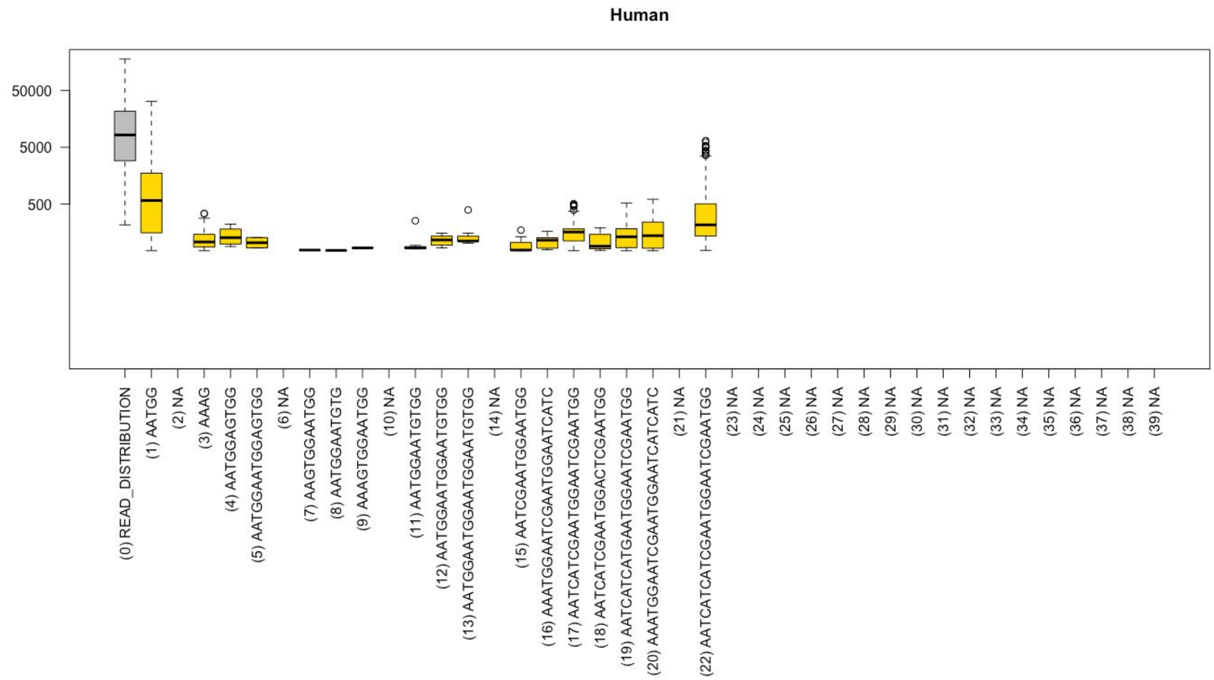

**B**

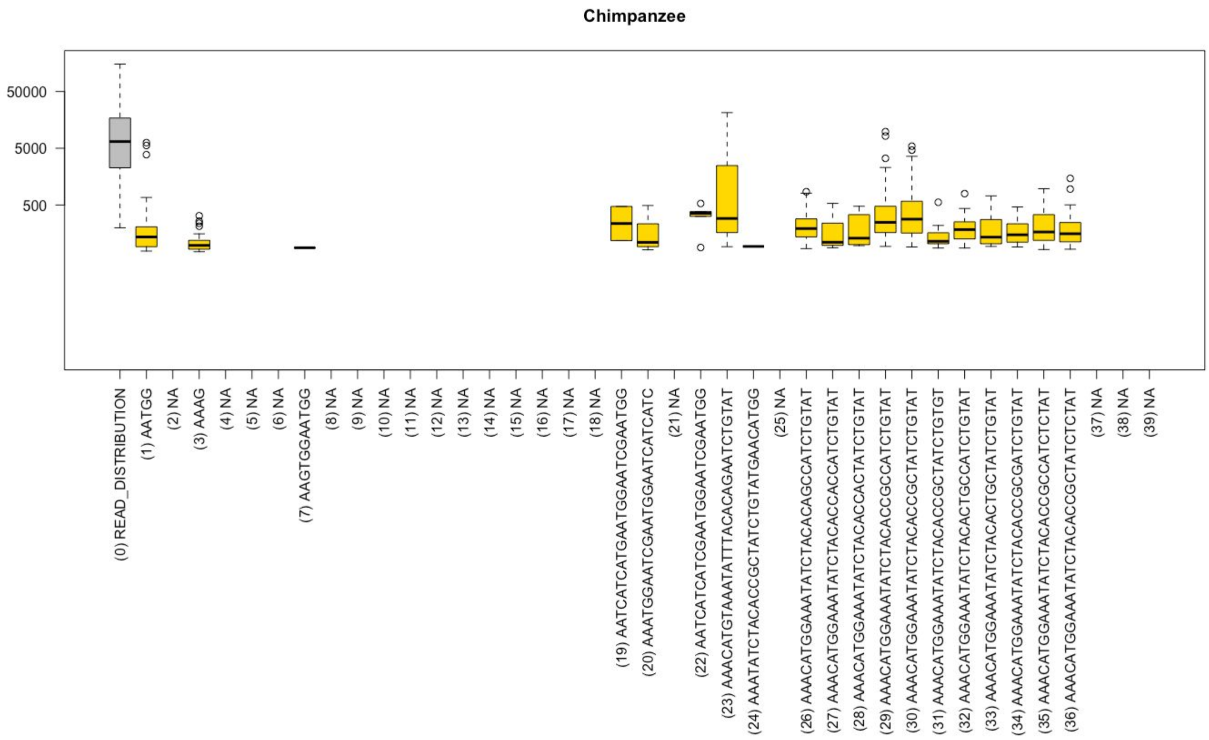
Chimpanzee
50000
5000
500
(0) READ_DISTRIBUTION
(1) AATGG
(2) NA
(3) AAAG
(4) NA
(5) NA
(6) NA
(7) AAGTGGAATGG
(8) NA
(9) NA
(10) NA
(11) NA
(12) NA
(13) NA
(14) NA
(15) NA
(16) NA
(17) NA
(18) NA
(19) AATCATCATGAATGGAATCGAATGG
(20) AAATGGAATCGAATGGAATCATCATC
(21) NA
(22) AATCATCATCGAATGGAATCGAATGG
(23) AAACATGTAAATATTTACACAGAATCTGTAT
(24) AAATATCTACACCGCTATCTGTATGAACATGG
(25) NA
(26) AAACATGGAAATATCTACACAGCCATCTGTAT
(27) AAACATGGAAATATCTACACCACCATCTGTAT
(28) AAACATGGAAATATCTACACCACTATCTGTAT
(29) AAACATGGAAATATCTACACCGCCATCTGTAT
(30) AAACATGGAAATATCTACACCGCTATCTGTAT
(31) AAACATGGAAATATCTACACCGCTATCTGTGT
(32) AAACATGGAAATATCTACACTGCCATCTGTAT
(33) AAACATGGAAATATCTACACTGCTATCTGTAT
(34) AAACATGGAAATATCTACACCGCGATCTGTAT
(35) AAACATGGAAATATCTACACCGCCATCTCTAT
(36) AAACATGGAAATATCTACACCGCTATCTCTAT
(37) NA
(38) NA
(39) NA

C

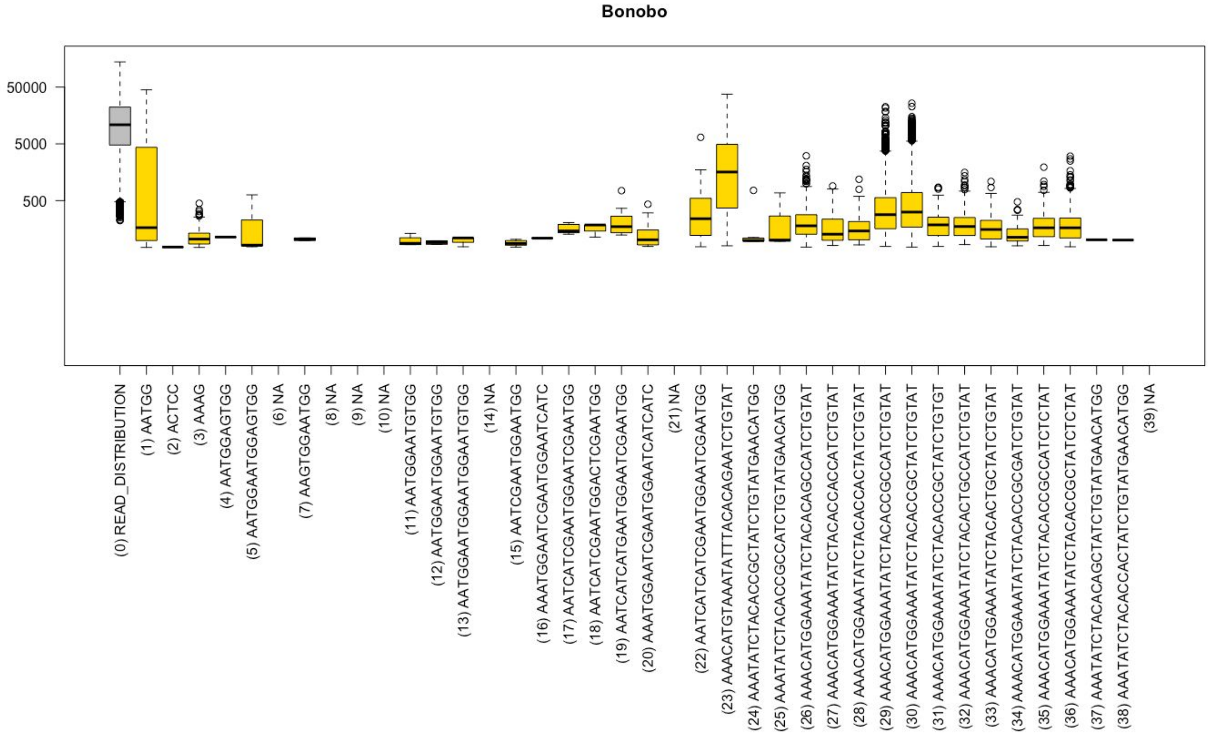

Bonobo
50000
5000
500
(0) READ_DISTRIBUTION
(1) AATGG
(2) ACTCC
(3) AAAG
(4) AATGGAGTGG
(5) AATGGAATGGAGTGG
(6) NA
(7) AAGTGGAATGG
(8) NA
(9) NA
(10) NA
(11) AATGGAATGTGG
(12) AATGGAATGGAATGTGG
(13) AATGGAATGGAATGGAATGTGG
(14) NA
(15) AATCGAATGGAATGG
(16) AAATGGAATCGAATGGAATCATC
(17) AATCATCGAATGGAATCGAATGG
(18) AATCATCGAATGGACTCGAATGG
(19) AATCATCATGAATGGAATCGAATGG
(20) AAATGGAATCGAATGGAATCATCATC
(21) NA
(22) AATCATCATCGAATGGAATCGAATGG
(23) AAACATGTAAATATTTACACAGAATCTGTAT
(24) AAATATCTACACCGCTATCTGTATGAACATGG
(25) AAATATCTACACCGCCATCTGTATGAACATGG
(26) AAACATGGAAATATCTACACAGCCATCTGTAT
(27) AAACATGGAAATATCTACACCACCATCTGTAT
(28) AAACATGGAAATATCTACACCACTATCTGTAT
(29) AAACATGGAAATATCTACACCGCCATCTGTAT
(30) AAACATGGAAATATCTACACCGCTATCTGTAT
(31) AAACATGGAAATATCTACACCGCTATCTGTGT
(32) AAACATGGAAATATCTACACTGCCATCTGTAT
(33) AAACATGGAAATATCTACACTGCTATCTGTAT
(34) AAACATGGAAATATCTACACCGCGATCTGTAT
(35) AAACATGGAAATATCTACACCGCCATCTCTAT
(36) AAACATGGAAATATCTACACCGCTATCTCTAT
(37) AAATATCTACACAGCTATCTGTATGAACATGG
(38) AAATATCTACACCACTATCTGTATGAACATGG
(39) NA


D

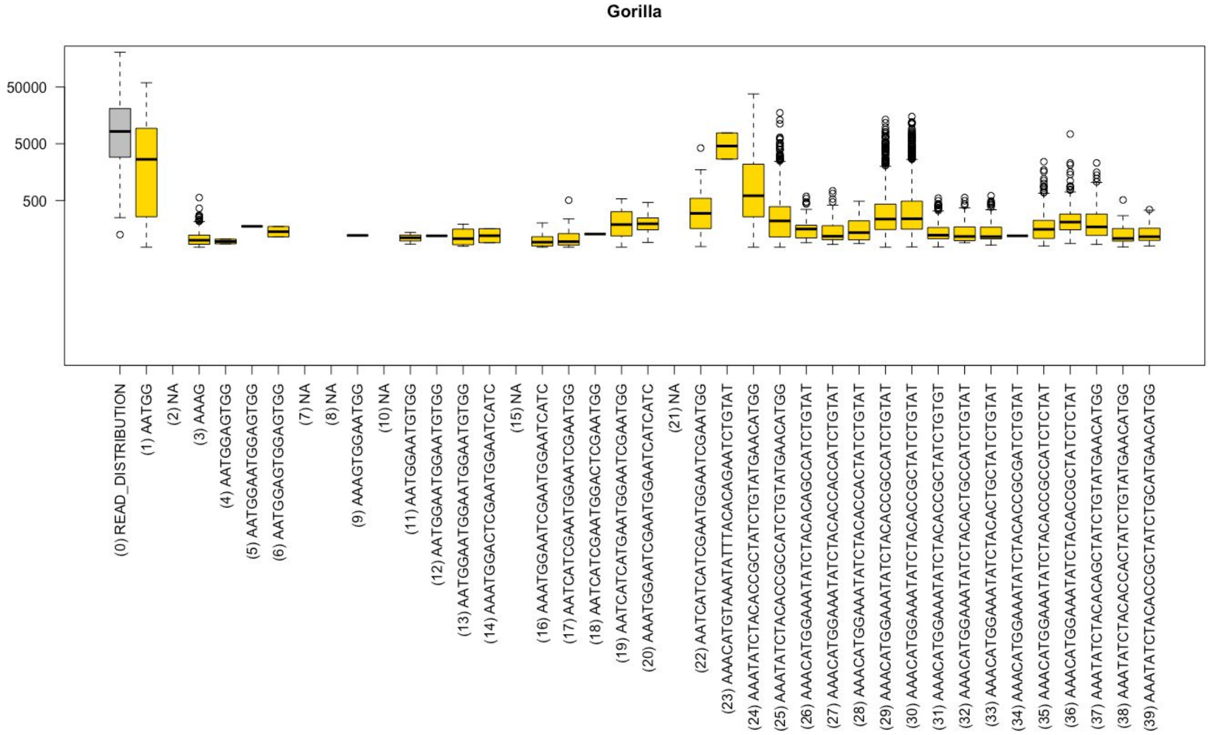

Gorilla
50000
5000
500
(0) READ_DISTRIBUTION
(1) AATGG
(2) NA
(3) AAAG
(4) AATGGAGTGG
(5) AATGGAATGGAGTGG
(6) AATGGAGTGGAGTGG
(7) NA
(8) NA
(9) AAAGTGGAATGG
(10) NA
(11) AATGGAATGTGG
(12) AATGGAATGGAATGTGG
(13) AATGGAATGGAATGGAATGTGG
(14) AAATGGACTCGAATGGAATCATC
(15) NA
(16) AAATGGAATCGAATGGAATCATC
(17) AATCATCGAATGGAATCGAATGG
(18) AATCATCGAATGGACTCGAATGG
(19) AATCATCATGAATGGAATCGAATGG
(20) AAATGGAATCGAATGGAATCATCATC
(21) NA
(22) AATCATCATCGAATGGAATCGAATGG
(23) AAACATGTAAATATTTACACAGAATCTGTAT
(24) AAATATCTACACCGCTATCTGTATGAACATGG
(25) AAATATCTACACCGCCATCTGTATGAACATGG
(26) AAACATGGAAATATCTACACAGCCATCTGTAT
(27) AAACATGGAAATATCTACACCACCATCTGTAT
(28) AAACATGGAAATATCTACACCACTATCTGTAT
(29) AAACATGGAAATATCTACACCGCCATCTGTAT
(30) AAACATGGAAATATCTACACCGCTATCTGTAT
(31) AAACATGGAAATATCTACACCGCTATCTGTGT
(32) AAACATGGAAATATCTACACTGCCATCTGTAT
(33) AAACATGGAAATATCTACACTGCTATCTGTAT
(34) AAACATGGAAATATCTACACCGCGATCTGTAT
(35) AAACATGGAAATATCTACACCGCCATCTCTAT
(36) AAACATGGAAATATCTACACCGCTATCTCTAT
(37) AAATATCTACACAGCTATCTGTATGAACATGG
(38) AAATATCTACACCACTATCTGTATGAACATGG
(39) AAATATCTACACCGCTATCTGCATGAACATGG

**E**

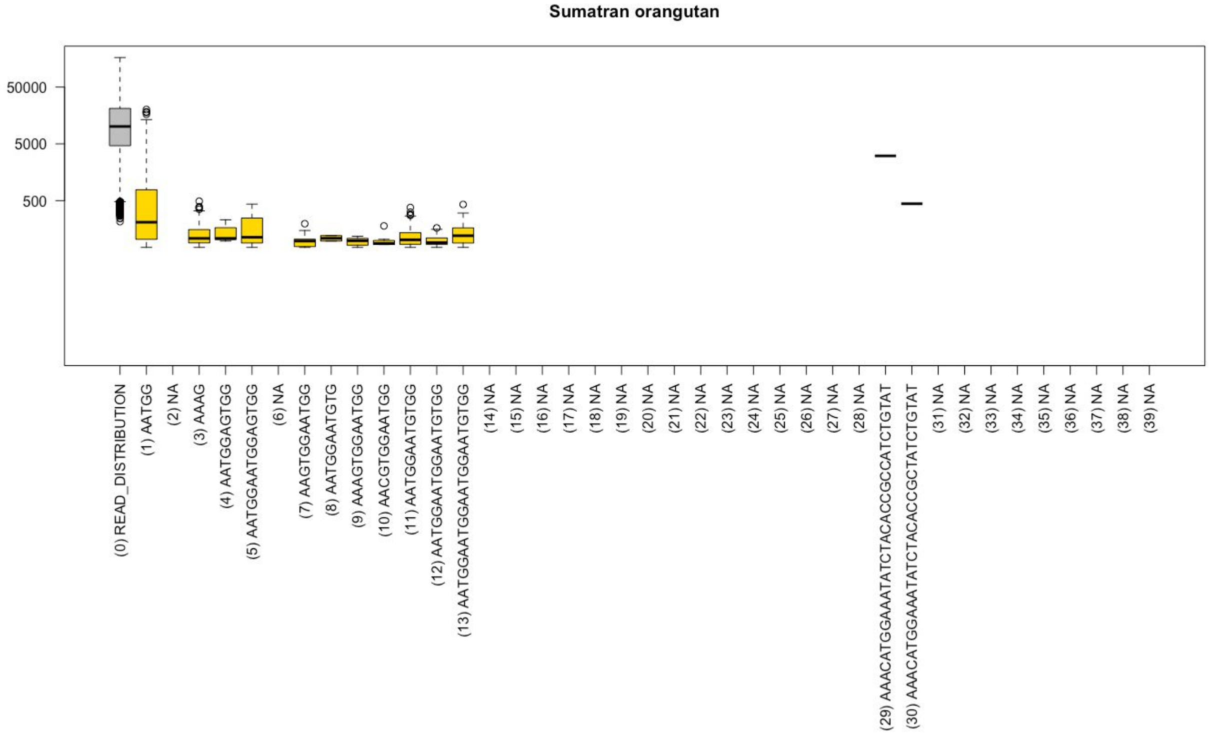

Sumatran orangutan
50000
5000
500
(0) READ_DISTRIBUTION
(1) AATGG
(2) NA
(3) AAAG
(4) AATGGAGTGG
(5) AATGGAATGGAGTGG
(6) NA
(7) AAGTGGAATGG
(8) AATGGAATGTG
(9) AAAGTGGAATGG
(10) AACGTGGAATGG
(11) AATGGAATGTGG
(12) AATGGAATGGAATGTGG
(13) AATGGAATGGAATGGAATGTGG
(14) NA
(15) NA
(16) NA
(17) NA
(18) NA
(19) NA
(20) NA
(21) NA
(22) NA
(23) NA
(24) NA
(25) NA
(26) NA
(27) NA
(28) NA
(29) AAACATGGAAATATCTACACCGCCATCTCTGTAT
(30) AAACATGGAAATATCTACACCGCTATCTCTGTAT
(31) NA
(32) NA
(33) NA
(34) NA
(35) NA
(36) NA
(37) NA
(38) NA
(39) NA


**F**

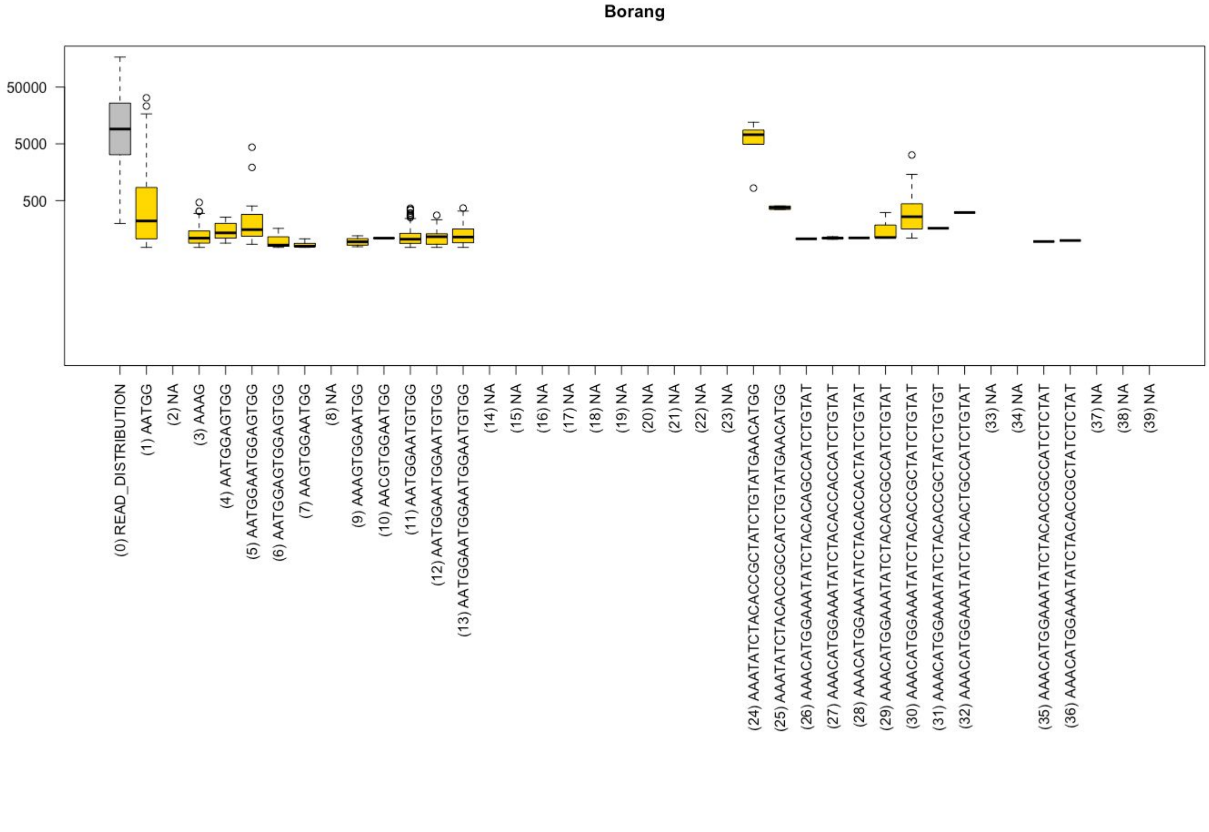

Borang
50000
5000
500
(0) READ_DISTRIBUTION
(1) AATGG
(2) NA
(3) AAAG
(4) AATGGAGTGG
(5) AATGGAATGGAGTGG
(6) AATGGAGTGGAGTGG
(7) AAGTGGAATGG
(8) NA
(9) AAAGTGGAATGG
(10) AACGTGGAATGG
(11) AATGGAATGTGG
(12) AATGGAATGGAATGTGG
(13) AATGGAATGGAATGGAATGTGG
(14) NA
(15) NA
(16) NA
(17) NA
(18) NA
(19) NA
(20) NA
(21) NA
(22) NA
(23) NA
(24) AAATATCTACACCGCTATCTCTGTATGAACATGG
(25) AAATATCTACACCGCCATCTCTGTATGAACATGG
(26) AAACATGGAAATATCTACACAGCCATCTCTGTAT
(27) AAACATGGAAATATCTACACCACCATCTCTGTAT
(28) AAACATGGAAATATCTACACCACTATCTCTGTAT
(29) AAACATGGAAATATCTACACCGCCATCTCTGTAT
(30) AAACATGGAAATATCTACACCGCTATCTCTGTAT
(31) AAACATGGAAATATCTACACCGCTATCTCTGTGT
(32) AAACATGGAAATATCTACACTGCCATCTCTGTAT
(33) NA
(34) NA
(35) AAACATGGAAATATCTACACCGCCATCTCTCTAT
(36) AAACATGGAAATATCTACACCGCTATCTCTCTAT
(37) NA
(38) NA
(39) NA

**Figure S15. Lengths of repeat arrays in PacBio data.** Boxplots are plotted with outline=TRUE option. **(A)** Human. **(B)** Chimpanzee. **(C)** Gorilla. **(D)** Sumatran orangutan.

**A**

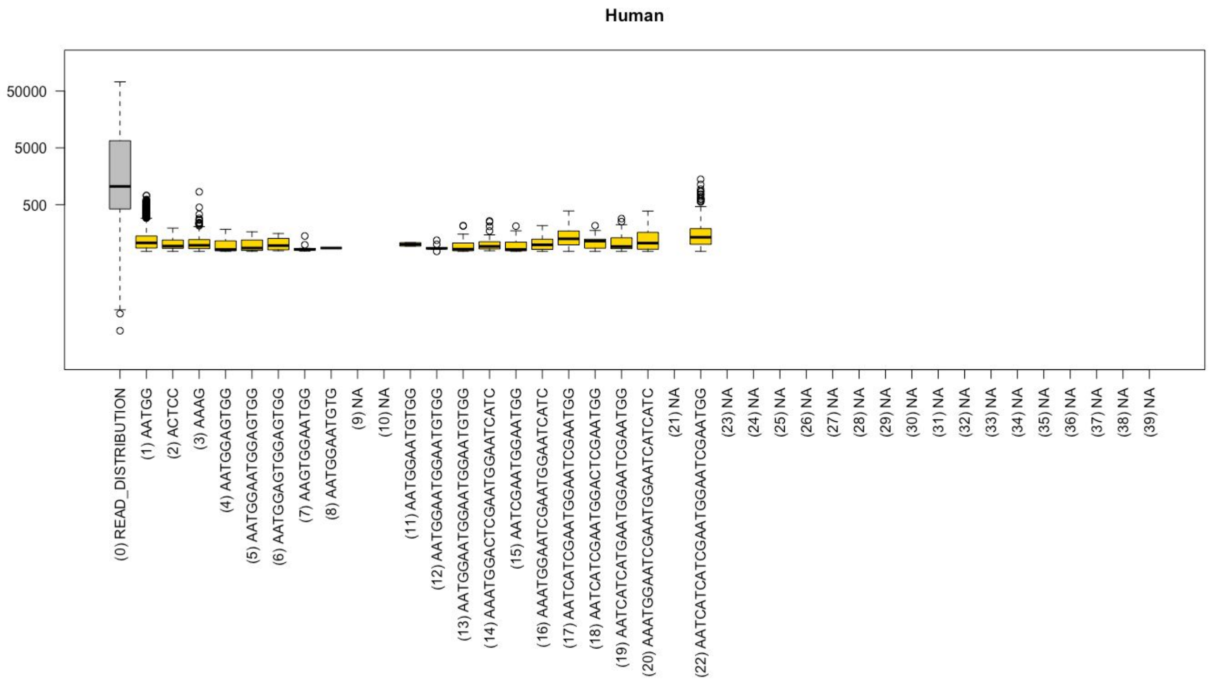


**B**

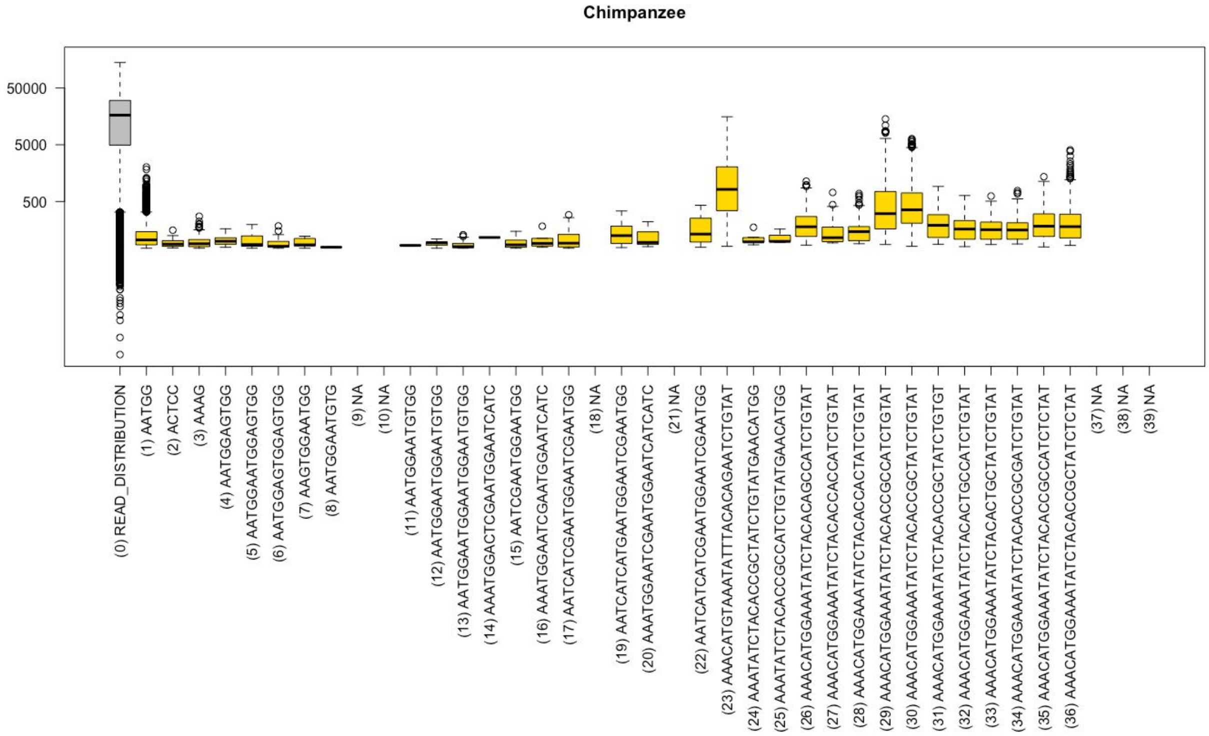

C

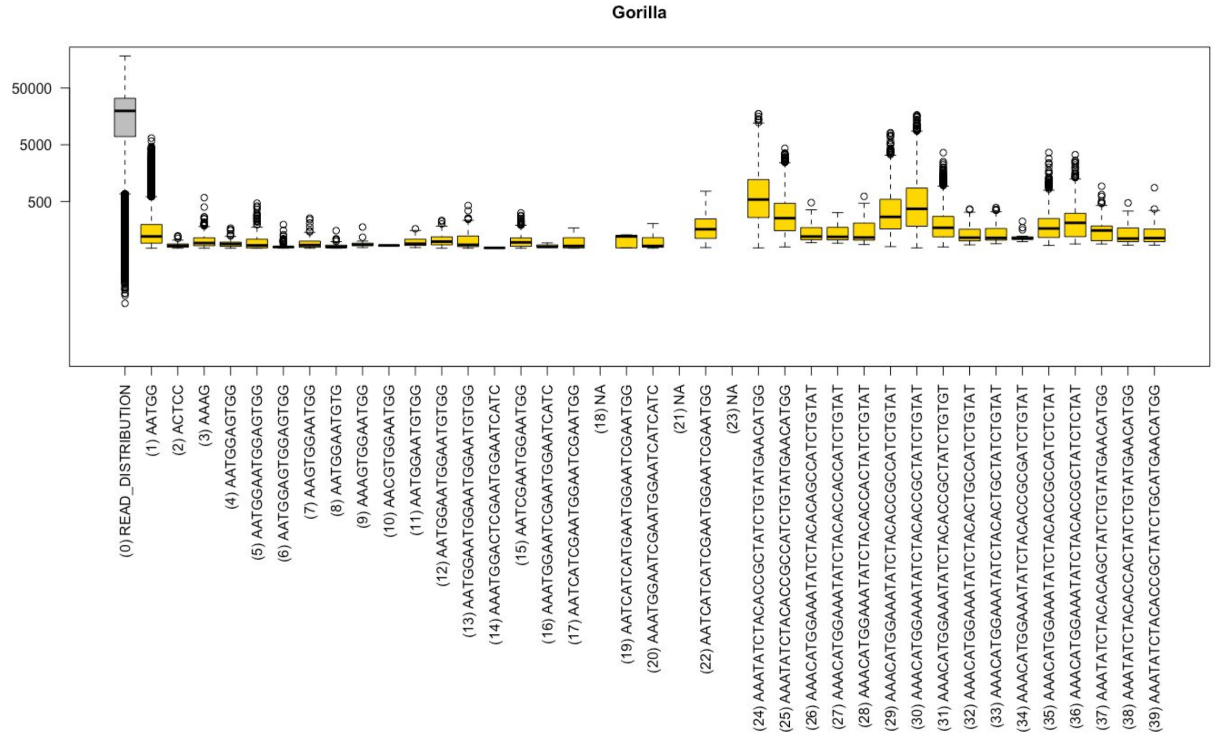

Gorilla
50000
5000
500
(0) READ_DISTRIBUTION
(1) AATGG
(2) ACTCC
(3) AAAG
(4) AATGGAGTGG
(5) AATGGAATGGAGTGG
(6) AATGGAGTGGAGTGG
(7) AAGTGGAATGG
(8) AATGGAATGTG
(9) AAAGTGGAATGG
(10) AACGTGGAATGG
(11) AATGGAATGTGG
(12) AATGGAATGGAATGTGG
(13) AATGGAATGGAATGGAATGTGG
(14) AAATGGACTCGAATGGAATCATC
(15) AATCGAATGGAATGG
(16) AAATGGAATCGAATGGAATCATC
(17) AATCATCGAATGGAATCGAATGG
(18) NA
(19) AATCATCATGAATGGAATCGAATGG
(20) AAATGGAATCGAATGGAATCATCATC
(21) NA
(22) AATCATCATCGAATGGAATCGAATGG
(23) NA
(24) AAATATCTACACCGCTATCTGTATGAACATGG
(25) AAATATCTACACCGCCATCTGTATGAACATGG
(26) AAACATGGAAATATCTACACAGCCATCTGTAT
(27) AAACATGGAAATATCTACACCACCATCTGTAT
(28) AAACATGGAAATATCTACACCACTATCTGTAT
(29) AAACATGGAAATATCTACACCGCCATCTGTAT
(30) AAACATGGAAATATCTACACCGCTATCTGTAT
(31) AAACATGGAAATATCTACACCGCTATCTGTGT
(32) AAACATGGAAATATCTACACTGCCATCTGTAT
(33) AAACATGGAAATATCTACACTGCTATCTGTAT
(34) AAACATGGAAATATCTACACCGCGATCTGTAT
(35) AAACATGGAAATATCTACACCGCCATCTCTAT
(36) AAACATGGAAATATCTACACCGCTATCTCTAT
(37) AAATATCTACACAGCTATCTGTATGAACATGG
(38) AAATATCTACACCACTATCTGTATGAACATGG
(39) AAATATCTACACCGCTATCTGCATGAACATGG


D

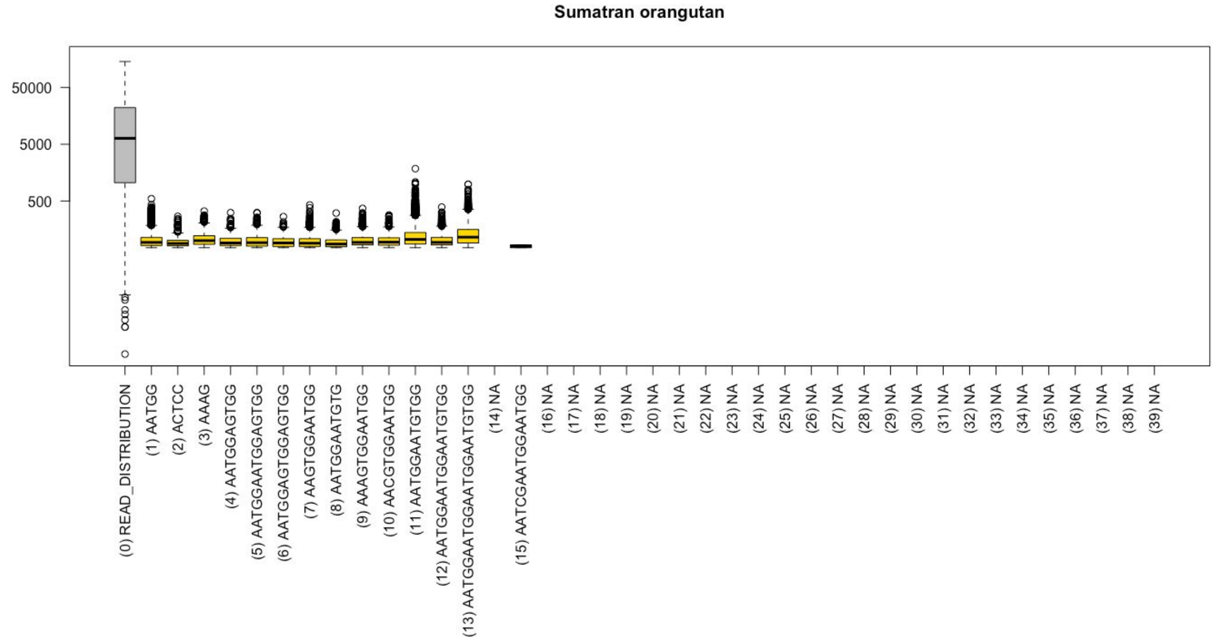

Sumatran orangutan
50000
5000
500
(0) READ_DISTRIBUTION
(1) AATGG
(2) ACTCC
(3) AAAG
(4) AATGGAGTGG
(5) AATGGAATGGAGTGG
(6) AATGGAGTGGAGTGG
(7) AAGTGGAATGG
(8) AATGGAATGTG
(9) AAAGTGGAATGG
(10) AACGTGGAATGG
(11) AATGGAATGTGG
(12) AATGGAATGGAATGTGG
(13) AATGGAATGGAATGGAATGTGG
(14) NA
(15) AATCGAATGGAATGG
(16) NA
(17) NA
(18) NA
(19) NA
(20) NA
(21) NA
(22) NA
(23) NA
(24) NA
(25) NA
(26) NA
(27) NA
(28) NA
(29) NA
(30) NA
(31) NA
(32) NA
(33) NA
(34) NA
(35) NA
(36) NA
(37) NA
(38) NA
(39) NA

**Figure S16. The analysis of telomeric satellite TTAGGG (AACCCT).** (A) Low repeat density (kb/Mb) of telomeric repeat in the Illumina data, separated by species. (B) Low repeat density (kb/Mb) of telomeric repeat in the Nanopore and PacBio data, respectively. (C) and (D) Telomeric satellite identified in the Nanopore and PacBio data, respectively. Each rectangle represents a single repeat array. The width of the rectangle represents a length of the repeat array, and the location on the x-axis the distance to the end of read. The size of the sequenced dataset (Mb) is indicated adjacent to the plot; note lower sequencing depth in Nanopore as opposed to PacBio data.

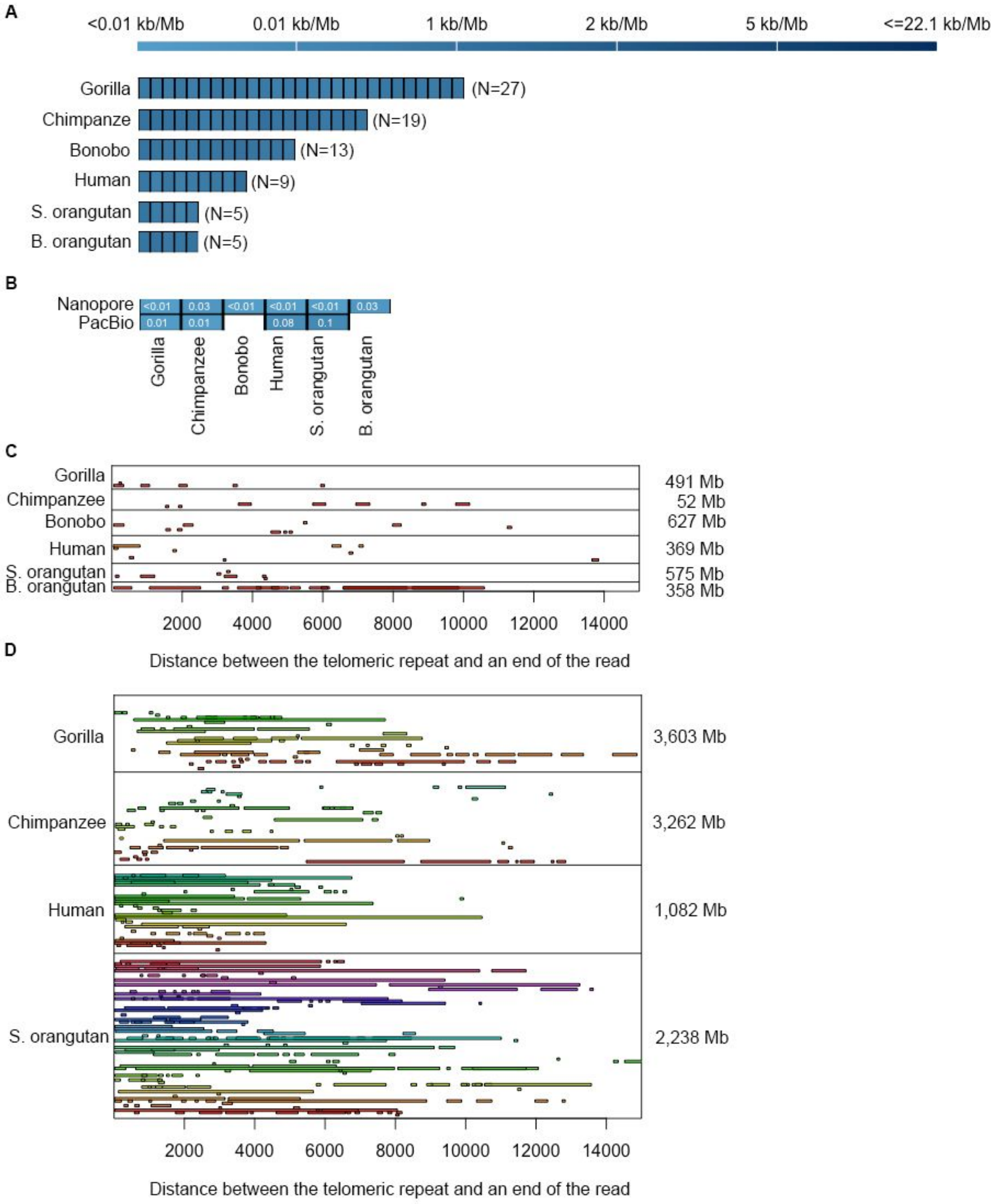
