## Supplementary Tables for "High satellite repeat turnover in great apes studied with short- and long-read technologies"

**Table S1. Data used in this study.**

The individual number of reads for each data file (after filtering) are available in the repository file “rcounts”.

| Species | Sample size | M:F* | Subspecies | Sample size | M:F* | number of libraries (technical replicates) | insert size |
|---|---|---|---|---|---|---|---|
| | | | Diverse human populations from HGDP | 9 | 9:0 | 9 | |
| Human (Homo sapiens) | 18 | 14:4 | Families of human trios | 9 | 5:4 | 9 | |
| | | | Nigeria-Cameroon chimpanzee (Pan troglodytes ellioti) | 4 | 1:3 | 4 | 200-233 |
| | | | Eastern chimpanzee (Pan troglodytes schweinfurthii) | 6 | 2:4 | 35 | 212-507 |
| | | | Central chimpanzee(Pan troglodytes troglodytes) | 4 | 1:3 | 19 | 434-501 |
| | | | Western chimpanzee(Pan troglodytes verus) | 4 | 3:1 | 21 | 211-492 |
| Chimpanzee(Pan troglodytes) | 19 | 10:9 | Hybrid of Western and Central chimpanzee(Pan troglo | 1 | 1:0 | 4 | 214-387 |
| Bonobo (Pan paniscus) | 13 | 2:11 | Bonobo (Pan paniscus) | | | 69 | 532 |
| Gorilla (Gorilla) | 27 | 6:21 | Eastern lowland gorilla (Gorilla beringei graueri) | 3 | 2:1 | 18 | 472 |
| | | | Cross river gorilla(Gorilla gorilla diehli) | 1 | 0:1 | 4 | 450 |
| | | | Western lowland gorilla(Gorilla gorilla gorilla) | 23 | 4:19 | 82 | 522 |
| Sumatran orangutan (Pongo abelii) | 5 | 1:4 | Sumatran orangutan(Pongo abelii) | 5 | 1:4 | 24 | 460-506 |
| Bornean orangutan(Pongo pygmaeus) | 5 | 1:4 | Bornean orangutan(Pongo pygmaeus) | 5 | 1:4 | 15 | 463-503 |

* M:F represents the ratio of males to females.

| Human trios | Source | amplification | Read length | sequencer |
|---|---|---|---|---|
| Son77 family | Illumina pla | PCR- | 101 | HiSeq2000 |
| Daughter78 family | Illumina pla | PCR- | 101 | HiSeq2000 |
| Ashkenazi family | GIAB | PCR- | 150 trimmed to 100 | HiSeq2500 |

**Table S2. 39 abundant repeated motifs.**

List of repeated motifs that are potential derivatives of (AATGG)n repeat:

| index | motif |
|---|---|
| 1 | AATGG |
| 2 | ACTCC |
| 4 | AATGGAGTGG |
| 7 | AAGTGGAATGG |
| 8 | AATGGAATGTG |
| 9 | AAAGTGGAATGG |
| 10 | AACGTGGAATGG |
| 11 | AATGGAATGTGG |
| 5 | AATGGAATGGAGTGG |
| 6 | AATGGAGTGGAGTGG |
| 15 | AATCGAATGGAATGG |
| 12 | AATGGAATGGAATGTGG |
| 13 | AATGGAATGGAATGGAATGTGG |
| 14 | AAATGGACTCGAATGGAATCATC |
| 16 | AAATGGAATCGAATGGAATCATC |
| 17 | AATCATCGAATGGAATCGAATGG |
| 18 | AATCATCGAATGGACTCGAATGG |
| 19 | AATCATCATGAATGGAATCGAATGG |
| 20 | AAATGGAATCGAATGGAATCATCATC |
| 21 | AAATGGAATCGAATGTAATCATCATC |
| 22 | AATCATCATCGAATGGAATCGAATGG |

List of StSats: 32-mers and single 31-mer:

| index | motif |
|---|---|
| 23 | AAACATGTAAATATTTACACAGAATCTGTAT |
| 26 | AAACATGGAAATATCTACACAGCCATCTGTAT |
| 27 | AAACATGGAAATATCTACACCACCATCTGTAT |
| 28 | AAACATGGAAATATCTACACCACTATCTGTAT |
| 29 | AAACATGGAAATATCTACACCGCCATCTGTAT |
| 30 | AAACATGGAAATATCTACACCGCTATCTGTAT |
| 31 | AAACATGGAAATATCTACACCGCTATCTGTGT |
| 32 | AAACATGGAAATATCTACACTGCCATCTGTAT |
| 33 | AAACATGGAAATATCTACACTGCTATCTGTAT |
| 34 | AAACATGGAAATATCTACACCGCGATCTGTAT |
| 35 | AAACATGGAAATATCTACACCGCCATCTCTAT |
| 36 | AAACATGGAAATATCTACACCGCTATCTCTAT |
| 37 | AAATATCTACACAGCTATCTGTATGAACATGG |
| 38 | AAATATCTACACCACTATCTGTATGAACATGG |
| 24 | AAATATCTACACCGCTATCTGTATGAACATGG |
| 25 | AAATATCTACACCGCCATCTGTATGAACATGG |
| 39 | AAATATCTACACCGCTATCTGCATGAACATGG |

**Table S3.** The analysis of the telomeric repeat (TTAGGG)n

| ILLUMINA | Repeat density (kb/Mb) | % of the total satellite repeat density |
|---|---|---|
| **Human** | 0.0278 | 0.234% |
| **Chimpanzee** | 0.0422 | 0.100% |
| **Bonobo** | 0.0332 | 0.063% |
| **Gorilla** | 0.0233 | 0.024% |
| **S. orangutan** | 0.0227 | 0.100% |
| **B. orangutan** | 0.0361 | 0.110% |

| NANOPORE | Repeat density (kb/Mb) | % of the total satellite repeat density |
|---|---|---|
| **Human** | 0.0049 | 0.041% |
| **Chimpanzee** | 0.0330 | 0.078% |
| **Bonobo** | 0.0026 | 0.005% |
| **Gorilla** | 0.0020 | 0.002% |
| **S. orangutan** | 0.0019 | 0.009% |
| **B. orangutan** | 0.0320 | 0.098% |

| PACBIO | Repeat density (kb/Mb) | % of the total satellite repeat density |
|---|---|---|
| **Human** | 0.0764 | 0.642% |
| **Chimpanzee** | 0.0110 | 0.026% |
| **Bonobo** | NA | NA |
| **Gorilla** | 0.0140 | 0.014% |
| **S. orangutan** | 0.0974 | 0.430% |
| **B. orangutan** | NA | NA |

**Table S4. Intra-species variability of GGAAT repeated motif.**

| species | GGAAT variability mean fold difference of | sample size |
|---|---|---|
| **Human** | 1.23 | 9* |
| **Chimpanzee** | 1.47 | 19 |
| **Bonobo** | 1.32 | 13 |
| **Gorilla** | 1.51 | 27 |
| **Sumatran orangutan** | 1.25 | 5 |
| **Bornean orangutan** | 1.43 | 5 |

| MALES | GGAAT variability mean fold [illegible] | sample size |
|---|---|---|
| **Human** | 1.33 | 14** |
| **Chimpanzee** | 1.55 | 10 |
| **Bonobo** | 1.18 | 2 |
| **Gorilla** | 1.60 | 6 |
| **Sumatran orangutan** | NA | 1 |
| **Bornean orangutan** | NA | 1 |

| FEMALES | GGAAT variability mean fold [illegible] | sample size |
|---|---|---|
| **Human** | 1.26 | 4** |
| **Chimpanzee** | 1.40 | 9 |
| **Bonobo** | 1.28 | 11 |
| **Gorilla** | 1.48 | 21 |
| **Sumatran orangutan** | 1.31 | 4 |
| **Bornean orangutan** | 1.49 | 4 |

*Human male individuals from HGDP panel

**Human male and female individuals from HGDP and trio families

**Table S5. Classification algorithms**

**PCA**

| | PC1 | PC2 | PC3 | PC4 | PC5 | PC6 |
|---|---|---|---|---|---|---|
| Standard deviation | 1214666.457 | 438132.9149 | 316137.1414 | 175591.5672 | 22994.35773 | 20259.71864 |
| Proportion of Variance | 0.8197 | 0.1066 | 0.05553 | 0.01713 | 0.00029 | 0.00023 |
| Cumulative Proportion | 0.8197 | 0.9264 | 0.98193 | 0.99906 | 0.99935 | 0.99958 |

We took the subset of 39 most abundant repeats and ran lda (R package MASS version 7.3-50) and Random Forest (R package randomForest 4.6-12) in order to classify individuals into species.

**Random Forest**

Bornean and Sumatran Orangutans were misclassified as each other and one chimpanzee was classified as bonobo. We used 10,000 trees with seed=1 and equal priors.

Confusion matrix:

Type of random forest: classification

Number of trees: 10000

No. of variables tried at each split: 6

OOB estimate of error rate: 7.69%

Confusion matrix:

| | Bonobo | Bornean | Chimpanzee | Gorilla | Homo | Sumatran | class.error |
|---|---|---|---|---|---|---|---|
| Bonobo | 13 | 0 | 0 | 0 | 0 | 0 | 0 |
| Bornean | 0 | 3 | 0 | 0 | 0 | 2 | 0.4 |
| Chimpanzee | 1 | 0 | 18 | 0 | 0 | 0 | 0.05263158 |
| Gorilla | 0 | 0 | 0 | 27 | 0 | 0 | 0 |
| Homo | 0 | 0 | 0 | 0 | 9 | 0 | 0 |
| Sumatran | 0 | 3 | 0 | 0 | 0 | 2 | 0.6 |

**Table S6. Analysis of male-baised repeated motifs.**

For each repeated motif, we tested for the differences in repeat density between male and females using Mann-Whitney test (see p-value column; significant values are listed in red) and listed the ratio of male-to-female repeat density. Cells with gray background represent repeated motifs not found to be male-biased in a given species. The repeated motifs used for the probe design is highlighted in yellow. The Sumatran and Bornean orangutans were analyzed jointly due to the lower sample size.

| | | **HUMAN** | | **CHIMPANZEE** | | **BONOBO** | | **GORILLA** | | **ORANGUTAN** | |
|---|---|---|---|---|---|---|---|---|---|---|---|
| index | motif | p-value | ratio | p-value | ratio | p-value | ratio | p-value | ratio | p-value | ratio |
| 1 | AATGG | 0.02 | 1.52 | 0.16 | 1.18 | 0.05 | 1.36 | 0.2 | 1.24 | | |
| 2 | ACTCC | | | | | | | | | 0.09 | 1.47 |
| 24 | AAATATCTACACCGCTATCTGTATGAACATGG | | | | | | | 0.11 | 1.25 | | |
| 25 | AAATATCTACACCGCCATCTGTATGAACATGG | | | | | | | 0.13 | 1.2 | | |
| 26 | AAACATGGAAATATCTACACAGCCATCTGTAT | | | | | 0.03 | 1.26 | | | | |
| 29 | AAACATGGAAATATCTACACCGCCATCTGTAT | | | 0.2 | 1.21 | 0.05 | 1.26 | 0.19 | 1.12 | | |
| 30 | AAACATGGAAATATCTACACCGCTATCTGTAT | | | 0.2 | 1.18 | 0.03 | 1.23 | 0.16 | 1.17 | | |
| 31 | AAACATGGAAATATCTACACCGCTATCTGTGT | | | | | | | 0.17 | 1.24 | | |
| 32 | AAACATGGAAATATCTACACTGCCATCTGTAT | | | 0.11 | 1.24 | 0.05 | 1.21 | | | | |
| 33 | AAACATGGAAATATCTACACTGCTATCTGTAT | | | 0.14 | 1.21 | 0.05 | 1.24 | 0.11 | 1.19 | | |
| 34 | AAACATGGAAATATCTACACCGCGATCTGTAT | | | | | 0.12 | 1.3 | | | | |
| 35 | AAACATGGAAATATCTACACCGCCATCTCTAT | | | | | 0.08 | 1.22 | 0.14 | 1.17 | | |
| 36 | AAACATGGAAATATCTACACCGCTATCTCTAT | | | | | 0.03 | 1.22 | 0.19 | 1.13 | | |
| 37 | AAATATCTACACAGCTATCTGTATGAACATGG | | | | | | | 0.2 | 1.17 | | |
| 38 | AAATATCTACACCACTATCTGTATGAACATGG | | | | | | | 0.2 | 1.17 | | |
| 39 | AAATATCTACACCGCTATCTGCATGAACATGG | | | | | | | 0.14 | 1.19 | | |
| - | AAATATCTACACAGCCATCTGTATGAACATGG | | | | | | | 0.17 | 1.17 | | |
| - | AAACAGATAGCGGTGTAGATATTTCCATGTTC | | | | | | | 0.12 | 1.21 | | |

Degenerate probe:
ATCTGTATAAACATGGAAATATCTACACCGCY

**Table S7. Long-read sequencing data used in this study.**

| | PacBio | | Nanopore | |
|---|---|---|---|---|
| **species** | **accession** | **reference** | **accession** | **reference** |
| **Human** | SRR2097942 | (Kronenberg et al. 2018) | PRJNA505331 | generated for this study |
| **Chimpanzee** | SRR5269473 | (Kronenberg et al. 2018) | PRJNA505331 | generated for this study |
| **Bonobo** | NA | NA | PRJNA505331 | generated for this study |
| **Gorilla** | ERR1294100 | (Gordon et al. 2016) | PRJNA505331 | generated for this study |
| **Sumatran orangutan** | SRR5235143 | (Kronenberg et al. 2018) | PRJNA505331 | generated for this study |
| **Bornean orangutan** | NA | NA | PRJNA505331 | generated for this study |

**Table S8. The Nanopore run statistics.** The sequencing run was performed in-house.

| Species | # reads | Largest read | Total length | GC (%) | N50 |
|---|---|---|---|---|---|
| **Human** | 22,792 | 158,560 | 369,006,091 | 40.87 | 30,969 |
| **Chimpanzee** | 3,720 | 152,385 | 51,497,917 | 40.4 | 26,503 |
| **Bonobo** | 38,896 | 138,808 | 626,666,773 | 40.66 | 25,559 |
| **Gorilla** | 31,920 | 205,659 | 491,213,258 | 40.7 | 28,398 |
| **Sumatran orangutan** | 36,024 | 165,866 | 574,554,722 | 41.06 | 25,765 |
| **Bornean orangutan** | 18,942 | 168,772 | 357,490,403 | 40.89 | 36,839 |
| **unclassified** | 20,243 | 149,989 | 233,922,964 | 41.59 | 24,026 |

| Species | barcode | # reads(>= 0 bp) | # reads (>= 10000 bp) | # reads (>= 25000 bp) | # reads (>= 50000 bp) |
|---|---|---|---|---|---|
| **Human** | 5 | 22,975 | 11,213 | 5,292 | 1,273 |
| **Chimpanzee*** | 3 | 3,767 | 1,636 | 662 | 133 |
| **Bonobo** | 6 | 39,094 | 21,997 | 8,602 | 1,099 |
| **Gorilla** | 2 | 32,147 | 15,797 | 6,944 | 1,297 |
| **Sumatran orangutan** | 4 | 36,188 | 19,464 | 7,494 | 1,376 |
| **Bornean orangutan** | 1 | 19,053 | 9,908 | 5,410 | 1,665 |
| **unclassified** | NA | 20,800 | 7,504 | 2,817 | 548 |

| Species | Total length (>= 0 bp) | Total length (>= 10000 bp) | Total length (>= 25000 bp) | Total length (>= 50000 bp) |
|---|---|---|---|---|
| **Human** | 369,082,543 | 320,941,552 | 223,775,004 | 82,372,883 |
| **Chimpanzee** | 51,518,473 | 43,074,664 | 27,084,394 | 8,685,972 |
| **Bonobo** | 626,750,977 | 542,823,356 | 322,027,811 | 66,560,937 |
| **Gorilla** | 491,309,945 | 424,516,824 | 278,640,860 | 82,259,580 |
| **Sumatran orangutan** | 574,626,365 | 491,328,784 | 296,629,892 | 86,890,275 |
| **Bornean orangutan** | 357,538,219 | 319,743,182 | 244,941,230 | 111,281,736 |
| **unclassified** | 234,155,959 | 187,541,455 | 112,730,956 | 35,327,841 |

*The lower than expected yield for the chimpanzee is a consequence of a manufacturing problems with Oxford Nanopore barcode NB03 that did not pass the quality control (Marta Tomaszkiewicz, personal communication).

**Table S9. The PacBio run statistics (from public data). One SMRT cell for each species was obtained from the publicly available data presented in Table S7.**

| Species | # reads | Largest read | Total length | GC (%) | N50 |
|---|---|---|---|---|---|
| **Human** | 124,343 | 70,526 | 1,071,435,753 | 41.77 | 18,950 |
| **Chimpanzee** | 152,937 | 140,683 | 3,260,339,990 | 42.23 | 32,814 |
| **Bonobo** | NA | NA | NA | NA | NA |
| **Gorilla** | 153,383 | 183,711 | 3,601,234,294 | 40.84 | 34,065 |
| **Sumatran orangutan** | 151,588 | 131,289 | 2,235,949,129 | 41.2 | 30,556 |
| **Bornean orangutan** | NA | NA | NA | NA | NA |

| Species | # reads(>= 0 bp) | # reads (>= 10000 bp) | # reads (>= 25000 bp) | # reads (>= 50000 bp) |
|---|---|---|---|---|
| **Human** | 163,457 | 40,958 | 10,574 | 142 |
| **Chimpanzee** | 163,478 | 104,702 | 54,845 | 10,778 |
| **Bonobo** | NA | NA | NA | NA |
| **Gorilla** | 163,476 | 113,964 | 63,855 | 12,493 |
| **Sumatran orangutan** | 163,480 | 71,326 | 35,926 | 5,159 |
| **Bornean orangutan** | NA | NA | NA | NA |

| Species | Total length (>= 0 bp) | Total length (>= 10000 bp) | Total length (>= 25000 bp) | Total length (>= 50000 bp) |
|---|---|---|---|---|
| **Human** | 1,082,238,511 | 836,373,266 | 338,565,009 | 7,663,135 |
| **Chimpanzee** | 3,261,732,462 | 3,064,398,591 | 2,203,125,979 | 665,555,907 |
| **Bonobo** | NA | NA | NA | NA |
| **Gorilla** | 3,602,565,804 | 3,437,464,782 | 2,555,171,130 | 754,140,618 |
| **Sumatran orangutan** | 2,237,886,885 | 1,983,055,179 | 1,386,626,874 | 304,119,898 |
| **Bornean orangutan** | NA | NA | NA | NA |

**Table S10. The location of repeat arrays inside long (A) Nanopore and (B) PacBio sequencing reads.**

Three possible locations were called for the 39 repeated motifs and six species. The repeat arrays fully encompassed within long reads, but with some flanks on both sides, were considered 'nested'. The repeat arrays starting or ending within 30 bp of the read end, with flanking sequence only on one side, were called 'peripheral'. The repeat arrays fully covering the whole read, without flanks (with the tolerance of 30 bp from either end in order to account for the possibility of a sequencing error breaking the edge of an alignment), were called 'spanning'.

**A**

| NANOPORE | nested | peripheral (adjacent to read start or end) | spanning |
|---|---|---|---|
| **Human** | 94.63% | 4.97% | 0.40% |
| **Chimpanzee** | 93.36% | 6.40% | 0.25% |
| **Bonobo** | 95.03% | 4.51% | 0.46% |
| **Gorilla** | 93.29% | 5.61% | 1.10% |
| **S. orangutan** | 90.95% | 8.72% | 0.33% |
| **B. orangutan** | 90.26% | 9.44% | 0.31% |

**B**

| PACBIO | nested | peripheral (adjacent to read start or end) | spanning |
|---|---|---|---|
| **Human** | 99.02% | 0.98% | 0.00% |
| **Chimpanzee** | 99.39% | 0.61% | 0.00% |
| **Gorilla** | 99.13% | 0.87% | 0.00% |
| **S. orangutan** | 99.53% | 0.47% | 0.00% |

**Table S11.** Median lengths of repeat arrays in Nanopore and PacBio reads. NA means that a given motif was not found in the reads, whereas dash means long reads were not available. The background color corresponds to the repeat array length from the shortest (white) to the longest (dark red).

| median length of a repeat array [bp] | NANOPORE | PACBIO |
|---|---|---|
| | 76-7,344 | 76-817 |

| | | Human | | Chimpanzee | | Bonobo | | Gorilla | | Sumatran Orangutan | | Bornean Orangutan | |
|---|---|---|---|---|---|---|---|---|---|---|---|---|---|
| index | longest repeat array of a given motif found [bp] | NANOPORE | PACBIO | NANOPORE | PACBIO | NANOPORE | PACBIO | NANOPORE | PACBIO | NANOPORE | PACBIO | NANOPORE | PACBIO |
| 1 | AATGG | 32,100 | 724 | 6,280 | 2,023 | 44,804 | - | 59,411 | 6,560 | 20,150 | 548 | 32,368 | - |
| 2 | ACTCC | NA | 193 | NA | 156 | 76 | - | NA | 124 | NA | 268 | NA | - |
| 3 | AAAG | 340 | 840 | 323 | 274 | 448 | - | 562 | 584 | 488 | 331 | 464 | - |
| 4 | AATGGAGTGG | 220 | 184 | NA | 165 | 114 | - | 104 | 169 | 230 | 311 | 255 | - |
| 5 | AATGGAATGGAGTGG | 128 | 166 | NA | 196 | 632 | - | 175 | 466 | 434 | 312 | 4,360 | - |
| 6 | AATGGAGTGGAGTGG | NA | 155 | NA | 185 | NA | - | 174 | 196 | NA | 265 | 162 | - |
| 7 | AAGTGGAATGG | 77 | 140 | 88 | 122 | 110 | - | NA | 254 | 196 | 425 | 106 | - |
| 8 | AATGGAATGTG | 76 | 86 | NA | 79 | NA | - | NA | 153 | 121 | 306 | NA | - |
| 9 | AAAGTGGAATGG | 84 | NA | NA | NA | NA | - | 121 | 177 | 118 | 370 | 120 | - |
| 10 | AACGTGGAATGG | NA | NA | NA | NA | NA | - | NA | 84 | 179 | 283 | 109 | - |
| 11 | AATGGAATGTGG | 253 | 107 | NA | 85 | 132 | - | 137 | 165 | 378 | 1,857 | 368 | - |
| 12 | AATGGAATGGAATGTGG | 153 | 118 | NA | 109 | 98 | - | 119 | 232 | 165 | 390 | 276 | - |
| 13 | AATGGAATGGAATGGAATGTGG | 393 | 213 | NA | 129 | 110 | - | 191 | 424 | 428 | 987 | 370 | - |
| 14 | AAATGGACTCGAATGGAATCATC | NA | 258 | NA | 116 | NA | - | 159 | 76 | NA | NA | NA | - |
| 15 | AATCGAATGGAATGG | 173 | 209 | NA | 149 | 104 | - | NA | 314 | NA | 84 | NA | - |
| 16 | AAATGGAATCGAATGGAATCATC | 165 | 214 | NA | 183 | 109 | - | 201 | 93 | NA | NA | NA | - |
| 17 | AATCATCGAATGGAATCGAATGG | 504 | 387 | NA | 291 | 205 | - | 505 | 171 | NA | NA | NA | - |
| 18 | AATCATCGAATGGACTCGAATGG | 189 | 213 | NA | NA | 186 | - | 128 | NA | NA | NA | NA | - |
| 19 | AATCATCATGAATGGAATCGAATGG | 520 | 284 | 471 | 340 | 749 | - | 535 | 129 | NA | NA | NA | - |
| 20 | AAATGGAATCGAATGGAATCATCATC | 606 | 385 | 490 | 220 | 432 | - | 461 | 204 | NA | NA | NA | - |
| 21 | AAATGGAATCGAATGTAATCATCATC | NA | NA | NA | NA | NA | - | NA | NA | NA | NA | NA | - |
| 22 | AATCATCATCGAATGGAATCGAATGG | 6,546 | 1,398 | 531 | 428 | 6,524 | - | 4,201 | 760 | NA | NA | NA | - |
| 23 | AAACATGTAAATATTTACACAGAATCTGTAT | NA | NA | 21,231 | 15,587 | 37,584 | - | 7,755 | NA | NA | NA | NA | - |
| 24 | AAATATCTACACCGCTATCTGTATGAACATGG | NA | NA | 93 | 175 | 757 | - | 37,846 | 17,583 | NA | NA | 12,003 | - |
| 25 | AAATATCTACACCGCCATCTGTATGAACATGG | NA | NA | NA | 163 | 684 | - | 17,586 | 4,336 | NA | NA | 402 | - |
| 26 | AAACATGGAAATATCTACACAGCCATCTGTAT | NA | NA | 856 | 1,139 | 3,076 | - | 591 | 477 | NA | NA | 106 | - |
| 27 | AAACATGGAAATATCTACACCACCATCTGTAT | NA | NA | 533 | 721 | 905 | - | 741 | 318 | NA | NA | 116 | - |
| 28 | AAACATGGAAATATCTACACCACTATCTGTAT | NA | NA | 475 | 691 | 1,189 | - | 486 | 615 | NA | NA | 110 | - |
| 29 | AAACATGGAAATATCTACACCGCCATCTGTAT | NA | NA | 9,806 | 14,262 | 22,518 | - | 13,391 | 8,151 | 3,070 | NA | 308 | - |
| 30 | AAACATGGAAATATCTACACCGCTATCTGTAT | NA | NA | 5,446 | 6,400 | 26,007 | - | 15,107 | 16,832 | 441 | NA | 3,179 | - |

| | | | | | | | | | | | | | |
|---|---|---|---|---|---|---|---|---|---|---|---|---|---|
| 31 | AAACATGGAAATATCTACACCGCTATCTGTGT | NA | NA | 558 | 921 | 861 | - | 551 | 3,632 | NA | NA | 163 | - |
| 32 | AAACATGGAAATATCTACACTGCCATCTGTAT | NA | NA | 792 | 636 | 1,567 | - | 563 | 368 | NA | NA | 308 | - |
| 33 | AAACATGGAAATATCTACACTGCTATCTGTAT | NA | NA | 724 | 622 | 1,087 | - | 607 | 391 | NA | NA | NA | - |
| 34 | AAACATGGAAATATCTACACCGCGATCTGTAT | NA | NA | 461 | 776 | 476 | - | 119 | 223 | NA | NA | NA | - |
| 35 | AAACATGGAAATATCTACACCGCCATCTCTAT | NA | NA | 969 | 1,367 | 1,941 | - | 2,414 | 3,643 | NA | NA | 95 | - |
| 36 | AAACATGGAAATATCTACACCGCTATCTCTAT | NA | NA | 1,472 | 4,063 | 3,032 | - | 7,401 | 3,340 | NA | NA | 99 | - |
| 37 | AAATATCTACACAGCTATCTGTATGAACATGG | NA | NA | NA | NA | 102 | - | 2,298 | 928 | NA | NA | NA | - |
| 38 | AAATATCTACACCACTATCTGTATGAACATGG | NA | NA | NA | NA | 101 | - | 511 | 472 | NA | NA | NA | - |
| 39 | AAATATCTACACCGCTATCTGCATGAACATGG | NA | NA | NA | NA | NA | - | 345 | 874 | NA | NA | NA | - |

**Table S12. Calculating the number of arrays annotated with multiple repeated motifs.**

| NANOPORE | Arrays annotated with 1 motif | with 2 motifs | with 3 motifs | with >3 motifs | % of arrays annotated with multiple repeats |
|---|---|---|---|---|---|
| human | 2,221 | 267 | 54 | 3 | 12.73% |
| chimpanzee | 615 | 66 | 19 | 1 | 12.27% |
| bonobo | 9,968 | 1,363 | 236 | 74 | 14.37% |
| gorilla | 9,804 | 2,251 | 730 | 343 | 25.32% |
| Sumatran orangutan | 964 | 68 | 27 | 8 | 9.65% |
| Bornean orangutan | 762 | 71 | 17 | 5 | 10.88% |

**Table S13. Inter-generational change in the (AATGG)n repeat density**

| Parent | Parent density [kb/Mb] | Child | Child density [kb/Mb] | Family | Fold change* |
|---|---|---|---|---|---|
| father | 4.22 | son | 4.21 | 77 | 1.001 |
| mother | 3.70 | daughter | 3.71 | 78 | 1.003 |
| father | 6.10 | son | 5.76 | HG | 1.059 |
| mother | 5.17 | son | 5.76 | HG | 1.115 |
| father | 4.21 | daughter | 3.71 | 78 | 1.137 |
| mother | 3.36 | son | 4.21 | 77 | 1.252 |

*The ratio of Child:Parent or Parent:Child, whichever is greater than one

**Table S14. The average difference in density in male-biased repeats in males vs. females for each species**

| Species | Gorilla | Chimpanzee | Bonobo | Human* |
|---|---|---|---|---|
| Difference in kb/Mb | 8.08 | 1.52 | 3.20 | 2.11 |

*In human, the difference was calculated using 9 HGDP males and the parents from trios
